# A Systematic Survey of Reversibly Covalent Dipeptidyl Inhibitors of the SARS-CoV-2 Main Protease

**DOI:** 10.1101/2023.01.17.524469

**Authors:** Zhi Zachary Geng, Sandeep Atla, Namir Shaabani, Veerabhadra R. Vulupala, Kai S. Yang, Yugendar R. Alugubelli, Kaustav Khatua, Peng-Hsun Chase Chen, Jing Xiao, Lauren R. Blankenship, Xinyu R. Ma, Erol C. Vatansever, Chia-Chuan Cho, Yuying Ma, Robert Allen, Henry Ji, Shiqing Xu, Wenshe Ray Liu

## Abstract

SARS-CoV-2 is the coronavirus pathogen of the currently prevailing COVID-19 pandemic. It relies on its main protease (M^Pro^) for replication and pathogenesis. M^Pro^ is a demonstrated target for the development of antivirals for SARS-CoV-2. Past studies have systematically explored tripeptidyl inhibitors such as nirmatrelvir as M^Pro^ inhibitors. However, dipeptidyl inhibitors especially those with a spiro residue at their P2 position have not been systematically investigated. In this work, we synthesized about 30 reversibly covalent dipeptidyl M^Pro^ inhibitors and characterized them on *in vitro* enzymatic inhibition potency, structures of their complexes with M^Pro^, cellular M^Pro^ inhibition potency, antiviral potency, cytotoxicity, and *in vitro* metabolic stability. Our results indicated that M^Pro^ has a flexible S2 pocket that accommodates dipeptidyl inhibitors with a large P2 residue and revealed that dipeptidyl inhibitors with a large P2 spiro residue such as (*S*)-2-azaspiro[4,4]nonane-3-carboxylate and (*S*)-2-azaspiro[4,5]decane-3-carboxylate have optimal characteristics. One compound MPI60 containing a P2 (*S*)-2-azaspiro[4,4]nonane-3-carboxylate displayed high antiviral potency, low cellular cytotoxicity, and high *in vitro* metabolic stability and can be potentially advanced to further preclinical tests.

## Introduction

Coronaviruses (CoVs) are RNA pathogens that infect vertebrates including humans. Although mildly pathogenic human CoVs were discovered in the 1960s, the first pandemic human CoV, severe acute respiratory syndrome (SARS)-CoV, was yet to emerge until 2003. Since then, within 20 years two more pandemic human CoVs, Middle East respiratory syndrome (MERS)-CoV and SARS-CoV-2, appeared with the later one creating a wreak havoc across the globe. All three pandemic CoVs were believed originated from animals and spread to humans during close human-animal interactions. The high outbreak frequency of CoV pandemics in the past two decades and the ever-increasing close human-animal interactions in the modern society combinedly portend the future pandemic CoV outbreaks. With the current coronavirus disease 2019 (COVID-19) pandemic prevailing and future CoV pandemics looming, it is paramount to develop orally available small-molecule drugs that can be broadly used as CoV antivirals for both treatment and prevention. So far, three orally available medications including remdesivir, molnupiravir, and PAXLOVID^TM^ have been approved for emergency use of treating COVID-19 patients. Both remdesivir and molnupiravir are nucleotide analogues. Remdesivir is an RNA replication inhibitor and known to have low efficacy in inhibiting SARS-CoV-2.^1^ On the contrary, molnupiravir is an RNA mutagen. Clinical tests showed that molnupiravir reduced the risk of hospitalization and death by 50% compared to placebo for patients with mild and moderate COVID-19. However, its mutagen nature that drives SARS-CoV-2 to undergo mutagenesis warrants use with caution. Unlike remdesivir and molnupiravir, PAXLOVID^TM^ is a combination therapy of nirmatrelvir and ritonavir. Nirmatrelvir is a reversibly covalent inhibitor of the SARS-CoV-2 main protease (M^Pro^) that serves an essential role in the viral pathogenesis and replication. Ritonavir is a human cytochrome P450 3A4 inhibitor that improves the metabolic stability of nirmatrelvir. Potential toxicity of PAXLOVID^TM^ requires its stop of use after 5 days and it has failed as a preventative for COVID-19 in clinical tests. The current published results have shown that nirmatrelvir is a substrate of P-glycoprotein multidrug transporter (P-pg) that continuously pumps various and structurally unrelated compounds to the outside of human cells. P-gp is known with varied expression levels in different tissues. Although ritonavir is a P-gp inhibitor as well, the expression variation of P-gp in different tissues likely causes different inhibition efficacy of PAXLOVID^TM^ in different tissues. This may explain why many patients had COVID-19 rebound after stopping taking PAXLOVID^TM^ and SARS-CoV-2 from these patients with COVID rebound did not show resistance to PAXLOVID^TM^. Due to concerns related to existing small molecule SARS-CoV-2 antivirals, the research of developing SARS-CoV-2 antivirals that have characteristics better than existing antivirals is still urgent.

M^Pro^ is a cysteine protease that uses four binding pockets S1, S2, S4, and S3’ in the active site to engage P1, P2, P4 and P3’ residues in a protein substrate for binding (Figure 1A). Nirmatrelvir can be classified as a tripeptidyl inhibitor that uses its P1 and P2 residues and *N*-terminal trifluoroacetyl group to bind S1, S2, and S4 pockets, respectively, in M^Pro^ and an activated nitrile warhead to covalently engage C145, the catalytic cysteine of M^Pro^ (Figure 1B).^2^ Similar to that in a protein substrate, the P3 side chain of nirmatrelvir doesn’t directly interact with M^Pro^. Since P3 is not necessary for an inhibitor to engage M^Pro^ for binding, multiple potent dipeptidyl inhibitors that use their P1 and P2 residues and *N*-terminal group to bind S1, S2, and S4 pockets in M^Pro^ and a covalent warhead to engage C145 of M^Pro^ have also been reported. Representative dipeptidyl M^Pro^ inhibitors include GC376, 11a, and PF-00835231 (Figure 1B). However, a systematic study of dipeptidyl M^Pro^ inhibitors on how different chemical identities in P1 and P2 residues, *N*-terminal groups, and warheads influence M^Pro^ inhibition, structural aspects in binding M^Pro^, cellular and antiviral potency, and metabolic stability has not been reported. In this work, we wish to report a systematic survey of dipeptidyl M^Pro^ inhibitors of M^Pro^ and their potential use as SARS-CoV-2 antivirals.

**Figure 1:**
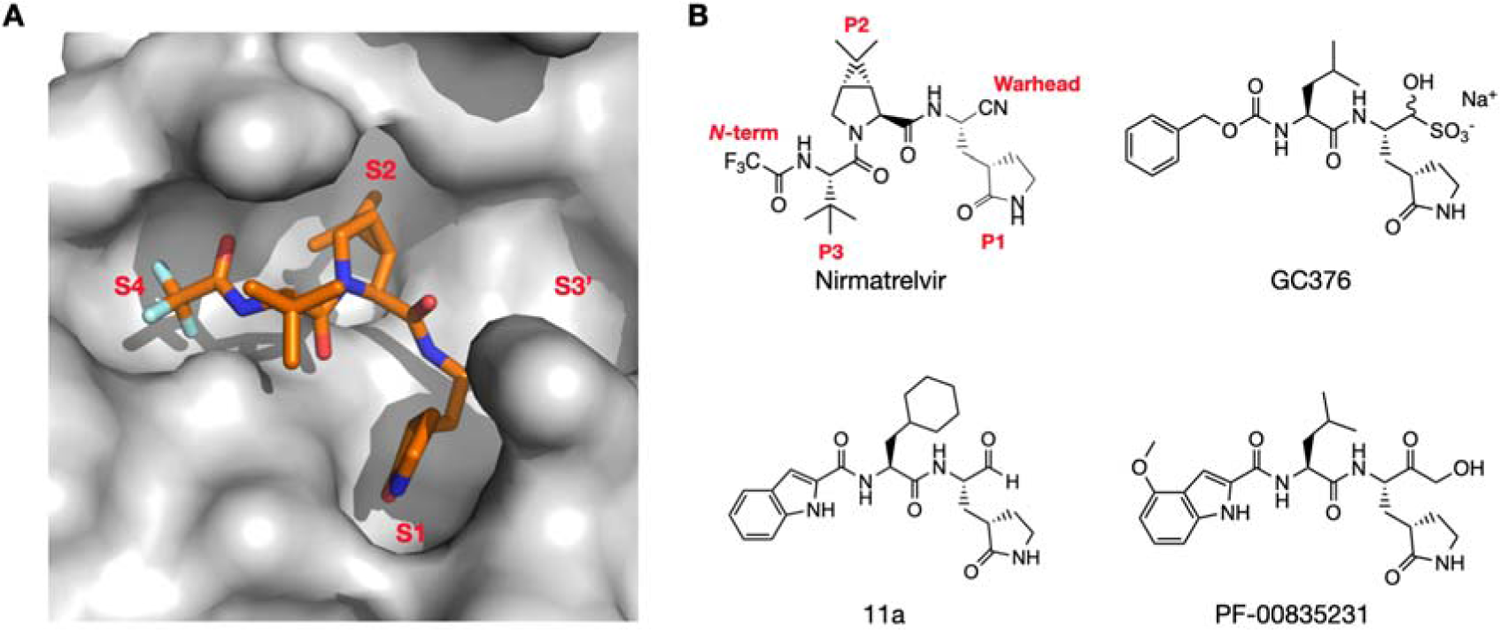
(**A**) The M^Pro^-nirmatrelvir complex. The structure is based on the pdb entry 7TE0.^2^ The contoured surface of M^Pro^ is shown. Four substrate binding pockets in M^Pro^ are labeled. (**B**) The structures of nirmatrelvir, GC376, 11a, and PF-00835231. Chemical positions in nirmatrelvir are labeled.

## Results and Discussion

### The Design and Synthesis of Dipeptidyl M^Pro^ Inhibitors

We followed two general designs shown in Figure 2 for the design and synthesis of dipeptidyl M^Pro^ inhibitors. Group A compounds were developed during the early phase of the pandemic. They were primary amino acid-based and contained a 3-methylpyrrolidin-2-one side chain at the P1 site due to the demonstrated high affinity of this side chain toward the S1 pocket of M^Pro^.^3^ Isopropyl (**h**), benzyl (**i**), *t*-butyl (**j**), and cyclohexylmethyl (**k**) as a side chain at the P2 site were previously tested in tripeptidyl M^Pro^ inhibitors and were included in the Group A compounds.^4^ Both P1 and P2 residues are in the L configuration. Our previously works showed that *O-t-*butyl-L-threonine (**l**) as the P3 residue in tripeptidyl M^Pro^ inhibitors led to high cellular and antiviral potency.^4, 5^ We included this residue at the P2 site as well hoping to observe a similar effect. The *N*-terminal groups were chosen between carboxybenzyl (CBZ, **a**), 3-chloro-CBZ (**b**), 3-acetoxy-CBZ (**c**), 4-chloro-2-fluorocinnamoyl (**d**), 1*H*-indole-2-carbonyl (**e**), 4-methoxyl-1*H*-indole-2-carbonyl (**f**), and trifluoroacetyl (**g**). Some of these groups were used in inhibitors for either SARS-CoV or SARS-CoV-2 inhibitors. The warhead was chosen between aldehyde (**m**) and nitrile (**n**) that reversibly react with C145 of M^Pro^ to form hemithioacetal and thioimidate, respectively. Group B compounds were developed later, and all contained a modified proline at the P2 site. For this group of dipeptidyl M^Pro^ inhibitors, the P1 side chain was primarily 3-methylpyrrolidin-2-one (**a4**) and one inhibitor had a 3-methylpiperidin-2-one (**a5**) side chain due to the demonstrated high potency of some inhibitors with this side chain. Proline-based P2 residues in Group B compounds included (*R*)-3-*t*-butyloxyl-L-proline (**v**), (*R*)-3-cyclohexyl-L-proline (**w**), (1*S*, 2*S*, 5*R*)-6,6-dimethyl-3-azabicyclo[3,1,0]hexane-2-carboxylate (**x**) that is the P2 residue in nirmatrelvir, (*S*)-5-azaspiro[2,4]heptane-6-carboxylate (**y**), (*S*)-6-azaspiro[3,4]octane-7-carboxylate (**z**), (*S*)-2-azaspiro[4,4]nonane-3-carboxylate (**a1**), (*S*)-2-azaspiro[4,5]decane-3-carboxylate (**a2**), and (1*S*, 2*S*, 5*R*)-3-azabicyclo[3,3,0]octane-2-carboxylate (**a3**). A survey of multiple M^Pro^-inhibitor complex structures revealed that the peptide region aa46-51 in M^Pro^ that caps the S2 pocket is highly flexible, and this structural flexibility allows even the flipping of C44 close to 180°C to form a Y-shaped, S-O-N-O-S-bridged crosslink with two other residues C22 and K61 in M^Pro^.^6^ This structural flexibility and the crosslink formation leave a much more open, large S2 pocket that accommodates potentially a large P2 residue in a peptidyl inhibitor. Proline-based P2 residues in Group B compounds were designed for this reason to test how deep and bulky the S2 pocket can turn to be. **V** and **W** are 3-substituted prolines, **x** and **a4** are bicyclic compounds, and **y**-**a2** are spiro compounds. They were selected for readily synthetic accessibility. The *N*-terminal groups for Group B compounds were more diverse than that in Group A. Besides several moieties used in Group A, other *N*-terminal groups included *t*-butyloxycarbonyl (Boc, **o**), 4-trifluoromethoxyphenoxycarbonyl (**p**), 2,4-dichlorophenoxycarbonyl (**q**), 3,4-dichlorophenoxycarbonyl (**r**), 4-chlorophenylcarbamoyl (**s**), 3-cyclohexylpropanoyl (**f**), and 2-cyclohexyloxyacetyl (**u**). Some of these *N*-terminal groups were previously used in dipeptidyl M^Pro^ inhibitors. Others were designed to explore different interactions with the S4 pocket of M^Pro^. The warhead was chosen between aldehyde (**m**) and nitrile (**n**) as well.

**Figure 2:**
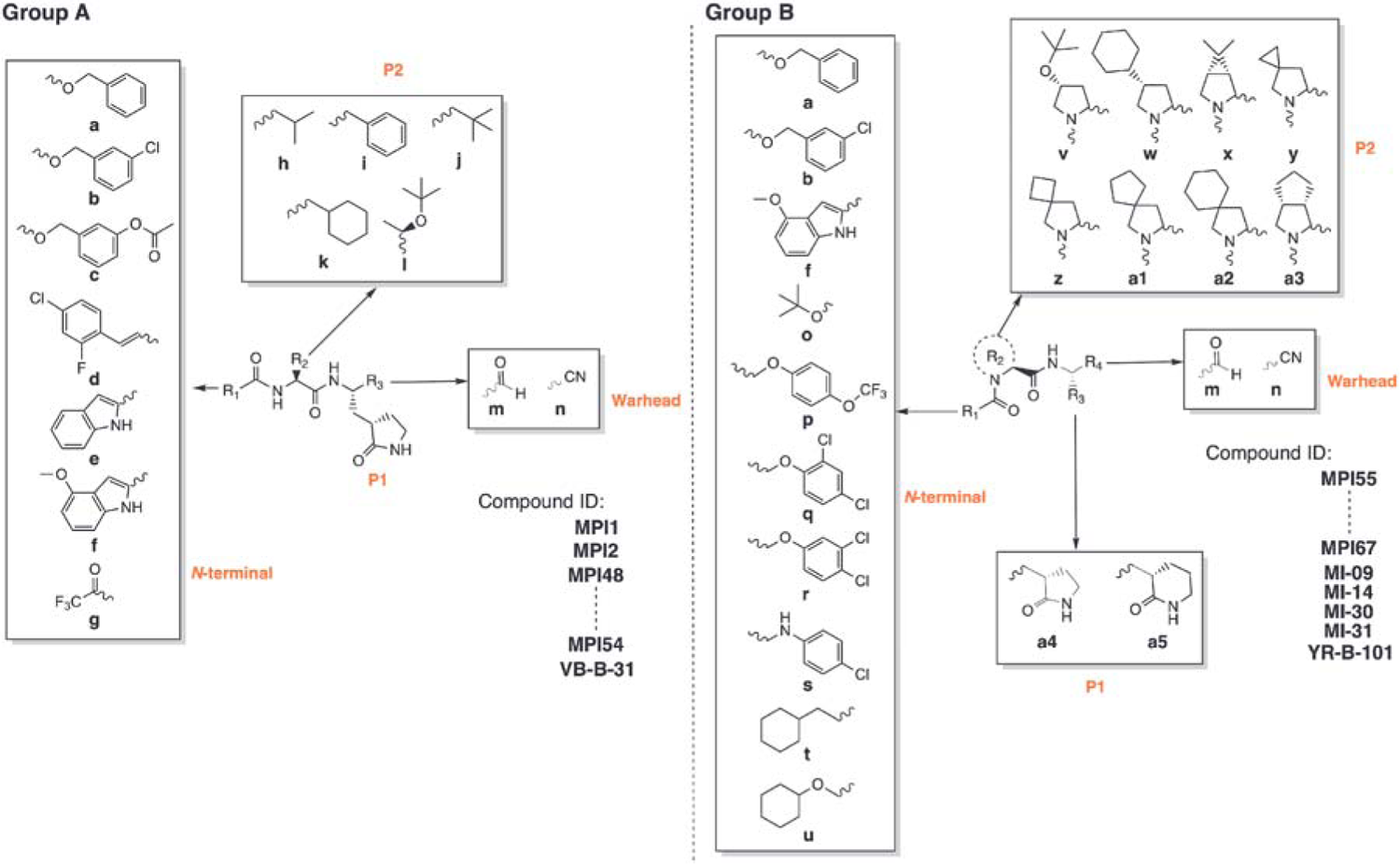
A diagram showing all dipeptidyl compounds that have been synthesized.

We followed two synthetic routes shown in figure 8 for the synthesis of aldehyde and nitrile-based dipeptidyl M^Pro^ inhibitors. In total, 29 dipeptidyl M^Pro^ inhibitors were synthesized including MI-09, MI-14, MI-30, and MI-31, four compounds that were previously developed by a different lab and included as comparison.^1^ MPI1, MPI2, and GC376 are three dipeptidyl M^Pro^ inhibitors that were previously characterized. They are included as comparison as well. All inhibitors have their compositions shown in Table 1 and their chemical structures are presented in Supplementary Figure 1 as well.

**Table 1:** M^Pro^ inhibitors, their enzymatic IC_50_, cellular EC_50_, antiviral EC_50_, CC_50_, and CL_int_ values

| ID | R1 | R2 | R3 | R4 | Enzymatic<br>IC<sub>50</sub> (μM) | Cellular<br>EC<sub>50</sub> (μM) | Antiviral<br>EC<sub>50</sub> (μM) | CC<sub>50</sub><br>(μM) | CL<sub>int</sub> (μL<br>/min/kg) | PDB<br>Entry |
| --- | --- | --- | --- | --- | --- | --- | --- | --- | --- | --- |
| **Group A** |  |  |  |  |  |  |  |  |  |  |
| *MPI1<sup>a</sup>* | *a* | *i* | *m* |  | *0.100* | *> 10* |  |  |  | *7JPZ* |
| *MPI2<sup>a</sup>* | *d* | *i* | *m* |  | *0.103* | *> 2* |  |  |  |  |
| *GC376<sup>a</sup>* | *a* | *h* | *m* |  | *0.030* | *> 2* |  |  |  | *7C6U* |
| *11a* | *e* | *k* | *m* |  | *0.053* | *0.66* |  |  |  | *6LZE* |
| MPI48 | e | j | m |  | 0.029 | 1.6 |  |  |  | 7SD9 |
| MPI49 | f | j | m |  | 0.074 | 0.91 | 1.3 |  |  | 7SDA |
| MPI50 | a | j | m |  | 0.053 | 0.73 | 0.75 |  |  |  |
| MPI51 | a | k | m |  | 0.056 | 0.77 | 6.7 |  |  |  |
| MPI52 | b | k | m |  | 0.11 | 0.59 | 2.6 |  |  |  |
| MPI53 | c | k | m |  | 0.098 | 3.50 |  |  |  |  |
| MPI54 | a | l | m |  | 0.40 | > 10 |  |  |  |  |
| VB-B-31 | g | k | n |  | > 2 |  |  |  |  |  |
| **Group B** |  |  |  |  |  |  |  |  |  |  |
| MPI55 | a | v | a4 | m | 0.56 |  |  |  |  |  |
| MPI56 | a | w | a4 | m | 0.51 |  |  |  |  |  |
| MPI57 | a | x | a4 | m | 0.025 | 1.1 | 1.5 |  |  |  |
| MPI58 | a | y | a4 | m | 0.025 | 0.66 | 3.6 |  |  |  |
| MPI59 | a | z | a4 | m | 0.030 | 0.43 | 0.56 |  |  |  |
| MPI60 | a | a1 | a4 | m | 0.022 | 0.088 | 0.37 | 95 | 11 |  |
| MPI61 | a | a2 | a4 | m | 0.049 | 0.085 | 0.37 | 230 | 45 |  |
| MI-09<sup>d</sup> | p | x | a4 | m | 0.055 | 0.27 |  |  |  | 7SDC |
| MI-14<sup>d</sup> | q | x | a4 | m | 0.028 | 0.28 |  |  |  |  |
| MI-30<sup>d</sup> | q | a3 | a4 | m | 0.040 | 0.65 |  |  |  |  |
| MI-31<sup>d</sup> | r | a3 | a4 | m | 0.042 | 2.7 |  |  |  |  |
| YR-B-101 | o | z | a4 | m | > 5 |  |  |  |  |  |
| MPI62 | s | x | a4 | m | 0.99 |  |  |  |  |  |
| MPI63 | q | y | a4 | m | 0.039 | 2.9 |  |  |  |  |
| MPI64 | u | x | a4 | m | 0.025 | 3.6 |  |  |  |  |
| MPI65 | t | x | a4 | m | 0.041 | 0.56 | 0.96 |  |  |  |
| MPI66-1 | a | a1 | a4 | n | 2.2 |  |  |  |  |  |
| MPI66-2 | b | a1 | a4 | n | 3.6 |  |  |  |  |  |
| MPI66-3 | f | a1 | a4 | m | 1.5 |  |  |  |  |  |
| MPI66-4 | f | a1 | a5 | n | 6.0 |  |  |  |  |  |
| MPI67 | a | a2 | a4 | n | 1.4 |  |  |  |  |  |

<sup>a</sup>Data were taken from Cao *et al.*
<sup>b</sup>Recharacterized compounds from Qiao *et al.*

### The Enzymatic Inhibition Potency of Dipeptidyl M^Pro^ Inhibitors

We followed a previously established protocol that uses Sub3 (Dabcyl-KTSAVLQSGFRKME-Edans), a fluorogenic peptide substrate of M^Pro^ to determine IC_50_ values for all synthesized compounds.^7^ In this assay, we incubated M^Pro^ with a compound with varied concentrations for 30 min before Sub3 was added and the fluorescent product formation (Ex: 336 nm/Em: 455 nm) was recorded and analyzed to determine IC_50_. 30 min incubation time is a standard procedure that has been used by multiple labs in the determination of IC_50_ values for M^Pro^ inhibitors.^7–10^ Since all synthesized compounds are reversible covalent inhibitors, their incubation times with M^Pro^ are not expected to significantly influence their determined IC_50_ values. A previous test of a reversible covalent inhibitor in three different incubation times, 15, 30, and 60 min, led to very similar determined IC_50_ values.^4^ Determined IC_50_ values for all compounds are presented in Table 1. For Group A compounds, except MPI54 and VB-B-31, all other compounds have an IC_50_ value around or below 100 nM, comparable to that for MPI1, MPI2, and GC376.^3, 11^ MPI54 has *O-t-*butyl-threonine at the P2 site. The S2 pocket of M^Pro^ is known to prefer leucine, phenylalanine, and their analogues at the P2 site of substrates and inhibitors. It was not surprising that the installation of *O-t-*butyl-threonine at the P2 site in MPI54 led to weaker binding. However, it was intriguing to observe that *O-t-*butyl-threonine that is structurally very different from leucine and phenylalanine at the P2 site in MPI54 did not significantly distort the binding to M^Pro^ compared to MPI1. This observation indicates that the high structural plasticity of the S2 pocket can potentially accommodate a large variety of structurally unique and bulky P2 residues and inspired us to design compounds in Group B.^6^ VB-B-31 has a small *N*-terminal group and nitrile warhead. Since the nitrile warhead is demonstrated to engage the catalytic cysteine very efficiently, the low enzymatic inhibition potency of BV-B-31 is likely due to the relatively small trifluoroacetyl *N*-terminal group not being able to engage the S4 pocket. MPI1, GC376, MPI50, and MPI51 are structurally different only at the P2 site with a leucine, phenylalanine, or analogue. They have similar IC_50_ values indicating the M^Pro^ S2 pocket has similar binding preference toward leucine, phenylalanine, and their derivatives with a similar size. MPI48, MPI49, and MPI50 differ at the *N*-terminal group and have similar IC_50_ values. Compared to CBZ (**a**), 1*H*-indole-2-carbonyl (**e**) and 4-methoxyl-1*H*-indole-2-carbonyl (**f**) are more structurally rigid. Previous works have already shown that these three groups involve different interactions with M^Pro^. Their contributions to similar binding toward M^Pro^ are likely accidental. MPI51, MPI52, and MPI53 are structurally similar compounds. MPI52 and MPI53 have a 3-substitution on the *N*-terminal group. Both 3-chloro and 3-acetoxy groups in MPI52 and MPI53, respectively, did not significantly influence the binding to M^Pro^. Previous determined structures of M^Pro^ complexed with dipeptidyl inhibitors showed a loosely bound *N*-terminal CBZ.^3^ This relatively weak engagement of CBZ to M^Pro^ may explain relatively weak influence of a substitution in CBZ toward binding to M^Pro^. Although this information is not useful in the design of more potent inhibitors, it is helpful in the design of metabolically stable compounds since adding substitutions to CBZ can significantly change its metabolic stability.

Group B compounds were designed and synthesized based on information learnt from Group A compounds and other developments by exploring bicyclic side chains in the P2 residue of M^Pro^ inhibitors such as nirmatrelvir, MI-09, MI-14, and MPI29-MPI47. MPI55 and MPI56 are two inhibitors with a large 3-substitution at its P2 proline. The P2 (*R*)-3-cyclohexyl-L-proline (**w**) in MPI56 is also a highly rigid residue. Although both inhibitors showed relatively mild inhibition potency, compared to other inhibitors, with IC_50_ values a little higher than that for MPI54, their IC_50_ values around 0.5 hM still indicate that they can engage M^Pro^ for binding efficiently. 2 (*R*)-3-cyclohexyl-L-proline (**w**) is probably the largest P2 residue that has been tested so far. It is obvious that the S2 pocket of M^Pro^ can rearrange to accommodate large and bulky P2 residues in inhibitors.

Therefore, it is possible that other large P2 residues with strong M^Pro^ binding can be developed. This potential warrants further exploration. MPI57-MPI61 are structurally similar compounds with variation at the P2 site. They all have a *N*-terminal CBZ (**a**), P1 3-methylpyrrolidin-2-one side chain, and aldehyde warhead. Their P2 side chain varied among bicyclic and spiro moieties **x**-**a2**. All five compounds displayed very high M^Pro^ inhibition potency with an IC_50_ value below 50 nM. Four compounds MPI57-MPI60 have an IC_50_ value at or below 30 nM. And MPI60 has the lowest IC_50_ value as 22 nM among all compounds that were tested in this series and one of the most potent M^Pro^ inhibitors that have been developed so far. From the structural perspective, **w**-**a2** can be considered as leucine and phenylalanine analogues with higher structural rigidity by forming a basic proline ring. This high structural rigidity likely contributes to strong binding of these moieties to the M^Pro^ pocket. From MPI57 to MPI61, there is a significant increase of the size of P2 side chain. The surprisingly similar IC_50_ values determined for all five compounds approves the high structural plasticity of the M^Pro^ S2 pocket. MI-09, MI-14, MI-30, and MI-31 are four previously reported dipeptidyl M^Pro^ inhibitors. They were synthesized and recharacterized for comparison with all other dipeptidyl M^Pro^ inhibitors. All four molecules have a determined IC_50_ value around or below 50 nM. MI-09 and MI-14 have a P2 **x** residue but two different *N*-terminal groups **p** and **q**, respectively. These two compounds are structurally similar to MPI57 but with the *N*-terminal carbamate oxygen in MPI57 moved one position and additional substituent(s) added to the *N*-terminal phenyl group. Both MI-09 and MI-14 have a similar and slightly higher IC_50_ value than MPI57. This is likely due to the relatively loose bound *N*-terminal CBZ like group to M^Pro^. MI-30 and MI-31 have a P2 **a3** residue and **q** and **r**, respectively, as their *N*-terminal groups. Both have an IC_50_ value similar to other compounds that have a bicyclic or spiro residue at the P2 site. Given that the size of **a3** is similar to **x**, this is expected. A compound YR-B-101 that is structurally similar to MPI59 but with a *N*-terminal BOC (**o**) group was also made. This compound showed very weak enzymatic inhibition potency. Since compound 11a has a *N*-terminal indole that uses its indole imine to form a hydrogen bond with the backbone carbonyl oxygen of E66 in M^Pro^ that potentially contributes to the strong binding of 11a to M^Pro^, we tried to recapitulate this interaction by introducing **s** as a *N*-terminal group for MPI62 that is structurally similar to

MPI57, MI-09, and MI-14. Unfortunately, the determined IC_50_ value is drastically higher than that for the other three compounds. Two possibilities may contribute to this affinity decrease. The introduction of a hydrogen donor likely makes the molecule more favorable to be dissolved in water. The proposed hydrogen bond may not form as well. Since MI-14 that has an *N*-terminal **q** group has a lower IC_50_ value than MI-09, we grafted this moiety into MPI58 to afford MPI63. Compared to MPI58, MPI63 has a higher determined IC_50_ value. Based on all collected data so far, we could conclude that the CBZ (**a**) group is the best *N*-terminal group for dipeptidyl M^Pro^ inhibitors to achieve high potency. Since the *N*-terminal CBZ (**a**) group only loosely binds to M^Pro^, we thought that changing it to **t** and **u** that have a saturated cyclohexane might introduce better interactions with M^Pro^. Replacing the CBZ (**a**) group in MPI57 with **u** and **t** afforded MPI64 and MPI65, respectively. MPI64 has a determined IC_50_ value as same as MPI57. MPI65 has a slightly higher IC_50_ value. Although using **t** and **u** did not lead to more potent inhibitors, the results demonstrated that *N*-terminal groups other than the CBZ (**a**) group can lead to equal enzymatic inhibition potency. Based on all discussed compounds, optimal P2 residues are two primary amino acids **h** and **j** and all tested bicyclic and spiro amino acids **x**-**a3**. Since they led to very similar enzymatic inhibition potency, it is difficult to conclude about which one is the best although MPI60 that has a P2 **a1** residue has the lowest IC_50_ value among all tested compounds.

Previous works with tripeptidyl and dipeptidyl M^Pro^ inhibitors showed that replacing the aldehyde (**m**) warhead with nitrile (**n**) can still lead to potent inhibitors. To recapitulate this observation, we synthesized MPI66-1 that contained **n** and was structurally different from MPI60 only at the warhead and MPI66-2 that had an additional 3-chloro substituent on the *N*-terminal group. However, both compounds showed much lower potency than MPI60 and had an IC_50_ value above 2 hM. It is possible that **a2** at the P2 site introduces unique M^Pro^-inhibitor interactions that makes the covalent interaction between M^Pro^ C145 and nitrile (**n**) less favorable than that in the M^Pro^-nirmatrelvir complex. Replacing the *N*-terminal CBZ (**a**) group in MPI60 with **f** to afford MPI66-3 to recapitulate strong binding that was observed in the M^Pro^-11a complex also failed. MPI66-3 had a determined IC_50_ value above 1 hM, drastically higher than that for 11a. This observation corroborated the possible unique interactions with M^Pro^ induced by **a2** at the P2 site. A previous work showed that a 3-methylpiperidin-2-one (**a5**) side chain led to better enzymatic inhibition potency in a dipeptidyl inhibitor than its corresponding 3-methylpyrrolidin-2-one (**a4**) inhibitor. To rescue the potency of MPI66-1, we replaced its P1 **a4** residue with **a5** to afford MPI66-4. However, MPI66-4 exhibited ever lower inhibition potency with a determined IC_50_ value as 6.0 M. As discussed below, MPI61 that contains **a2** at its P2 site showed high cellular and antiviral potency. Based on this information, we synthesized MPI67, a nitrile (**b**)-containing MPI61 equivalent. However, this molecule showed low enzymatic inhibition potency with a determined IC_50_ value as 1.4 M. In our series of dipeptidyl M^Pro^ inhibitors, we concluded that the aldehyde (**m**) warhead is better than the nitrile (**n**) warhead.

### X-Ray Crystallography Analysis of M^Pro^ bound with MPI48, MPI49, and MI-09

We previously determined the crystal structure of M^Pro^ bound with MPI1. In this structure (pdb: 7JPZ), the *N*-terminal CBZ (**a**) group of MPI1 did not have defined electron density around it. Since the *N*-terminal group of MPI1 is similar to that in most Group A compounds and peptidyl M^Pro^ inhibitors with different primary amino acid residues at the P2 site were structurally well characterized before, we chose to conduct X-ray crystallography analysis of M^Pro^ bound with MPI48 and MPI49 in Group A since both have an *N*-terminal group not based on CBZ. We followed a previously established procedure to conduct the X-ray crystallography analysis of M^Pro^ bound with MPI48 and MPI49, respectively. We crystalized M^Pro^ in its apo form and then soaked obtained crystals with MPI48 or MPI49 before these crystals were mounted on an X-ray diffractometer for X-ray diffraction data collection. Collected data were then used to refine structures for M^Pro^ bound with MPI48 and MPI49, respectively. For M^Pro^-MPI48, the structure was determined with a resolution ofÅ. As shown in Figure 4A, the electron density at the active site of M^Pro^-MPI48 clearly showed the bound inhibitor and allowed the ambiguous refinement of all non-hydrogen atoms. The three methyl groups at the P2 *t*-butyl group were clearly observable. There was continuous electron density that connected the thiolate of M^Pro^ C145 with the P1 Ccr atom of MPI48 indicating a covalent bond formation. The electron density around the P1 Ccr of MPI48 allowed the refinement of a hemithioacetal hydroxyl group that had a strictly *S* confirmation and pointed exactly at the anion hole with a hydrogen bond distance to three backbone cr-amines from M^Pro^ residues G143, S144, and C145. This strict *S* confirmation of hemithioacetal hydroxide has been observed in M^Pro^ bound with other aldehyde-based inhibitors. The P1 side chain lactam used its amide oxygen to form a hydrogen bond with the H163 imidazole nitrogen and its amide nitrogen to form two hydrogen bonds with the E166 side chain carboxylate and the F140 backbone cr-amide oxygen. The P1 cr-amine engaged the H164 backbone cr-amide oxygen to form a hydrogen bond. The P2 *t*-butyl group fit neatly to the S2 pocket and was in close distance to side chains of H41, M165, and E189. M49 is a residue in the aa45-51 region that caps the S2 pocket. Its side chain was observed to fold into the S2 pocket in apo-M^Pro^ but typically flipped its position to open the S2 pocket to bind a peptidyl inhibitor. In determined M^Pro^-inhibitor complexes that were co-crystalized and had a closed active site due to protein packing in crystals, the M49 side chain was usually observed to cap the S2 pocket. However, our M^Pro^ crystals were obtained with an open active site allowing structural rearrangement around the active site and soaking them with peptidyl inhibitors always led to a flexible aa45-51 region whose structure could not be refined. In M^Pro^-MPI48, there was also no strong electron density at this region to allow refining its structure indicating a high structural flexibility of aa45-51. MPI48 has a *N*-terminal 1*H*-indole-2-carbonyl (**e**) group. It used its carbonyl oxygen and indole nitrogen to form a hydrogen bond with the E166 〈-amine and 〈-carbonyl oxygen, respectively.

**Figure 3:**
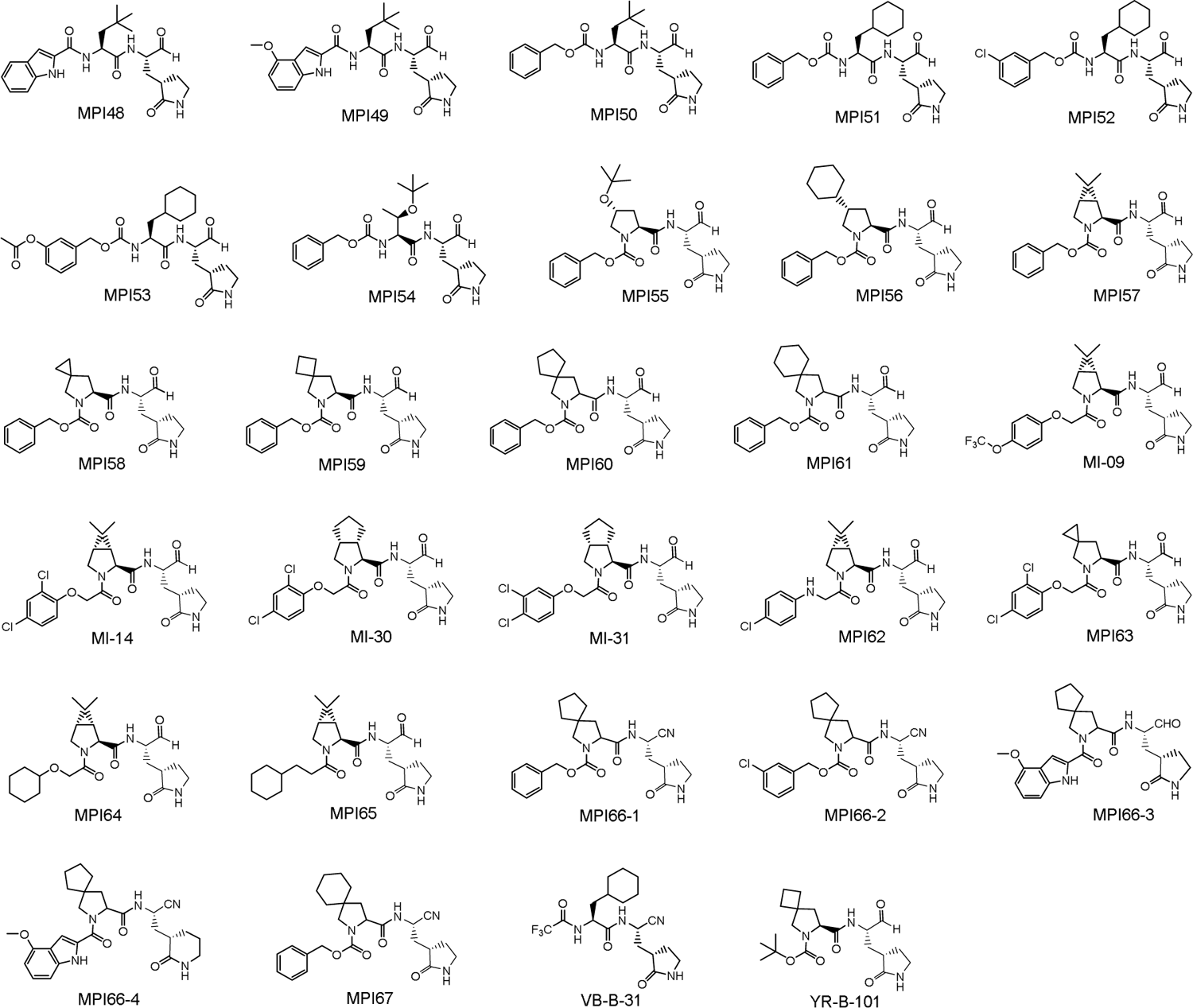
Structures of dipeptidyl M^Pro^ inhibitors.

The hydrogen bond between the MPI48 indole nitrogen and the E166 〈-carbonyl oxygen is unique to MPI48 and other dipeptidyl inhibitors with a *N*-terminal 1*H*-indole-2-carbonyl (**e**) group or analogue. A same hydrogen bond has been observed in M^Pro^ bound with other similar dipeptidyl inhibitors such as 11a as well. The M^Pro^-MPI49 complex structure was refined to a resolution ofÅ. As shown in Figure 4B, the active site electron density allowed the structural refinement of all chemical compositions of MPI49 except the *O*-methyl moiety of the *N*-terminal group. In the structure, there were not defined interactions with M^Pro^ that could make this *O*-methyl moiety to adopt a fixed confirmation. MPI49 involved interactions both covalent and other types with M^Pro^ that were mostly observed in the M^Pro^-MPI48 complex. One additional hydrogen bond that formed between the P2 〈-amine and a water molecule was observed in M^Pro^-MPI49. This water molecule was also within a hydrogen bond distance to the Q189 side chain amide.

**Figure 4.**
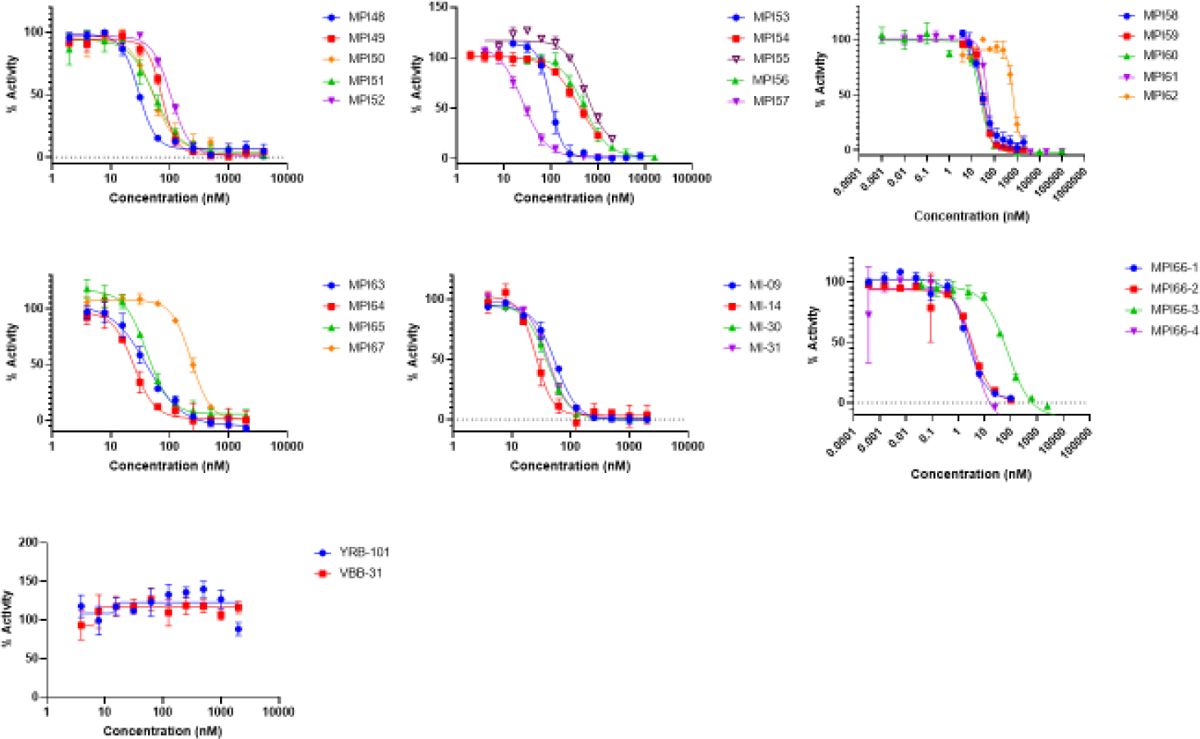
Inhibition curves of compounds on M. Triplicate experiments were performed for each compound. For all experiments, 20 or 10 nM M^Pro^ was incubated with an inhibitor for 30 min before 10 μ Sub3 was added. The M-catalyzed Sub3 hydrolysis rate was determined by measuring linear increase of product fluorescence (Ex: 336 nm/Em: 455 nm) for 5 min.

**Figure 5:**
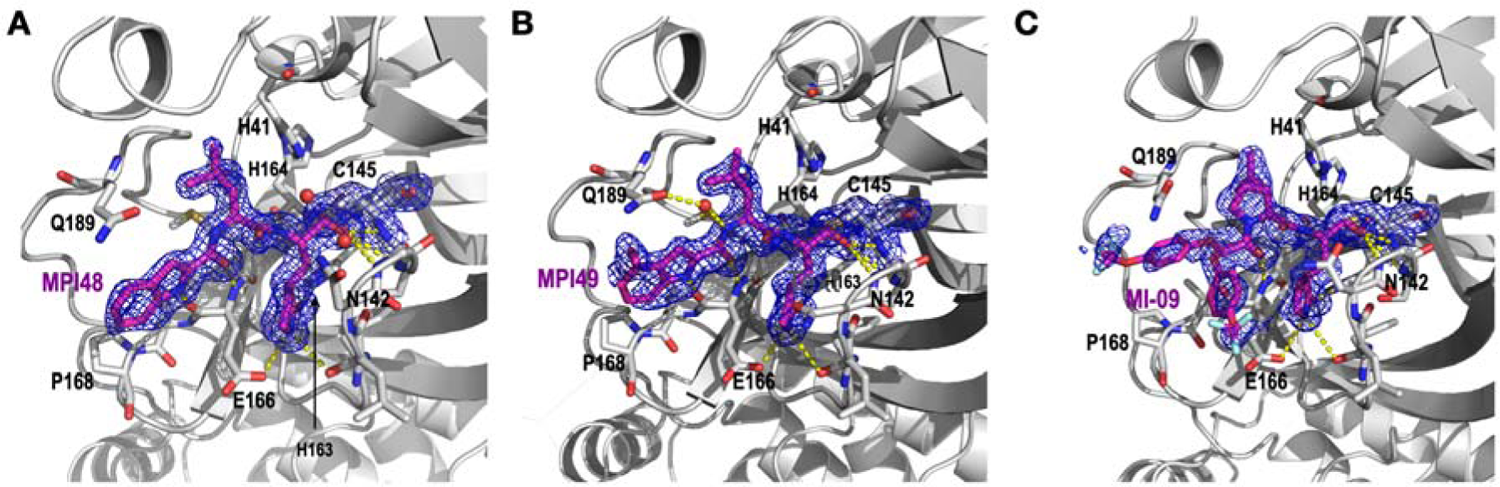
The crystal structures of (**A**) M^Pro^-MPI48, (**B**) M^Pro^-MPI49, and (**C**) M^Pro^-MI-09. The 2fo-fc maps around the inhibitor and C145 in all three structures were contoured at 1.

Most Group B compounds are structurally similar. We chose MI-09 as a representative for the structural characterization due to its high potency. MI-09 was a previously published compound that showed antiviral potency in mice. However, the structure of its complex with M^Pro^ was not reported. We used our established soaking method to determine and refine the structure of M^Pro^ bound with MI-09 to a resolution ofÅ. As shown in Figure 4C, the electron density at the active site of M^Pro^-MI-09 showed a defined confirmation for the P1 and P2 residues and hemithioacetal hydroxide that was generated after the covalent interaction between MI-09 and C145 of M^Pro^. However, the MI-09 *N*-terminal groups adopted two confirmations that were clearly observable. The collected data allowed the refinement only for the *N*-terminal phenyl group but not its 4-trifluoromethoxy substituent. Except at the *N*-terminal group, MI-09 involved hydrogen bonds formed with M^Pro^ as same as MPI48. The P2 side chain of MI-09 has a rigid confirmation that fit nicely to the S2 pocket of M^Pro^. The P2 proline backbone in MI-09 pushed the side chain of Q189 to adopt a confirmation differ from that in M^Pro^-MPI48. As observed in other M^Pro^-inhibitor structures, the aa45-51 region had an undefined confirmation.

In all three determined structures, the inhibitors did not fully engage the S4 pocket. MPI48 and MPI49 had a refined *N*-terminal group in their M^Pro^ complexes due to its rigidity. But the *N*-terminal group in either molecule did not interact with the S4 pocket. MI-09 showed two confirmations for its *N*-terminal group at the active site of M^Pro^ indicating no strong interactions with the S4 pocket. A similar flexible *N*-terminal group has been observed in other M^Pro^-dipeptidyl inhibitor complexes as well. For future designs of dipeptidyl inhibitors, novel *N*-terminal groups that can better engage the S4 pocket will certainly improve the binding affinity to M^Pro^ and requires innovative inputs.

### Cellular M^Pro^ Inhibition and Antiviral Potency of Dipeptidyl Inhibitors

M^Pro^ showed acute toxicity to human cells when it was recombinantly expressed. Based on this observation, we developed a cell-based assay to characterize cellular potency of M^Pro^ inhibitors.^12^ In this assay, an inhibitor with cellular potency suppresses cytotoxicity from an M^Pro^-eGFP (enhanced green fluorescent protein) fusion protein that is transiently expressed in 293T cells and consequently leads to host cell survival and enhanced overall expression of M^Pro^-eGFP that can be characterized by flow cytometry. This assay allows quick assessment of M^Pro^ inhibitors in cells by avoiding tedious characterizations of cellular permeability and stability of developed compounds. This assay is also more accurate in assessing M^Pro^ inhibition in cells than a direct antiviral assay since a compound may block functions of host proteases such as TMPRSS2, furin, and cathepsin L that are critical for SARS-CoV-2 infection and therefore provides false positive results about M^Pro^ inhibition in cells.^13–15^ Using this cellular assay system, we previously characterized a number of repurposed SARS-CoV-2 inhibitors and pointed out that some repurposed inhibitors inhibit SARS-CoV-2 via mechanisms different from M^Pro^ inhibition.^12^ Compounds that showed potency in this cellular assay displayed roughly similar potency in antiviral tests. Using this cellular assay, we characterized all synthesized inhibitors in this work that displayed an enzymatic inhibition IC_50_ value below 0.5 M. The characterized cellular M^Pro^ inhibition EC_50_ values are presented in Table 1. For Group A compounds with *t*-butylalanine (**j**) and cyclohexylalanine (**k**) at the P2 site, they showed measurable cellular EC_50_ values. Four compounds MPI49-MPI52 had determined cellular EC_50_ values below 1 M. In comparison to other peptidyl inhibitors with leucine (**h**) and phenylalanine (**i**) at the P2 site, these compounds showed generally better cellular M^Pro^ inhibition potency indicating both **j** and **k** are optimal primary amino acid residues at the P2 site for improved cellular M^Pro^ inhibition potency. **J** and **k** at the P2 site likely improve the cellular permeability or stability of their containing compounds in cells. MPI54 that has a P2 *O*-*t*-butylthreonine (**l**) showed very weak cellular potency. In a previous work, we showed that a P3 *O*-*t*-butylthreonine (**l**) generally improves the cellular potency of tripeptidyl M^Pro^ inhibitors. The low cellular potency of MPI52 indicated that a similar effect cannot be achieved by moving *O*-*t*-butylthreonine (**l**) from P3 to the P2 site.

**Figure 6:**
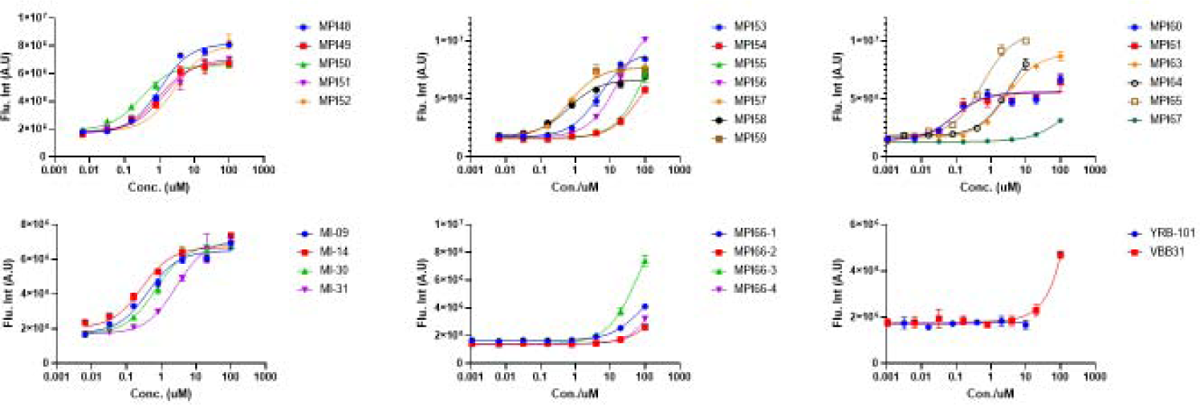
Cellular potency of inhibitors in their inhibition of M^Pro^ to drive host 293T cell survival and overall M^Pro^-eGFP expression.

**Figure 7:**
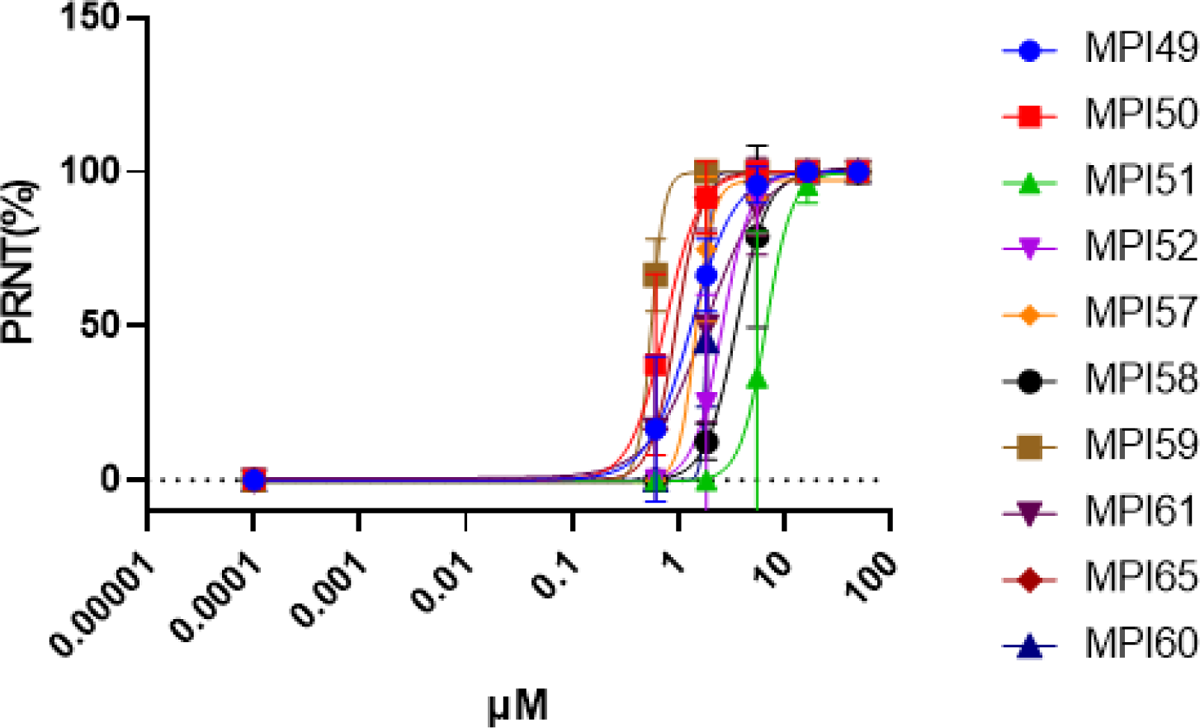
Plaque reduction neutralization tests (PRNTs) of selected compounds on their inhibition of three SARS-CoV-2 in Vero E6 cells. Two repeats were conducted for each concentration.

**Figure 8:**
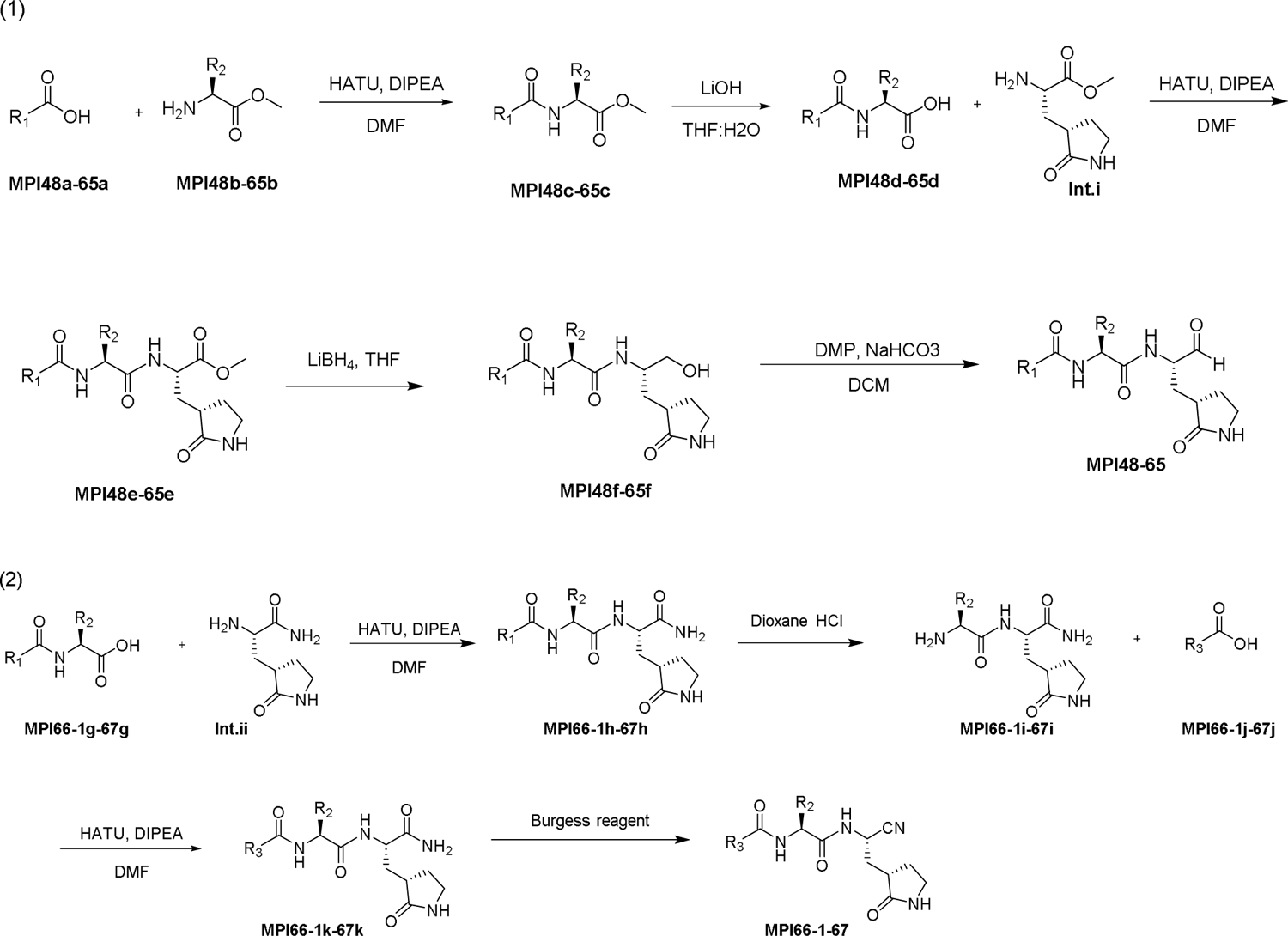
General synthesis route for compounds studied.

All Group B compounds with an IC_50_ value below 0.5 µ M had measurable cellular M^pro^ inhibition potency. The two most potency compounds are MPI60 and MPI61that showed determined cellular EC_50_ values below 100 nM. One interesting observation was that among MPI57-MPI61, all spiro compounds performed better than MPI57 that has a P2 bicyclic residue as in nirmatrelvir. Among all spiro compounds MPI58-MPI61, cellular potency was positively correlated with the size of the P2 spiro structure. The large size of the P2 residue likely improves cellular permeability of their containing compounds. For the four previously reported compounds MI-09, MI-14, MI-30, and MI-31, they displayed mild (> 0.5 µ M) to high (< 0.5 µ M) cellular potency but none of their potency reached to the level of MPI60 and MPI61. Compared to MPI57, **q** as the *N*-terminal group in MI-14 led to better cellular potency. However, replacing the *N*-terminal group in MPI58 with **q** to afford MPI63 led to worse cellular potency indicating the *N*-terminal group effects in cellular potency cannot be generalized. Replacing the *N*-terminal CBZ (**a**) group of MPI57 with **u** in MPI64 and **t** in MPI65 had opposite effects with worse cellular potency for MPI64 and better cellular potency for MPI65. However, the cellular potency of MPI65 was still significantly lower than that of MPI60 and MPI61. Among all inhibitors in both Group A and Group B, MPI60 and MPI61 had the highest cellular potency. Peptidyl M^Pro^ inhibitors with a P2 primary amino acid or bicyclic residue have been extensively explored. However, Peptidyl M^Pro^ inhibitors with a P2 spiro residue have not been studied much. As the first of its kind, the current work revealed that peptidyl M^Pro^ inhibitors with a P2 **a1** or **a2** residue significantly outperform other inhibitors.

For newly developed inhibitors that showed cellular potency with an EC_50_ value below 1 µ, we went further to characterize their antiviral potency. MPI57 was included for comparison with MPI58-MPI61. For four previously developed compounds MI-09, MI-14, MI-30, and MI-31, their antiviral tests were not conducted since they were characterized in previously reported paper ^1^ and their detected cellular potency was lower than that of MPI60 and MPI61. To quantify antiviral EC_50_, we conducted plaque reduction neutralization tests of three SARS-CoV-2 variants including USA-WA1/2020 in Vero E6 cells for all four inhibitors. We infected Vero E6 cells by virus in the presence of an inhibitor at various concentrations for three days and then quantified viral plaque reduction. Based on viral plaque reduction data, we determined antiviral EC_50_ values for all tested inhibitors. The determined antiviral EC_50_ values are presented in Table 1. They all had measurable antiviral EC_30_ values below 10 µ M. Five compounds MPI50, MPI59, MPI60, MPI61, and MPI65 had antiviral EC_50_ values below 1 µ M. As same as shown in the cellular potency tests, MPI60 and MPI61 had the highest antiviral potency with a same EC_50_ value as 0.37 µ M. So, we can conclude that a P2 **a1** or **a2** residue in a dipeptidyl M^Pro^ inhibitor leads to optimal antiviral potency and performs better than **x** that has been used in nirmatrelvir.

### Cytotoxicity and *In Vitro* Metabolic Stability of MPI60 and MPI61

Due to their high antiviral potency, MPI60 and MPI61 were advanced to characterizations of cytotoxicity and *in vitro* metabolic stability. For the cytotoxicity characterization, we used 293T cells and the MTT assay.^16^ Determined CC_50_ values for MPI60 and MPI61 were 95 and 230 µ M, respectively and presented in Table 1 as well. Both MPI60 and MPI61 showed low toxicity. These CC_50_ values are similar to that of nirmatrelvir and lead to high calculated selectivity indices (CC_50_/antiviral EC_50_) for both compounds. The *in vitro* metabolic stability analysis for MPI60 and MPI61 was conducted using human liver microsomes. Their determined CL_int_ values were 11 and 45 µ L/min/kg, respectively, compared to 24.5 µ L/min/kg, a literature value for nirmatrelvir. All collected data combinedly showed that MPI60 has probably the most favorable characteristics for moving forward to animal studies.

## Conclusion

We have systematically surveyed reversibly covalent dipeptidyl M^Pro^ inhibitors on their characteristics including enzymatic inhibition, crystal structures of their complexes with M^Pro^, cellular and antiviral potency, cytotoxicity, and *in vitro* metabolic stability. Our results showed that the M^Pro^ S2 pocket is flexible in accommodating large P2 residues in dipeptidyl M^Pro^ inhibitors and inhibitors with two large P2 spiro residues, (*S*)-2-azaspiro[4,4]nonane-3-carboxylate (**a1**) and (*S*)-2-azaspiro[4,5]decane-3-carboxylate (**a2**) are optimal on most characteristics. One compound MPI60 displayed high antiviral potency and *in vitro* metabolic stability, suggesting that it is ready for the next level of preclinical assessment.

### Experimental Section Materials

HEK293T/17 cells were from ATCC; DMEM with high glucose with GlutaMAX supplement, fetal bovine serum, 0.25% trypsin-EDTA, phenol red and dimethyl sulfoxide from Thermo Fisher Scientific; linear polyethylenimine MW 25000 from Polysciences;

### *In Vitro* M^Pro^ Inhibition Potency Characterizations of MPIs

For most MPIs, the assay was conducted using 20 nM M^Pro^ and 10 M Sub3. We dissolved all inhibitors in DMSO as 10 mM stock solutions. Sub3 was dissolved in DMSO as a 1 mM stock solution and diluted 100 times in the final assay buffer containing 10 mM Na_x_H_y_PO_4_, 10 mM NaCl, 0.5 mM EDTA, and 1.25% DMSO at pH 7.6. M^Pro^ and an inhibitor were incubated in the final assay buffer for 30 min before adding the substrate to initiate the reaction catalyzed by M^Pro^. The production format was monitored in a fluorescence plate reader with excitation at 336 nm and emission at 490 nm. More assay details can be found in a previous study.

### Cellular M^Pro^ Inhibition Potency Characterizations of MPIs

HEK 293T/17 cells were grown in high-glucose DMEM with GlutaMAX supplement and 10% fetal bovine serum in 10 cm culture plates under 37 °C and 5% CO2 to ∼80−90% and then transfected cells with the pLVX-MProeGFP-2 plasmid. For each transfection, 30 mg/mL polyethylenimine and a total of 8 μg of the plasmid in 500 μL of the opti-MEM medium were used. Cells were incubated with transfecting reagents overnight. On the second day, remove the medium, wash cells with a PBS buffer, digest them with 0.05% trypsin-EDTA, resuspend the cells in the original growth media, adjustthe cell density to 5 × 10^5^ cells/mL, provide 500 μL of suspended cells in the growth media to each well of μL of a drug solution to the growth media. These cells were incubated under 37L and 5% CO_2_ before flow cytometry analysis.

### Recombinant M^Pro^ protein expression and purification

The pET28a-His-SUMO-M^Pro^ expression and purification are according to our previous report.^1^ The pET28a-His-SUMO-M^Pro^ construct was transformed into *E. coli* BL21(DE3) cells. Transformed cells were cultured at 37°C in 2xYT medium with kanamycin (50 g/mL) until OD_600_ reaching 0.6, then induced with 1 mM isopropyl-D-1-thiogalactoside (IPTG) at 37 °C. After 3 hours, cells were harvested and lysed in buffer A (20 mM Tris, 100 mM NaCl, 10 mM imidazole, pH 8.0). The supernatant was loaded onto a nickel-chelating column (GenScript) washed with buffer A, followed by elution with buffer B (20 mM Tris, 100 mM NaCl, 250 mM imidazole, pH 8.0). The eluted protein solution was desalted to buffer C (20 mM Tris, 10 mM NaCl, pH 8.0) by HiPrep 26/10 desalting column (GE Healthcare). The His-SUMO-M^Pro^ proteins were digested with SUMO protease overnight at 4°C. The digested protein was applied to nickel-chelating column again to remove the His-tagged SUMO protease, the His-SUMO tag, and the expressed protein with uncleaved His-SUMO tag. The tag-free M^Pro^ protein was loaded to the size exclusion column HiPrep 16/60 Sephacryl S-100 HR (GE Healthcare) pre-equilibrated with buffer D (20 mM Tris, 100 mM NaCl, 1 mM EDTA, pH 7.8). The eluted M^Pro^ protein was stored with buffer D in −80 °C for further use.

### X-Ray Crystallography Analysis

The production of crystals of apo M^Pro^ and M^Pro^-inhibitor complexes was following the previous protocols with the crystal growing condition of 0.1 M Bis-tris, pH6.5, 16% w/v PEG10k.^16^ The data of M^Pro^ with MPI48, MPI49, MI-09 were collected on a Bruker Photon II detector. The diffraction data were indexed, integrated and scaled with iMosflm or PROTEUM3^17^. The structure was determined by molecular replacement using the structure model of the free enzyme of the SARS-CoV-2 M^pro^ [Protein Data Bank (PDB) ID code 7JPY] as the search model using Phaser in the Phenix package ^16, 19^. *JLigand* and *Sketcher* from the CCP4 suite were employed for the generation of PDB and geometric restraints for the inhibitors. The inhibitors were built into the *Fo-Fc* density by using *Coot* ^20^. Refinement of all the structures was performed with Real-space Refinement in Phenix ^19^. Details of data quality and structure refinement are summarized in Table S1. All structural figures were generated with PyMOL (https://www.pymol.org).

### Compound Synthesis

**General procedure A**: To a solution of **a** (1 eq) and b (1 eq) in anhydrous DMF was added DIPEA (4 eq) and was cooled to 0 °C. HATU (1.2 eq) was added to the solution under 0 °C and then stirred at room temperature overnight. The reaction mixture was then diluted with ethyl acetate and washed with saturated NaHCO_3_ solution (2 times), 1 M HCl solution (2 times), and saturated brine solution (2 times) sequentially. The organic layer was dried over anhydrous Na_2_SO_4_ and then concentrated *on vacuum*. The residue was then purified with flash chromatography (50-100% EtOAc in hexanes as the eluent) to afford **c** as white solid/gummy solid.

**General procedure B**: The compound **c** (1 eq) was dissolved in THF/H_2_O (1:1). LiOH (2.5 eq) was added at 0 °C. The mixture was stirred at room temperature overnight. Then THF was removed *on vacuum* and the aqueous layer was acidified with 1 M HCl and extracted with dichloromethane (3 times). The organic layer was dried over anhydrous Na_2_SO_4_ and concentrated to yield **d** as white solid/ gummy solid. Proceeded for next step without further purification.

**General procedure C**: To a solution of **d** (1 eq) and **Int.i** (1 eq) in anhydrous DMF was added DIPEA (4 eq) and was cooled to 0 °C. HATU (1.2 eq) was added to the solution under 0 °C and then stirred at room temperature overnight. The reaction mixture was then diluted with ethyl acetate and washed with saturated NaHCO_3_ solution (2 times), 1 M HCl solution (2 times), and saturated brine solution (2 times) sequentially. The organic layer was dried over anhydrous Na_2_SO_4_ and then concentrated *on vacuum*. The residue was then purified with flash chromatography (0-10% MeOH in dichloromethane as the eluent) to afford **e** as white solid/gummy solid.

**General procedure D**: To a stirred solution of compound **e** (1 eq) in THF was added LiBH_4_ (2.0 M in THF, 5 eq) in several portions at 0 °C under a nitrogen atmosphere. The reaction mixture was stirred at 0 °C for 1 h, then allowed to warm up to room temperature, and stirred for an additional 2 h. The reaction was quenched by the drop wise addition of 1.0 M HCl (aq) (1.2 mL) with cooling in an ice bath. The solution was diluted with ethyl acetate and H_2_O. The phases were separated, and the aqueous layer was extracted with ethyl acetate (3 times). The organic phases were combined, dried over MgSO_4_, filtered, and concentrated on a rotorvap to give a yellow oily residue. Column chromatographic purification of the residue (2-10% MeOH in CH_2_Cl_2_ as the eluent) afforded **f** as white solid/ gummy solid.

**General procedure E**: To a solution of **f** in CH_2_Cl_2_ was added NaHCO_3_ (4 eq) and the Dess-Martin reagent (3 eq). The resulting mixture was stirred at rt for 12 h. Then the reaction was quenched with a saturated NaHCO_3_ solution containing 10 % Na_2_S_2_O_3_. The layers were separated. The organic layer was then washed with saturated brine solution, dried over anhydrous Na_2_SO_4,_ and concentrated *on vacuum*. The residue was then purified with flash chromatography afford **MPI** compounds as white solid.

**General procedure F**: To a stirred solution of **Compound** (1 eq) in 1,4-Dioxane at 0 °C was added 4N HCl (10 eq). Reaction mixture was stirred at rt for 3 h. After completion of reaction, solvent was concentrated in a vacuum. The residue was used in the next step without further purification.

**General procedure G**: To a stirred solution of Compound (1 eq) in THF at 0 °C was added DIPEA (2 eq). After 15 min, Cbz-Cl (1.2 eq) was added, and the mixture was stirred at rt for 3 h. The reaction was quenched with water (5 mL), and the mixture was concentrated in a vacuum. The residue was partitioned between EtOAc (10 mL) and H_2_O (5 mL). The aqueous layer was extracted with EtOAc (2 times). The combined organic layer was washed with brine, dried over MgSO_4,_ and concentrated in a vacuum. The residue was then purified with flash chromatography (0-10% MeOH in CH_2_Cl_2_ as the eluent) to afford **k** as a yellow liquid.

**General procedure H**: To a stirred solution of **K** (1 eq) in DCM (10 mL) at 0 °C was added Burgess reagent (2.5 eq) and the mixture was stirred at rt for 2 h. The reaction was quenched with saturated NaHCO_3_ solution (5 mL) and extracted with DCM (2 × 10 mL). The combined organic layer was washed with brine, dried over MgSO_4,_ and concentrated in a vacuum. The residue was then purified with flash chromatography (0-10% MeOH in Dichloromethane as the eluent) to afford MPI66-1-67 as a white solid.

### Methyl (S)-2-(1H-indole-2-carboxamido)-4,4-dimethylpentanoate (MPI48c)

**MPI48c** was prepared with methyl (S)-2-amino-4,4-dimethylpentanoate hydrochloride (MPI48b) and 1H-indole-2-carboxylic acid (MPI48a) as a white solid following a general procedure **A** (yield 70%).

### (S)-2-(1H-indole-2-carboxamido)-4,4-dimethylpentanoic acid (MPI48d)

**MPI48d** was prepared as a white solid following a general procedure **B**.

### methyl (S)-2-((S)-2-(1H-indole-2-carboxamido)-4,4-dimethylpentanamido)-3-((S)-2-oxopyrrolidin-3-yl)propanoate (MPI48e)

**MPI48e** was prepared with Int.i and 1H-indole-2-carboxylic acid (MPI48d) as a white solid following a general procedure **C** (yield 60%). ^1^H NMR (400 MHz, DMSO-d6) δ11.57 (s, 1H), 8.48 (dd, J = 21.5, 8.1 Hz, 2H), 7.70 – 7.58 (m, 2H), 7.42 (d, J = 8.2 Hz, 1H), 7.23 (s, 1H), 7.17 (t, J = 7.6 Hz, 1H), 7.03 (t, J = 7.4 Hz, 1H), 4.68 – 4.55 (m, 1H), 4.38 – 4.28 (m, 1H), 3.61 (s, 3H), 3.17 – 3.02 (m, 2H), 2.38 – 2.27 (m, 1H), 2.16 – 2.02 (m, 2H), 1.82 – 1.52 (m, 4H), 0.94 (s, 9H).

### N-((S)-1-(((S)-1-hydroxy-3-((S)-2-oxopyrrolidin-3-yl)propan-2-yl)amino)-4,4-dimethyl-1-oxopentan-2-yl)-1H-indole-2-carboxamide (MPI48f)

**MPI48f** was prepared as a white solid following a general procedure **D** (yield 60%). ^1^H NMR (400 MHz, DMSO-d6) δ 11.57 (s, 1H), 8.48 (dd, J = 21.5, 8.1 Hz, 2H), 7.70 – 7.58 (m, 2H), 7.42 (d, J = 8.2 Hz, 1H), 7.23 (s, 1H), 7.17 (t, J = 7.6 Hz, 1H), 7.03 (t, J = 7.4 Hz, 1H), 4.68 – 4.55 (m, 1H), 4.38 – 4.28 (m, 1H), 3.61 (s, 3H), 3.17 – 3.02 (m, 2H), 2.38 – 2.27 (m, 1H), 2.16 – 2.02 (m, 2H), 1.82 – 1.52 (m, 4H), 0.94 (s, 9H).

### N-((S)-4,4-dimethyl-1-oxo-1-(((S)-1-oxo-3-((S)-2-oxopyrrolidin-3-yl)propan-2-yl)amino)pentan-2-yl)-1H-indole-2-carboxamide (MPI48)

**MPI48** was prepared as a white solid following a general procedure **E** (yield 45%). ^1^H NMR (400 MHz, DMSO-d6) δ 11.60 (s, 1H), 9.42 (s, 1H), 8.71 – 8.44 (m, 2H), 7.62 (d, J = 9.5 Hz, 2H), 7.42 (d, J = 8.3 Hz, 1H), 7.25 (s, 1H), 7.18 (t, J = 7.5 Hz, 1H), 7.03 (t, J = 7.5 Hz, 1H), 4.62 (td, J = 9.0, 3.3 Hz, 1H), 4.20 (ddt, J = 14.9, 11.4, 5.6 Hz, 1H), 3.19 – 2.95 (m, 2H), 2.39 – 2.21 (m, 1H), 2.21 – 2.06 (m, 1H), 1.97 – 1.88 (m, 1H), 1.88 – 1.72 (m, 2H), 1.72 – 1.55 (m, 2H), 0.94 (s, 9H). ^13^C NMR (101 MHz, δDMSO) 201.3, 178.8, 173.8, 161.2, 136.9, 131.9, 127.5, 123.9, 122.0, 120.2, 112.7, 103.9, 60.2, 56.9, 51.1, 45.1, 37.8, 30.9, 30.1, 29.8, 27.8.

### (S)-methyl 2-(4-methoxy-1H-indole-2-carboxamido)-4,4-dimethylpentanoate (MPI49c)

**MPI49c** was prepared with methyl (S)-2-amino-4,4-dimethylpentanoate hydrochloride (MPI48b) and 4-methoxy-1H-indole-2-carboxylic acid (MPI49a) as a white solid following general procedure **A** (yield 79%). ^1^H NMR (400 MHz, Chloroform-d) δ 10.02 (s, 1H), 7.18 (t, J = 8.0 Hz, 1H), 7.09 – 7.03 (m, 2H), 6.77 (d, J = 8.6 Hz, 1H), 6.48 (d, J = 7.7 Hz, 1H), 4.94 (td, J = 8.9, 3.6 Hz, 1H), 3.93 (s, 3H), 3.76 (s, 3H), 1.89 (dd, J = 14.4, 3.6 Hz, 1H), 1.68 (dd, J = 14.4, 9.1 Hz, 1H), 1.02 (s, 9H). ^13^C NMR (101 MHz, CDCl3): δ 173.99, 161.39, 154.13, 138.11, 128.91, 125.47, 118.84, 105.32, 100.55, 99.53, 60.44, 55.26, 52.51, 50.14, 30.80, 29.67.

### (S)-2-(4-methoxy-1H-indole-2-carboxamido)-4,4-dimethylpentanoic acid (MPI49d)

**MPI49d** was prepared as a white solid following a general procedure **B** (290 mg, 86%). ^1^H NMR (400 MHz, DMSO-d6) δ 12.60 (s, 1H), 11.58 (s, 1H), 8.53 (d, J = 8.3 Hz, 1H), 7.33 (d, J = 5.2 Hz, 1H), 7.17 – 6.93 (m, 2H), 6.51 (t, J = 6.2 Hz, 1H), 4.52 (t, J = 8.4 Hz, 1H), 3.89 (s, 3H), 1.93 – 1.80 (m, 1H), 1.78 – 1.70 (m, 1H), 0.95 (s, 9H). 13C NMR (101 MHz, DMSO): δ 175.06, 161.16, 154.08, 138.29, 130.48, 124.89, 118.54, 105.88, 101.34, 99.65, 60.23, 55.50, 49.95, 44.16, 30.89, 29.84.

### (S)-methyl 2-((S)-2-(4-methoxy-1H-indole-2-carboxamido)-4,4-dimethylpentanamido)-3-((S)-2-oxopyrrolidin-3-yl)propanoate (MPI49e):)

**MPI49e** was prepared with Int.i and MPI49d as a white gummy solid following general procedure **C** (yield 72%). ^1^H NMR (400 MHz, Methanol-d4) δ 8.47 (dd, J = 28.2, 7.9 Hz, 1H), 8.22 (dd, J = 17.4, 8.1 Hz, 1H), 7.14 (d, J = 2.8 Hz, 1H), 7.03 (td, J = 8.0, 2.7 Hz, 1H), 6.92 (dd, J = 8.3, 4.9 Hz, 1H), 6.39 (dd, J = 7.8, 2.7 Hz, 1H), 4.68 – 4.59 (m, 1H), 4.48 – 4.35 (m, 1H), 3.81 (s, 3H), 3.60 (s, 2H), 3.16 – 3.02 (m, 2H), 2.51 – 2.39 (m, 1H), 2.26 – 2.02 (m, 2H), 1.86 – 1.55 (m, 4H), 0.92 (s, 9H). ^13^C NMR (101 MHz, MeOD): δ 180.37, 174.37, 172.27, 162.14, 154.24, 138.41, 129.05, 124.97, 118.71, 104.89, 104.81, 101.53, 98.93, 54.32, 51.47, 51.15, 50.69, 44.44, 40.03, 38.18, 30.13, 30.07, 28.81, 27.29.

### N-((S)-1-(((S)-1-hydroxy-3-((S)-2-oxopyrrolidin-3-yl)propan-2-yl)amino)-4,4-dimethyl-1-oxopentan-2-yl)-4-methoxy-1H-indole-2-carboxamide (MPI49f)

**MPI49f** was prepared as a white solid following a general procedure **D** (yield 58%). ^1^H NMR (400 MHz, Chloroform-d) δ 10.39 (s, 1H), 8.06 (d, J = 8.0 Hz, 1H), 7.20 – 7.12 (m, 1H), 7.10 – 6.98 (m, 2H), 6.91 (s, 1H), 6.55 – 6.37 (m, 1H), 5.92 (s, 1H), 4.81 – 4.63 (m, 1H), 4.10 – 3.98 (m, 1H), 3.93 (s, 3H), 3.72 – 3.58 (m, 2H), 3.24 – 2.90 (m, 2H), 2.45 – 2.33 (m, 1H), 2.28 – 2.17 (m, 1H), 2.13 – 1.92 (m, 3H), 1.63 (dd, J = 13.8, 8.8 Hz, 2H), 1.03 (s, 3H), 0.96 (s, 6H).

### N-((S)-4,4-dimethyl-1-oxo-1-(((S)-1-oxo-3-((S)-2-oxopyrrolidin-3-yl)propan-2-yl)amino)pentan-2-yl)-4-methoxy-1H-indole-2-carboxamide (MPI49)

**MPI49** was prepared as a white solid following a general procedure **E** (yield 72%). ^1^H NMR (400 MHz, Chloroform-d) δ 10.85 (s, 1H), 9.69 – 9.50 (m, 1H), 9.41 (s, 1H), 8.41 (d, J = 6.1 Hz, 1H), 7.15 – 7.09 (m, 1H), 7.07 – 7.00 (m, 1H), 6.97 – 6.92 (m, 1H), 6.73 (dd, J = 18.8, 8.3 Hz, 1H), 6.42 (dd, J = 7.7, 3.6 Hz, 1H), 5.96 (s, 1H), 4.90 – 4.71 (m, 1H), 4.22 (ddt, J = 23.4, 11.1, 5.5 Hz, 1H), 3.87 (s, 3H), 3.29 – 2.90 (m, 2H), 2.44 – 2.17 (m, 2H), 2.03 – 1.79 (m, 3H), 1.72 – 1.63 (m, 2H), 0.94 (s, 9H). ^13^C NMR (101 MHz, CDCl3): δ199.82, 180.19, 173.89, 161.58, 154.11, 138.50, 138.12, 128.96, 125.66, 118.81, 108.03, 99.61, 57.77, 55.30, 51.84, 50.84, 40.99, 37.63, 30.66, 29.86, 29.79, 29.72, 28.35.

### (S)-2-(((benzyloxy)carbonyl)amino)-4,4-dimethylpentanoic acid (MPI50d)

**MPI50d** was prepared as a white solid following a general procedure **G** (yield 84%). ^1^H NMR (400 MHz, Chloroform-d) δ 7.15 (s, 5H), 5.71 (s, 1H), 5.08 (d, J = 12.5 Hz, 1H), 4.75 (d, J = 12.6 Hz, 1H), 4.10 (s, 1H), 1.67 (d, J = 14.3 Hz, 1H), 1.29 (dd, J = 14.4, 9.1 Hz, 1H), 0.79 (s, 9H). ^13^C NMR (100 MHz, Chloroform-d) δ 156.70, 136.50, 128.40, 127.89, 66.77, 30.53, 29.68.

### Methyl (S)-2-((S)-2-(((benzyloxy)carbonyl)amino)-4,4-dimethylpentanamido)-3-((S)-2-oxopyrrolidin-3-yl)propanoate (MPI50e)

**MPI50e** was prepared with Int.i and MPI50d as a white gummy solid following general procedure **C** (yield 67%). ^1^H NMR (400 MHz, Chloroform-d) δ 7.74 (d, J = 7.8 Hz, 1H), 7.32 – 7.22 (m, 5H), 6.78 (s, 1H), 5.46 (d, J = 9.2 Hz, 1H), 5.01 (s, 2H), 4.64 – 4.52 (m, 1H), 4.52 – 4.40 (m, 1H), 3.94 – 3.85 (m, 1H), 3.62 (s, 3H), 3.30 – 3.15 (m, 2H), 2.29 (dd, J = 7.1, 3.2 Hz, 2H), 2.23 – 2.10 (m, 1H), 2.10 – 2.00 (m, 1H), 1.80 – 1.68 (m, 3H), 0.85 (s, 9H). ^13^C NMR (100 MHz, Chloroform-d) δ 179.83, 173.02, 172.00, 170.90, 156.52, 136.22, 128.54, 128.18, 128.00, 67.07, 60.41, 52.33, 50.90, 46.48, 40.54, 30.95, 30.56, 29.55, 27.95, 21.06, 19.20, 18.08, 14.20.

### Benzyl ((S)-1-(((S)-1-hydroxy-3-((S)-2-oxopyrrolidin-3-yl)propan-2-yl)amino)-4,4-dimethyl-1-oxopentan-2-yl)carbamate (MPI50f)

**MPI50f** was prepared as a white solid following a general procedure **D** (yield 51%). ^1^H NMR (400 MHz, Chloroform-d) δ 7.41 – 7.22 (m, 5H), 6.50 (d, J = 6.3 Hz, 1H), 5.93 (s, 1H), 5.41 (s, 1H), 5.07 (d, J = 4.1 Hz, 2H), 4.17 (d, J = 2.9 Hz, 1H), 3.97 (s, 1H), 3.60 (dd, J = 11.8, 3.4 Hz, 1H), 3.50 (dd, J = 11.7, 6.7 Hz, 1H), 3.26 – 3.05 (m, 2H), 2.40 – 2.29 (m, 1H), 2.29 – 2.19 (m, 1H), 2.08 (d, J = 15.7 Hz, 1H), 1.97 – 1.77 (m, 2H), 1.77 – 1.61 (m, 1H), 0.86 (s, 9H). 13C NMR (100 MHz, Chloroform-d) δ 180.44, 173.62, 173.42, 156.25, 135.91, 128.67, 128.46, 128.37, 67.62, 65.70, 63.37, 53.04, 50.63, 45.79, 40.33, 38.10, 35.36, 32.36, 30.69, 29.69, 28.37, 17.45, 17.11, 17.00.

### Benzyl ((S)-4,4-dimethyl-1-oxo-1-(((S)-1-oxo-3-((S)-2-oxopyrrolidin-3-yl)propan-2-yl)amino)pentan-2-yl)carbamate (MPI50)

**MPI50** was prepared as a white solid following a general procedure **E** (yield 25%). ^1^H NMR (400 MHz, Chloroform-d) δ 9.46 (s, 1H), 8.01 (d, J = 7.7 Hz, 1H), 7.28 (d, J = 4.1 Hz, 5H), 6.71 (d, J = 7.4 Hz, 1H), 6.28 – 6.15 (m, 1H), 5.41 (s, 1H), 5.08 – 4.92 (m, 2H), 4.45 – 4.33 (m, 1H), 4.17 (d, J = 10.3 Hz, 1H), 3.26 – 3.07 (m, 2H), 2.28 (d, J = 16.7 Hz, 2H), 2.01 (p, J = 7.4, 6.7 Hz, 2H), 1.89 (d, J = 14.1 Hz, 1H), 1.74 (dd, J = 45.0, 9.9 Hz, 2H), 1.65 (s, 1H), 0.88 (s, 9H). ^13^C NMR (100 MHz, Chloroform-d) δ 200.61, 173.84, 173.18, 156.24, 136.07, 128.63, 128.35, 128.17, 67.27, 63.36, 57.13, 51.84, 45.79, 40.34, 34.88, 30.66, 30.24, 29.67, 28.12, 17.37, 17.10.

### methyl (S)-2-((S)-2-(((benzyloxy)carbonyl)amino)-3-cyclohexylpropanamido)-3-((S)-2-oxopyrrolidin-3-yl)propanoate (MPI51e)

**MPI51e** was prepared with Int.i and (S)-2-(((benzyloxy)carbonyl)amino)-3-cyclohexylpropanoic acid (MPI51d) as a white gummy solid following general procedure **C** (yield 67%). ^1^H NMR (400 MHz, Chloroform-d) δ 7.69 (d, J = 7.0 Hz, 1H), 7.42 – 7.29 (m, 5H), 5.92 (s, 1H), 5.29 (d, J = 8.7 Hz, 1H), 5.18 – 5.04 (m, 2H), 4.50 (s, 1H), 4.29 (d, J = 6.3 Hz, 1H), 3.73 (s, 3H), 3.39 – 3.25 (m, 2H), 2.42 (s, 2H), 2.19 – 2.10 (m, 1H), 1.97 – 1.57 (m, 4H), 1.57 – 1.33 (m, 3H), 1.32 – 1.05 (m, 4H), 1.05 – 0.76 (m, 4H).

### benzyl ((S)-3-cyclohexyl-1-oxo-1-(((S)-1-oxo-3-((S)-2-oxopyrrolidin-3-yl)propan-2-yl)amino)propan-2-yl)carbamate (MPI51)

**MPI51** was prepared as a white solid following a general procedures **D** and **E** (yield 47%). ^1^H NMR (400 MHz, DMSO-d6) δ 9.40 (s, 1H), 8.48 (d, J = 7.6 Hz, 1H), 7.63 (s, 1H), 7.47 (d, J = 8.1 Hz, 1H), 7.38 – 7.27 (m, 5H), 5.03 (s, 2H), 4.18 (ddd, J = 11.4, 7.4, 4.0 Hz, 0H), 4.10 (q, J = 7.8 Hz, 1H), 3.24 – 2.96 (m, 3H), 2.36 – 2.21 (m, 1H), 2.21 – 2.04 (m, 1H), 1.95 – 1.83 (m, 1H), 1.77 – 1.53 (m, 6H), 1.53 – 1.40 (m, 2H), 1.39 – 1.27 (m, 1H), 1.25 – 1.01 (m, 4H), 0.98 – 0.76 (m, 2H). ^13^C NMR (101 MHz, CDCl3) δ 199.9, 180.3, 173.9, 156.3, 136.5, 128.6, 128.3, 128.1, 67.0, 57.9, 40.8, 40.7, 38.3, 34.2, 33.8, 32.6, 29.8, 28.8, 26.5, 26.3, 26.2.

### (S)-methyl 2-((((3-chlorobenzyl)oxy)carbonyl)amino)-3-cyclohexylpropanoate (MPI52c)

To 3,5-dichlorobenzyl alcohol (0.201 g, 1.39 mmol) in THF (5 mL) were added K_2_CO_3_ (193 mg, 1.39 mmol) and Triphosgene (166 mg, 0.56 mmol) and the mixture was stirred at room temperature for 1 h. The mixture was then poured into water (10 mL) and extracted with ethyl acetate (2×20 mL), Combine organic layers and dried over Na_2_SO_4_. The organic phase was evaporated to dryness and the crude material was used directly in the next step. 3,5-Dichlorobenzyl Chloroformate in THF (5 mL) was added to drop wise to a mixture of methyl (S)-2-amino-3-cyclohexylpropanoate (320 mg,1.39mmol) and DIPEA (0.3 ml, 2.78mmol).The reaction mixture stirred for 12 h. The mixture was then poured into water (30 mL) and extracted with ethyl acetate (4×20 mL). The organic layer was washed with aqueous hydrochloric acid 10% v/v (2×20 mL), saturated aqueous NaHCO_3_ (2×20 mL), brine (2×20 mL) and dried over Na_2_SO_4_. The organic phase was evaporated to dryness and the crude material purified by silica gel column chromatography (15-50% EtOAc in n-hexane as the eluent) to afford **MPI52c** white solid (280 mg, 59%). ^1^H NMR (400 MHz, Chloroform-d) δ 7.45 – 7.05 (m, 4H), 5.22 – 4.86 (m, 2H), 4.34 (td, J = 9.0, 5.1 Hz, 1H), 3.66 (s, 3H), 1.79 – 1.67 (m, 1H), 1.67 – 1.49 (m, 5H), 1.49 – 1.37 (m, 1H), 1.33 – 1.24 (m, 1H), 1.22 – 1.01 (m, 3H), 0.94 – 0.74 (m, 2H).

### (S)-2-((((3-chlorobenzyl)oxy)carbonyl)amino)-3-cyclohexylpropanoic acid (MPI52d)

**MPI52d** was prepared as a white solid following a general procedure **B**. ^1^H NMR (400 MHz, Chloroform-d) δ 8.13 (s, 1H), 7.27 (s, 1H), 7.21 (d, J = 4.4 Hz, 2H), 7.15 (d, J = 4.6 Hz, 1H), 5.28 – 4.79 (m, 2H), 4.44 – 4.16 (m, 1H), 1.82 – 1.70 (m, 1H), 1.69 – 1.53 (m, 5H), 1.51 – 1.42 (m, 1H), 1.38 – 1.28 (m, 1H), 1.22 – 1.02 (m, 3H), 0.95 – 0.77 (m, 2H).

### (S)-methyl 2-((S)-2-((((3-chlorobenzyl)oxy)carbonyl)amino)-3-cyclohexylpropanamido)-3-((S)-2-oxopyrrolidin-3-yl)propanoate (MPI52e)

**MPI50e** was prepared with Int.i and MPI52d as a white gummy solid following general procedure **C** (yield 54%). ^1^H NMR (400 MHz, Chloroform-d) δ 7.87 (d, J = 6.7 Hz, 1H), 7.37 (s, 1H), 7.29 (d, J = 2.8 Hz, 2H), 7.24 (t, J = 4.1 Hz, 1H), 6.01 (s, 1H), 5.39 (d, J = 8.6 Hz, 1H), 5.10 (s, 2H), 4.50 (s, 1H), 4.34 (d, J = 6.7 Hz, 1H), 3.75 (s, 3H), 3.45 – 3.30 (m, 2H), 2.52 – 2.34 (m, 2H), 2.23 – 2.00 (m, 3H), 1.99 – 1.81 (m, 3H), 1.76 – 1.68 (m, 4H), 1.56 – 1.49 (m, 1H), 1.31 – 1.13 (m, 3H), 1.06 – 0.90 (m, 2H).

### 2-chlorobenzyl ((S)-3-cyclohexyl-1-(((S)-1-hydroxy-3-((S)-2-oxopyrrolidin-3-yl)propan-2-yl)amino)-1-oxopropan-2-yl)carbamate (MPI52f)

**MPI52f** was prepared as a white solid following a general procedure D (yield 80%). ^1^H NMR (400 MHz, Chloroform-d) δ 7.75 (d, J = 7.2 Hz, 1H), 7.27 (s, 1H), 7.24 (s, 2H), 7.14 (t, J = 4.6 Hz, 1H), 6.16 (s, 1H), 5.52 (d, J = 8.2 Hz, 1H), 5.00 (s, 2H), 4.39 – 4.10 (m, 1H), 4.03 – 3.82 (m, 1H), 3.65 – 3.46 (m, 2H), 3.32 – 3.16 (m, 2H), 2.44 – 2.25 (m, 2H), 2.02 – 1.89 (m, 1H), 1.79 – 1.69 (m, 2H), 1.64 – 1.50 (m, 6H), 1.50 – 1.39 (m, 2H), 1.27 (s, 1H), 1.17 – 1.01 (m, 3H), 0.95 – 0.73 (m, 2H). ^13^C NMR (101 MHz, CDCl_3_) δ 181.09, 173.72, 155.96, 138.55, 134.37, 129.82, 128.20, 127.77, 125.82, 65.91, 53.29, 51.21, 40.67, 38.46, 34.12, 33.71, 32.52, 32.03, 30.96, 28.81, 26.38, 26.24, 26.05.

### 3-chlorobenzyl ((S)-3-cyclohexyl-1-oxo-1-(((S)-1-oxo-3-((S)-2-oxopyrrolidin-3-yl)propan-2-yl)amino)propan-2-yl)carbamate (MPI52)

**MPI52** was prepared as a white solid following a general procedure **E** (yield 80%). ^1^H NMR (400 MHz, Chloroform-d) δ 9.41 (s, 1H), 8.33 (d, J = 5.9 Hz, 1H), 7.42 – 7.03 (m, 4H), 6.27 (s, 1H), 5.47 (d, J = 8.5 Hz, 1H), 5.01 (s, 2H), 4.42 – 4.18 (m, 2H), 3.40 – 3.14 (m, 2H), 2.45 – 2.27 (m, 2H), 1.94 – 1.70 (m, 4H), 1.65 – 1.55 (m, 5H), 1.51 – 1.42 (m, 1H), 1.35 – 1.29 (m, 1H), 1.19 – 1.04 (m, 3H), 0.96 – 0.80 (m, 2H).^13^C NMR (101 MHz, CDCl_3_) δ 199.74, 180.22, 173.77, 155.88, 138.55, 134.36, 129.82, 128.19, 127.80, 125.83, 67.98, 65.92, 57.94, 40.77, 40.68, 38.33, 34.09, 33.67, 32.52, 29.67, 28.73, 26.38, 26.21, 26.03.

### 3-(((((S)-3-cyclohexyl-1-(((S)-1-hydroxy-3-((S)-2-oxopyrrolidin-3-yl)propan-2-yl)amino)-1-oxopropan-2-yl)carbamoyl)oxy)methyl)phenyl acetate (MPI53f)

To a stirred solution of 3-(hydroxymethyl)phenyl acetate (100 mg, 0.599 mmol) and DIPEA (0.31 mL, 1.79 mmol) in dry CH_2_Cl_2_ (10 mL) was added N,N’-disuccinimidyl carbonate (214 mg, 0.838 mmol) at 0 °C. After 10 h at rt, (S)-2-amino-3-cyclohexyl-N-((S)-1-hydroxy-3-((S)-2-oxopyrrolidin-3-yl)propan-2-yl)propanamide (186 mg, 0.599 mmol) was added one portion at 0 °C. After 10 h at rt, the reaction mixture was evaporated in vacuo. Purification by silica gel chromatography (Dichloromethane/MeOH = 9:1). 150 mg of compound isolated. Yield 50%. ^1^H NMR (400 MHz, Chloroform-d) δ 7.68 (d, J = 8.3 Hz, 1H), 7.33 (t, J = 7.8 Hz, 1H), 7.18 (d, J = 7.6 Hz, 1H), 7.07 (s, 1H), 7.01 (dd, J = 8.0, 2.3 Hz, 1H), 6.43 (d, J = 17.5 Hz, 1H), 5.70 (t, J = 10.0 Hz, 1H), 5.07 (s, 2H), 4.25 (dd, J = 8.9, 5.3 Hz, 1H), 4.02 – 3.90 (m, 1H), 3.57 (q, J = 8.4, 5.4 Hz, 2H), 3.24 (t, J = 8.4 Hz, 2H), 2.45 – 2.30 (m, 2H), 2.28 (s, 3H), 1.99 (ddd, J = 14.2, 11.0, 5.2 Hz, 1H), 1.81 – 1.73 (m, 2H), 1.64 (td, J = 11.0, 8.6, 4.8 Hz, 6H), 1.49 (td, J = 8.9, 8.3, 4.5 Hz, 1H), 1.33 (s, 1H), 1.16 (ddd, J = 25.5, 16.6, 10.9 Hz, 3H), 0.98 – 0.84 (m, 2H). 13C NMR (101 MHz, CDCl3) δ 181.06, 173.63, 169.52, 156.03, 150.76, 138.20, 129.53, 125.15, 121.26, 120.94, 66.09, 65.78, 53.28, 50.75, 40.68, 40.61, 38.37, 34.10, 33.70, 32.52, 32.15, 28.61, 26.40, 26.24, 26.05, 21.13.

### 3-(((((S)-3-cyclohexyl-1-oxo-1-(((S)-1-oxo-3-((S)-2-oxopyrrolidin-3-yl)propan-2-yl)amino)propan-2-yl)carbamoyl)oxy)methyl)phenyl acetate (MPI53)

**MPI53** was prepared as a white solid following a general procedure **E** (yield 80%). ^1^H NMR (400 MHz, Chloroform-d) δ 9.47 (s, 1H), 8.36 (d, J = 5.9 Hz, 1H), 7.34 (t, J = 7.9 Hz, 1H), 7.20 (d, J = 7.7 Hz, 1H), 7.10 (s, 1H), 7.02 (ddd, J = 8.1, 2.4, 1.0 Hz, 1H), 6.25 (d, J = 14.9 Hz, 1H), 5.47 (d, J = 8.5 Hz, 1H), 5.10 (s, 2H), 4.43 – 4.25 (m, 2H), 3.37 – 3.23 (m, 2H), 2.50 – 2.32 (m, 2H), 2.29 (s, 3H), 2.04 – 1.77 (m, 4H), 1.75 – 1.61 (m, 5H), 1.53 (ddd, J = 14.0, 9.2, 5.3 Hz, 1H), 1.38 (s, 1H), 1.26 – 1.11 (m, 3H), 0.94 (dd, J = 23.0, 11.7 Hz, 2H).

### Methyl (S)-2-((2S,3S)-2-(((benzyloxy)carbonyl)amino)-3-(tert-butoxy)butanamido)-3-((S)-2-oxopyrrolidin-3-yl)propanoate (MPI54e)

**MPI54e** was prepared with Int.i and N-((benzyloxy)carbonyl)-O-(tert-butyl)-L-allothreonine (MPI54e) as a white gummy solid following general procedure **C** (yield 91%). ^1^H NMR (400 MHz, Methanol-d_4_) δ 7.5 – 7.3 (m, 5H), 5.2 – 5.1 (m, 2H), 4.6 (dd, J = 11.4, 4.1 Hz, 1H), 4.2 (d, J = 6.2 Hz, 1H), 4.1 (p, J = 6.2 Hz, 1H), 3.8 (s, 3H), 3.3 – 3.2 (m, 2H), 2.6 (qd, J = 10.1, 3.9 Hz, 1H), 2.3 – 2.1 (m, 2H), 1.9 – 1.7 (m, 2H), 1.2 (d, J = 11.1 Hz, 12H). ^13^C NMR (100 MHz, Methanol-d_4_) δ 181.7, 173.4, 173.0, 158.3, 138.2, 129.4, 129.0, 128.8, 75.5, 68.5, 67.6, 62.0, 52.8, 51.8, 41.4, 39.4, 34.0, 28.6, 28.6, 19.9.

### Benzyl ((2S,3S)-3-(tert-butoxy)-1-(((S)-1-hydroxy-3-((S)-2-oxopyrrolidin-3-yl)propan-2-yl)amino)-1-oxobutan-2-yl)carbamate (MPI54f)

MPI54f was prepared as a white solid following a general procedure **D** (yield 91%). ^1^H NMR (400 MHz, Methanol-d_4_) δ 7.4 – 7.3 (m, 5H), 5.1 (s, 2H), 4.2 (d, J = 5.8 Hz, 1H), 4.1 – 3.9 (m, 2H), 3.6 (dd, J = 10.9, 4.8 Hz, 1H), 3.5 (dd, J = 11.0, 6.2 Hz, 1H), 3.3 (td, J = 9.3, 2.7 Hz, 1H), 3.2 – 3.2 (m, 1H), 2.6 – 2.4 (m, 1H), 2.4 – 2.2 (m, 1H), 2.1 – 1.9 (m, 1H), 1.8 (dq, J = 12.5, 8.9 Hz, 1H), 1.6 (ddd, J = 14.2, 11.1, 3.3 Hz, 1H), 1.3 – 1.1 (m, 12H). ^13^C NMR (100 MHz, Methanol-d_4_) δ 182.7, 172.9, 158.4, 138.2, 129.4, 129.0, 128.8, 75.5, 68.6, 67.7, 65.4, 62.3, 50.6, 41.5, 39.5, 33.6, 28.9, 28.7, 19.8.

### Benzyl ((2S,3S)-3-(tert-butoxy)-1-oxo-1-(((S)-1-oxo-3-((S)-2-oxopyrrolidin-3-yl)propan-2-yl)amino)butan-2-yl)carbamate (MPI54)

**MPI54** was prepared as a white solid following a general procedure **E** (yield 91%). ^1^H NMR (400 MHz, Chloroform-d) δ 9.4 (s, 1H), 8.1 (d, J = 6.6 Hz, 1H), 7.3 – 7.2 (m, 5H), 6.7 – 6.5 (m, 1H), 6.1 – 5.9 (m, 1H), 5.1 – 4.9 (m, 2H), 4.3 (s, 2H), 4.0 – 3.9 (m, 1H), 3.2 (d, J = 8.3 Hz, 2H), 2.5 – 2.3 (m, 1H), 2.2 (dd, J = 14.5, 6.6 Hz, 1H), 2.0 – 1.9 (m, 1H), 1.9 – 1.6 (m, 2H), 1.1 (s, 12H). ^13^C NMR (100 MHz, Chloroform-d) δ 199.8, 180.1, 171.2, 156.4, 136.2, 128.6, 128.3, 128.2, 74.7, 67.9, 67.3, 61.4, 55.1, 50.8, 40.6, 38.1, 29.9, 28.3, 19.6.

### Methyl (1R,2S,5S)-3-(2-(2,4-dichlorophenoxy)acetyl)-6,6-dimethyl-3-azabicyclo[3.1.0]hexane-2-carboxylate (MPI55c)

**MPI55c** was prepared with methyl (1R,2S,5S)-6,6-dimethyl-3-azabicyclo[3.1.0]hexane-2-carboxylate hydrogen chloride (MPI55b) and 2-(2,4-dichlorophenoxy)acetic acid (MPI55a) as a white solid following a general procedure **A** (yield 54%). ^1^H NMR (400 MHz, Chloroform-d) δ 7.29 (d, J = 2.5 Hz, 1H), 7.14 – 7.02 (m, 1H), 6.79 (dd, J = 8.9, 7.5 Hz, 1H), 4.64 – 4.41 (m, 2H), 4.37 (s, 1H), 3.82 (dd, J = 10.5, 5.3 Hz, 1H), 3.67 (d, J = 1.5 Hz, 1H), 3.64 (d, J = 4.3 Hz, 3H), 1.45 (dd, J = 7.4, 5.1 Hz, 1H), 1.37 (d, J = 7.5 Hz, 1H), 0.98 (d, J = 4.1 Hz, 3H), 0.84 (s, 2H), 0.76 (s, 1H). 13C NMR (100MHz, Chloroform-d) δ 171.82, 171.57, 166.18, 165.75, 152.24, 130.15, 130.10, 127.78, 127.67, 126.72, 123.56, 123.47, 114.52, 114.32, 69.33, 68.48, 59.91, 59.03, 52.75, 52.44, 47.35, 46.25, 32.28, 29.89, 27.61, 26.18, 24.69, 19.53, 19.45, 12.52, 12.33.

### Synthesis of (1R,2S,5S)-3-(2-(2,4-dichlorophenoxy)acetyl)-6,6-dimethyl-3-azabicyclo[3.1.0]hexane-2-carboxylic acid (MPI55d)

**MPI55d** was prepared as a white solid following a general procedure **B**. ^1^H NMR (400 MHz, Methanol-d4) δ 7.30 (dd, J = 4.3, 2.6 Hz, 1H), 7.10 (dd, J = 8.9, 2.6 Hz, 1H), 6.83 (dd, J = 8.9, 6.9 Hz, 1H), 4.82 – 4.69 (m, 2H), 4.24 (s, 1H), 3.79 (dd, J = 10.6, 5.3 Hz, 1H), 3.65 – 3.48 (m, 1H), 1.49 (dd, J = 7.5, 5.2 Hz, 1H), 1.45 – 1.35 (m, 1H), 0.97 (d, J = 3.6 Hz, 3H), 0.87 (s, 2H), 0.81 (s, 1H). ^13^C NMR (100 MHz, Methanol-d_4_) δ 173.04, 166.78, 129.44, 127.41, 126.07, 123.24, 114.71, 67.18, 59.79, 48.29, 48.08, 47.87, 47.65, 47.44, 47.23, 47.01, 45.66, 29.95, 27.21, 25.07, 19.10, 11.54.

### Methyl (S)-2-((1R,2S,5S)-3-(2-(2,4-dichlorophenoxy)acetyl)-6,6-dimethyl-3-azabicyclo[3.1.0]hexane-2-carboxamido)-3-((S)-2-oxopyrrolidin-3-yl)propanoate (MPI55e)

**MPI55e** was prepared with Int.i and (MPI55d) as a white gummy solid following general procedure **C.** ^1^H NMR (400 MHz, Chloroform-d) δ 7.27 (dd, J = 4.6, 2.5 Hz, 1H), 7.13 – 6.97 (m, 1H), 6.79 (dd, J = 22.5, 8.9 Hz, 1H), 4.65 – 4.55 (m, 2H), 4.51 – 4.39 (m, 1H), 4.34 – 4.28 (m, 1H), 3.87 – 3.68 (m, 2H), 3.65 (d, J = 11.1 Hz, 3H), 3.35 – 3.05 (m, 2H), 2.52 – 2.35 (m, 1H), 2.35 – 2.23 (m, 1H), 2.09 – 1.95 (m, 1H), 1.95 – 1.71 (m, 3H), 1.65 – 1.48 (m, 3H), 1.39 – 1.27 (m, 1H), 0.98 (d, J = 4.5 Hz, 3H), 0.81 (d, J = 10.9 Hz, 3H).

### (1R,2S,5S)-3-(2-(2,4-dichlorophenoxy)acetyl)-N-((S)-1-hydroxy-3-((S)-2-oxopyrrolidin-3-yl)propan-2-yl)-6,6-dimethyl-3-azabicyclo[3.1.0]hexane-2-carboxamide (MPI55f)

MPI55f was prepared as a white solid following a general procedure **D** (yield 63%). ^1^H NMR (400 MHz, Chloroform-d) δ 7.34 – 7.24 (m, 1H), 7.15 – 7.02 (m, 1H), 6.88 – 6.60 (m, 1H), 4.64 – 4.51 (m, 2H), 4.05 (dd, J = 5.1, 3.9 Hz, 1H), 4.00 – 3.76 (m, 3H), 3.70 – 3.43 (m, 5H), 3.32 – 3.08 (m, 2H), 2.48 – 2.32 (m, 1H), 2.34 – 2.17 (m, 1H), 2.03 – 1.84 (m, 1H), 1.82 – 1.63 (m, 2H), 1.59 – 1.49 (m, 1H), 0.98 (s, 3H), 0.80 (d, J = 14.5 Hz, 3H). ^13^C NMR (100 MHz, Chloroform-d) δ 181.24, 171.79, 166.09, 152.43, 130.05, 127.70, 127.62, 126.60, 126.35, 123.61, 114.73, 114.64, 71.03, 68.13, 67.98, 65.19, 61.88, 61.15, 51.35, 46.57, 40.59, 38.21, 31.88, 30.78, 29.06, 27.61, 26.14, 25.62, 19.37, 12.66.

### Synthesis of (1R,2S,5S)-3-(2-(2,4-dichlorophenoxy)acetyl)-6,6-dimethyl-N-((S)-1-oxo-3-((S)-2-oxopyrrolidin-3-yl)propan-2-yl)-3-azabicyclo[3.1.0]hexane-2-carboxamide (MPI55)

**MPI55** was prepared as a white solid following a general procedure **E** (yield 56%). ^1^H NMR (400 MHz, Chloroform-d) δ 9.40 (dd, J = 5.3, 1.0 Hz, 1H), 7.27 (t, J = 2.7 Hz, 1H), 7.10 – 7.00 (m, 1H), 6.80 (dd, J = 8.9, 6.3 Hz, 1H), 4.64 (s, 2H), 4.22 – 4.09 (m, 1H), 3.95 – 3.77 (m, 1H), 3.59 (d, J = 10.4 Hz, 1H), 3.35 – 3.18 (m, 2H), 2.49 – 2.36 (m, 1H), 2.36 – 2.19 (m, 1H), 2.00 – 1.63 (m, 3H), 1.55 – 1.37 (m, 6H), 0.98 (d, J = 2.0 Hz, 3H), 0.82 (d, J = 14.1 Hz, 3H).

### (2S,4S)-1-((benzyloxy)carbonyl)-4-cyclohexylpyrrolidine-2-carboxylic acid (MPI56d)

**MPI56d** was prepared as a white solid following a general procedure **G**. ^1^H NMR (400 MHz, Chloroform-d) δ 7.43 – 7.28 (m, 5H), 5.37 – 5.04 (m, 2H), 4.46 (d, J = 8.9 Hz, 1H), 3.75 – 3.60 (m, 1H), 3.13 – 2.98 (m, 1H), 2.42 (dd, J = 12.8, 6.2 Hz, 1H), 2.05 (d, J = 7.4 Hz, 1H), 1.80 – 1.54 (m, 6H), 1.16 (dq, J = 16.7, 5.9 Hz, 5H), 1.03 – 0.82 (m, 2H).

### (2S,4S)-benzyl 4-cyclohexyl-2-(((S)-1-methoxy-1-oxo-3-((S)-2-oxopyrrolidin-3-yl)propan-2-yl)carbamoyl)pyrrolidine-1-carboxylate (MPI56e)

**MPI56e** was prepared with **Int.i** and MPI56d as a white gummy solid following general procedure **C** (yield 60%). ^1^H NMR (400 MHz, Chloroform-d) δ 7.74 (s, 0H), 7.54 (d, J = 7.2 Hz, 1H), 7.42 – 7.27 (m, 5H), 6.04 (dd, J = 54.4, 22.8 Hz, 1H), 5.14 (s, 2H), 4.55 (s, 1H), 4.47 – 4.32 (m, 1H), 3.85 – 3.58 (m, 5H), 3.31 – 3.21 (m, 2H), 2.52 – 2.01 (m, 5H), 1.90 – 1.53 (m, 9H), 1.26 – 1.06 (m, 5H), 1.03 – 0.85 (m, 2H).

### (2S,4S)-benzyl 4-cyclohexyl-2-(((S)-1-hydroxy-3-((S)-2-oxopyrrolidin-3-yl)propan-2-yl)carbamoyl)pyrrolidine-1-carboxylate (MPI56f)

**MPI56f** was prepared as a white solid following a general procedure **D** (yield 59%). ^1^H NMR (400 MHz, Chloroform-d) δ 7.80 (t, J = 8.4 Hz, 1H), 7.36 – 7.22 (m, 5H), 5.06 (d, J = 9.6 Hz, 2H), 4.39 – 4.16 (m, 1H), 4.00 – 3.81 (m, 1H), 3.74 (dd, J = 10.2, 7.7 Hz, 1H), 3.49 – 3.31 (m, 2H), 3.25 – 3.13 (m, 2H), 3.07 – 2.91 (m, 2H), 2.51 – 2.22 (m, 1H), 2.11 – 1.99 (m, 3H), 1.94 – 1.73 (m, 2H), 1.71 – 1.56 (m, 7H), 1.47 (ddd, J = 14.7, 10.8, 3.6 Hz, 1H), 1.27 – 1.09 (m, 5H), 1.00 – 0.88 (m, 2H).

### (2S,4S)-benzyl 4-cyclohexyl-2-(((S)-1-oxo-3-((S)-2-oxopyrrolidin-3-yl)propan-2-yl)carbamoyl)pyrrolidine-1-carboxylate(MPI56)

**MPI56** was prepared as a white solid following a general procedure **E** (yield 65%). ^1^H NMR (400 MHz, Chloroform-d) δ 9.50 (s, 0.5H), 9.16 (s, 0.5H), 8.65 – 8.45 (m, 0.5H), 8.12 (d, J = 6.3 Hz, 0.5H), 7.43 – 7.27 (m, 5H), 6.39 (s, 0.5H), 6.11 (s, 0.5H), 5.32 – 5.23 (m, 0.5H), 5.14 (t, J = 3.7 Hz, 1H), 5.01 (d, J = 12.4 Hz, 0.5H), 4.50 – 4.27 (m, 2H), 4.13 (d, J = 6.7 Hz, 0.5H), 3.75 (tdd, J = 11.8, 9.0, 8.1, 4.5 Hz, 1H), 3.39 – 3.15 (m, 3.5H), 3.06 (dt, J = 29.1, 10.0 Hz, 1H), 2.53 – 2.16 (m, 3H), 2.16 – 1.54 (m, 11H), 1.28 – 1.06 (m, 5H), 1.00 – 0.83 (m, 2H). ^13^C NMR (101 MHz, CDCl_3_) δ 200.09, 199.79, 179.99, 179.88, 173.91, 173.24, 155.68, 154.81, 136.61, 128.47, 128.05, 127.93, 67.05, 61.05, 57.12, 55.08, 51.16, 50.80, 43.71, 41.71, 40.52, 37.99, 31.89, 31.41, 26.29, 26.02.

### Synthesis of 3-benzyl 2-methyl (1R,2S,5S)-6,6-dimethyl-3-azabicyclo[3.1.0]hexane-2,3-dicarboxylate (MPI57c)

To a solution of methyl (1R,2S,5S)-6,6-dimethyl-3-azabicyclo[3.1.0]hexane-2-carboxylate (300 mg, 1.46 mmol) in dichloromethane (20 mL) was added benzyl chloroformate (300 mg, 0.25 mL, 1.75 mmol) dropwise, cooled to 0°C, followed by the addition of DIPEA (566 mg, 0.79 mL, 4.38 mmol). The reaction was allowed to stir at RT for overnight. The product was extracted with ethyl acetate (50 mL) and washed with saturated NaHCO_3_ solution (2×20 mL), 1 M HCl solution (2×20 mL), and saturated brine solution (2×20 mL) sequentially. The organic layer was dried over anhydrous Na_2_SO_4_ and then concentrated on vacuo. The residue was then purified with flash chromatography (50-100% EtOAc in hexanes as the eluent) to afford **MPI57c** as white solid (380 mg, 77%). ^1^H NMR (400 MHz, Chloroform-d) δ 7.24 – 7.15 (m, 5H), 5.13 – 4.86 (m, 3H), 4.16 (d, J = 23.4 Hz, 1H), 3.65 (d, J = 14.5 Hz, 3H), 3.43 (dd, J = 13.9, 10.9 Hz, 1H), 1.32 (d, J = 4.5 Hz, 3H), 0.95 (s, 4H), 0.87 (d, J = 1.9 Hz, 4H). ^13^C NMR (100 MHz, Chloroform-d) δ 154.18, 153.60, 136.68, 136.58, 128.45, 128.39, 127.91, 127.63, 127.59, 66.96, 66.90, 59.87, 59.53, 52.31, 52.29, 52.16, 46.89, 46.34, 32.03, 31.05, 27.32, 26.48, 26.26, 26.25, 19.40, 19.36, 12.56.

### Synthesis of (1R,2S,5S)-3-((benzyloxy)carbonyl)-6,6-dimethyl-3-azabicyclo[3.1.0]hexane-2-carboxylic acid (MPI57d)

**MPI57d** was prepared as a white solid following a general procedure **B**. ^1^H NMR (400 MHz, Chloroform-d) δ 7.30 – 7.13 (m, 5H), 5.13 – 4.94 (m, 2H), 4.22 (s, 1H), 3.69 – 3.59 (m, 1H), 3.46 (dd, J = 15.5, 11.0 Hz, 1H), 1.46 (dd, J = 20.2, 7.4 Hz, 1H), 1.38 – 1.32 (m, 1H), 0.98 (d, J = 1.8 Hz, 3H), 0.89 (s, 3H). ^13^C NMR (100 MHz, Chloroform-d) δ 177.88, 176.93, 154.75, 153.80, 136.40, 128.52, 128.43, 128.06, 127.90, 127.67, 127.50, 67.37, 67.19, 59.86, 59.34, 46.46, 32.00, 30.86, 27.25, 26.43, 26.30, 26.27, 19.54, 19.43, 12.59.

### Synthesis of benzyl (1R,2S,5S)-2-((1-methoxy-1-oxo-3-(2-oxopyrrolidin-3-yl)propan-2-yl)carbamoyl)-6,6-dimethyl-3-azabicyclo[3.1.0]hexane-3-carboxylate (MPI57e)

**MPI56e** was prepared with **Int.i** and MPI56d as a white gummy solid following general procedure **C** (yield 82%). ^1^H NMR (400 MHz, Chloroform-d) δ 7.29 – 7.12 (m, 5H), 5.12 – 4.95 (m, 2H), 4.37 – 4.21 (m, 1H), 4.08 (d, J = 2.9 Hz, 1H), 3.79 – 3.61 (m, 3H), 3.61 – 3.38 (m, 3H), 3.30 – 3.05 (m, 2H), 2.48 – 1.89 (m, 4H), 1.88 – 1.60 (m, 2H), 1.53 – 1.26 (m, 3H), 0.96 (d, J = 1.7 Hz, 3H), 0.84 (d, J = 2.4 Hz, 3H). ^13^C NMR (100 MHz, Chloroform-d) δ 179.87, 179.85, 172.64, 172.30, 172.10, 162.58, 154.52, 153.99, 136.72, 136.65, 128.46, 128.33, 127.92, 127.71, 127.53, 127.49, 67.05, 66.99, 61.27, 52.47, 52.37, 52.09, 51.35, 47.35, 38.63, 38.56, 36.51, 33.26, 32.80, 31.45, 31.19, 28.67, 26.40, 19.29, 19.14, 12.68, 12.60.

### Synthesis of benzyl (1R,2S,5S)-2-((1-hydroxy-3-(2-oxopyrrolidin-3-yl)propan-2-yl)carbamoyl)-6,6-dimethyl-3-azabicyclo[3.1.0]hexane-3-carboxylate (MPI57f)

**MPI57f** was prepared as a white solid following a general procedure **D** (yield 61%). ^1^H NMR (400 MHz, Chloroform-d) δ 7.36 – 7.15 (m, 6H), 5.30 – 4.90 (m, 2H), 4.01 – 3.79 (m, 1H), 3.76 – 3.58 (m, 2H), 3.52 – 3.36 (m, 2H), 3.26 – 3.08 (m, 2H), 2.47 – 2.07 (m, 2H), 2.00 – 1.63 (m, 2H), 1.59 – 1.26 (m, 3H), 0.96 (s, 3H), 0.84 (d, J = 4.3 Hz, 3H). ^13^C NMR (100 MHz, Chloroform-d) δ 181.08, 180.89, 173.46, 172.64, 154.68, 154.11, 136.56, 128.50, 128.47, 128.01, 127.99, 127.74, 127.60, 67.17, 67.07, 66.08, 65.42, 61.90, 51.43, 50.89, 47.34, 46.90, 40.60, 38.38, 38.16, 33.09, 31.95, 31.84, 31.64, 28.82, 28.74, 27.40, 26.23, 26.15, 19.27, 19.22, 12.66, 12.56.

### Synthesis of benzyl (1R,2S,5S)-6,6-dimethyl-2-((1-oxo-3-(2-oxopyrrolidin-3-yl)propan-2-yl)carbamoyl)-3-azabicyclo[3.1.0]hexane-3-carboxylate (MPI57).)

**MPI57** was prepared as a white solid following a general procedure **E** (yield 58%). ^1^H NMR (400 MHz, Chloroform-d) δ 9.44 (d, J = 0.8 Hz, 0H), 9.05 (d, J = 1.5 Hz, 0H), 7.37 – 7.15 (m, 5H), 5.26 – 5.13 (m, 1H), 5.12 – 5.00 (m, 1H), 4.44 – 4.24 (m, 1H), 4.14 (d, J = 6.7 Hz, 1H), 4.08 – 3.95 (m, 1H), 3.74 – 3.66 (m, 1H), 3.56 – 3.44 (m, 1H), 3.30 – 3.21 (m, 2H), 2.47 – 2.09 (m, 2H), 2.00 – 1.85 (m, 1H), 1.85 – 1.60 (m, 3H), 1.52 – 1.16 (m, 3H), 0.97 (d, J = 2.7 Hz, 3H), 0.86 (s, 3H). ^13^C NMR (100 MHz, Chloroform-d) δ 200.16, 199.86, 180.09, 173.51, 172.89, 154.55, 153.94, 136.65, 136.58, 128.46, 128.45, 128.05, 128.00, 127.96, 127.59, 67.13, 67.05, 61.45, 61.40, 58.49, 57.93, 50.85, 47.26, 40.73, 40.59, 38.67, 38.09, 32.96, 31.51, 29.55, 29.08, 26.32, 26.24, 26.16, 19.33, 19.22, 12.63, 12.55.

### Synthesis of (2S,4R)-1-((benzyloxy)carbonyl)-4-(tert-butoxy)pyrrolidine-2-carboxylic acid (MPI58d)

**MPI58d** was prepared as a white solid following a general procedure **G**. ^1^H NMR (400 MHz, Chloroform-d) δ 7.33 – 7.16 (m, 5H), 5.18 – 4.92 (m, 2H), 4.45 – 4.34 (m, 1H), 4.27 – 4.16 (m, 1H), 3.71 – 3.60 (m, 1H), 3.35 – 3.15 (m, 1H), 2.21 – 1.97 (m, 2H), 1.16 (t, J = 7.2 Hz, 1H), 1.09 (d, J = 2.1 Hz, 9H). ^13^C NMR (100 MHz, Chloroform-d) δ 177.30, 176.38, 155.60, 154.47, 136.33, 136.30, 128.51, 128.40, 128.11, 127.89, 127.57, 74.30, 69.14, 68.48, 67.51, 67.27, 60.55, 57.92, 57.54, 53.82, 53.20, 38.49, 37.29, 28.23, 14.18.

### Synthesis of benzyl (2S,4R)-4-(tert-butoxy)-2-(((S)-1-methoxy-1-oxo-3-((S)-2-oxopyrrolidin-3-yl)propan-2-yl)carbamoyl)pyrrolidine-1-carboxylate--methane (MPI58e)

**MPI58e** was prepared with **Int.i** and MPI58d as a white gummy solid following general procedure **C** (yield 87%). ^1^H NMR (400 MHz, DMSO-d6) δ 8.77 (dd, J = 4.4, 1.5 Hz, 1H), 8.62 – 8.49 (m, 2H), 7.62 (d, J = 3.5 Hz, 1H), 7.52 (dd, J = 8.4, 4.4 Hz, 1H), 7.42 – 7.22 (m, 5H), 5.13 – 4.92 (m, 2H), 4.42 – 4.22 (m, 3H), 3.65 – 3.57 (m, 4H), 3.23 – 3.02 (m, 3H), 2.16 – 1.86 (m, 5H), 1.66 – 1.47 (m, 2H), 1.26 (dd, J = 6.9, 4.5 Hz, 4H), 1.13 (d, J = 3.9 Hz, 9H). ^13^C NMR (100 MHz, DMSO-d6) δ 178.57, 178.27, 172.80, 172.76, 172.72, 172.43, 162.78, 154.21, 151.50, 140.08, 137.46, 137.34, 135.10, 129.29, 128.84, 128.66, 128.23, 128.00, 127.82, 127.24, 121.15, 73.96, 73.93, 68.67, 66.29, 58.89, 58.49, 54.06, 52.42, 50.77, 42.31, 38.72, 38.36, 38.01, 37.93, 36.25, 32.88, 32.72, 31.24, 28.48, 27.45, 18.56, 17.20, 12.96.

### Synthesis of benzyl (2S,4R)-4-(tert-butoxy)-2-(((S)-1-hydroxy-3-((S)-2-oxopyrrolidin-3-yl)propan-2-yl)carbamoyl)pyrrolidine-1-carboxylate--methane (MPI58f)

**MPI58f** was prepared as a white solid following a general procedure **D**. Yield (59%). ^1^H NMR (400 MHz, Chloroform-d) δ 7.42 (d, J = 7.7 Hz, 1H), 7.26 (d, J = 3.7 Hz, 5H), 6.04 (d, J = 11.2 Hz, 1H), 5.19 – 4.94 (m, 2H), 4.36 – 4.15 (m, 2H), 3.86 (d, J = 55.4 Hz, 1H), 3.73 – 3.52 (m, 2H), 3.50 – 3.31 (m, 2H), 3.22 (dd, J = 10.3, 4.2 Hz, 3H), 2.43 (s, 1H), 2.28 (s, 1H), 2.18 – 1.85 (m, 4H), 1.72 (p, J = 9.6 Hz, 2H), 1.51 (dd, J = 18.9, 10.4 Hz, 1H), 1.46 – 1.32 (m, 1H), 1.10 (s, 10H). ^13^C NMR (100 MHz, Chloroform-d) δ 181.08, 172.94, 155.66, 154.88, 136.54, 128.49, 128.01, 127.73, 74.03, 69.41, 67.20, 65.38, 59.86, 53.61, 50.80, 40.55, 37.73, 31.86, 28.84, 28.27.

### Synthesis of benzyl (2S,4R)-4-(tert-butoxy)-2-(((S)-1-oxo-3-((S)-2-oxopyrrolidin-3-yl)propan-2-yl)carbamoyl)pyrrolidine-1-carboxylate--methane (MPI58)

**MPI58** was prepared as a white solid following a general procedure **E**. Yield (66%). ^1^H NMR (400 MHz, Chloroform-d) δ 9.57 (d, J = 25.9 Hz, 1H), 9.20 (s, 0H), 8.61 – 8.37 (m, 0H), 8.23 – 7.98 (m, 1H), 7.34 (dt, J = 12.7, 6.5 Hz, 6H), 6.13 – 5.81 (m, 1H), 5.34 – 5.01 (m, 2H), 4.52 – 4.22 (m, 3H), 3.78 (ddd, J = 17.3, 10.6, 6.0 Hz, 1H), 3.47 – 3.20 (m, 3H), 2.62 – 1.63 (m, 9H), 1.19 (s, 9H). ^13^C NMR (100 MHz, Chloroform-d) δ 199.67, 179.99, 173.03, δ 136.60, 128.47, 127.99, 127.72, 74.05, 69.47, 59.36, 37.45, 28.27.

### (S)-N-((S)-1-hydroxy-3-((S)-2-oxopyrrolidin-3-yl)propan-2-yl)-6-azaspiro[3.4]octane-7-carboxamide hydrogen chloride (MPI-59i-1)

To a stirred solution of **YR-B-101c** (200 mg, 0.506 mmol) in 1,4-Dioxane (2 mL) at 0 °C was added 4N HCl (1.26 mL, 5.06 mmol). Reaction mixture was stirred at rt for 3 h. After completion of reaction, solvent was concentrated in a vacuum. The residue was used in the next step without further purification. (150 mg). Crude product proceeded for next step without purification.

### benzyl (S)-7-(((S)-1-hydroxy-3-((S)-2-oxopyrrolidin-3-yl)propan-2-yl)carbamoyl)-6-azaspiro[3.4]octane-6-carboxylate (MPI59f)

MPI59f was prepared by using procedure of **MPI57c.** Yield (54%). ^1^H NMR (400 MHz, CDCl3) δ 7.36 (s, 5H), 5.67 (d, J = 28.6 Hz, 1H), 5.25 – 4.99 (m, 2H), 4.25 (dd, J = 8.1, 6.0 Hz, 1H), 4.11 – 3.70 (m, 2H), 3.53 (s, 2H), 3.42 (dd, J = 11.3, 5.0 Hz, 1H), 3.36 – 3.24 (m, 2H), 2.41 (s, 1H), 2.30 – 2.11 (m, 3H), 2.05 – 1.78 (m, 9H).

### Benzyl (S)-7-(((S)-1-oxo-3-((S)-2-oxopyrrolidin-3-yl)propan-2-yl)carbamoyl)-6-azaspiro[3.4]octane-6-carboxylate (MPI59) (diastereomers)

**MPI59** was prepared as a white solid following a general procedure **E**. Yield (67%). ^1^H NMR (400 MHz, CDCl3) δ 9.44 (s, 0.5H), 9.02 (s, 0.5H), 8.69 – 8.63 (m, 0.5H), 8.17 (d, J = 6.4 Hz, 0.5H), 7.33 – 7.14 (m, 5H), 6.47 – 6.42 (m, 0.5H), 6.06 – 6.00 (m, 0.5H), 5.34 – 4.86 (m, 2H), 4.28 (dd, J = 7.8, 5.9 Hz, 1.5H), 4.05 – 3.93 (m, 0.5H), 3.55 – 3.44 (m, 2H), 3.26 (p, J = 8.1 Hz, 2H), 2.57 – 2.04 (m, 8H), 1.95 – 1.60 (m, 9H).

### tert-butyl 3-(((S)-1-methoxy-1-oxo-3-((S)-2-oxopyrrolidin-3-yl)propan-2-yl)carbamoyl)-2-azaspiro[4.4]nonane-2-carboxylate (MPI60e)

**MPI60e** was prepared with **Int.i** and 2-(tert-butoxycarbonyl)-2-azaspiro[4.4]nonane-3-carboxylic acid as a white gummy solid following general procedure **C** (yield 69%). ^1^H NMR (400 MHz, DMSO) δ 8.41 (dd, J = 19.5, 7.8 Hz, 1H), 7.64 (d, J = 34.9 Hz, 1H), 4.35 – 4.18 (m, 1H), 4.13 (t, J = 7.8 Hz, 1H), 3.62 (s, 3H), 3.25 (d, J = 10.8 Hz, 1H), 3.12 (td, J = 19.7, 9.1 Hz, 3H), 2.31 – 1.93 (m, 4H), 1.80 – 1.50 (m, 9H), 1.50 – 1.25 (m, 11H).

### benzyl 3-(((S)-1-methoxy-1-oxo-3-((S)-2-oxopyrrolidin-3-yl)propan-2-yl)carbamoyl)-2-azaspiro[4.4]nonane-2-carboxylate (MPI60e-1)

**MPI60e-1** was prepared with as a white gummy solid following general procedures **F** and **G** (yield 77%). ^1^H NMR (400 MHz, DMSO) δ 8.50 (dd, J = 13.0, 7.8 Hz, 1H), 7.61 (s, 1H), 7.41 – 7.23 (m, 5H), 5.11 – 4.91 (m, 2H), 4.40 – 4.18 (m, 2H), 3.61 (d, J = 12.8 Hz, 3H), 3.37 (d, J = 10.2 Hz, 2H), 3.22 (t, J = 10.0 Hz, 1H), 3.15 – 3.03 (m, 1H), 2.20 – 1.86 (m, 4H), 1.75 (td, J = 13.2, 7.8 Hz, 1H), 1.64 – 1.39 (m, 10H).

### Benzyl 3-(((S)-1-hydroxy-3-((S)-2-oxopyrrolidin-3-yl)propan-2-yl)carbamoyl)-2-azaspiro[4.4]nonane-2-carboxylate (MPI60f)

**MPI60f** was prepared as a white solid following a general procedure **D**. Yield (58%). ^1^H NMR (400 MHz, CDCl_3_) δ 7.79 (d, J = 6.9 Hz, 1H), 7.31 – 7.16 (m, 5H), 6.15 (d, J = 52.2 Hz, 1H), 5.16 – 4.91 (m, 2H), 4.20 (t, J = 7.8 Hz, 1H), 3.88 (d, J = 55.0 Hz, 1H), 3.63 – 3.12 (m, 6H), 2.44 – 2.03 (m, 3H), 1.94 (d, J = 7.6 Hz, 2H), 1.85 – 1.31 (m, 10H).

### Benzyl 3-(((S)-1-oxo-3-((S)-2-oxopyrrolidin-3-yl)propan-2-yl)carbamoyl)-2-azaspiro[4.4]nonane-2-carboxylate (MPI60)(1:1 diastereomers)

**MPI60** was prepared as a white solid following a general procedure **E**. Yield (60%). ^1^H NMR (400 MHz, CDCl3) δ 9.44 (s, 0.5H), 9.04 (s, 0.5H), 8.58 (s, 0.5H), 8.14 (s, 0.5H), 7.31 – 7.18 (m, 5H), 6.23 (s, 0.5H), 5.94 (s, 0.5H), 5.34 – 4.88 (m, H), 4.28 (t, J = 7.7 Hz, 1.5H), 4.03 – 3.97 (m, 0.5H), 3.57 – 3.34 (m, 1H), 3.31 – 3.23 (m, 3H), 2.58 – 2.27 (m, 1H), 2.23 – 2.07 (m, 1H), 1.94 – 1.36 (m, 13H).

### tert-butyl 3-(((S)-1-methoxy-1-oxo-3-((S)-2-oxopyrrolidin-3-yl)propan-2-yl)carbamoyl)-2-azaspiro[4.5]decane-2-carboxylate (MPI61e)

**MPI61e** was prepared with **Int.i** and 2-(tert-butoxycarbonyl)-2-azaspiro[4.5]decane-3-carboxylic acid a white gummy solid following general procedure **C** (yield 75%). ^1^H NMR (400 MHz, CDCl_3_) δ 6.38 (d, J = 141.5 Hz, 1H), 4.46 (d, J = 60.3 Hz, 1H), 4.18 (dd, J = 8.5, 7.3 Hz, 1H), 3.65 (d, J = 2.7 Hz, 3H), 3.33 – 3.21 (m, 2H), 3.14 – 2.99 (m, 1H), 2.48 – 2.27 (m, 2H), 2.11 (ddd, J = 13.2, 10.5, 4.8 Hz, 2H), 1.82 (dp, J = 11.5, 4.1 Hz, 3H), 1.35 (d, J = 18.7 Hz, 19H).

### Benzyl 3-(((S)-1-methoxy-1-oxo-3-((S)-2-oxopyrrolidin-3-yl)propan-2-yl)carbamoyl)-2-azaspiro[4.5]decane-2-carboxylate (MPI61e-1)

**MPI61e-1** was prepared with as a white gummy solid following general procedures **F** and **G** (yield 90%). ^1^H NMR (400 MHz, DMSO-d_6_) δ 8.57 (ddd, J = 20.3, 10.2, 7.2 Hz, 1H), 7.70 (dd, J = 12.9, 4.7 Hz, 1H), 7.49 – 7.30 (m, 6H), 5.14 (q, J = 6.3, 4.8 Hz, 1H), 5.13 – 4.98 (m, 1H), 4.46 – 4.23 (m, 2H), 3.73 – 3.65 (m, 2H), 3.63 (s, 1H), 3.56 – 3.47 (m, 1H), 3.31 – 3.17 (m, 1H), 3.20 – 3.13 (m, 1H), 3.15 – 2.89 (m, 1H), 2.17 (s, 2H), 2.28 – 2.02 (m, 1H), 1.75 – 1.57 (m, 2H), 1.66 (s, 2H), 1.51 (d, J = 17.1 Hz, 5H), 1.42 (d, J = 14.5 Hz, 8H). ^13^C NMR (101 MHz, DMSO) δ 22.04, 22.59, 24.93, 33.71, 34.57, 36.95, 37.61, 40.14, 40.99, 49.41, 51.28, 57.64, 65.17, 126.06, 126.64, 126.84, 127.55, 136.34, 153.33, 153.44, 171.66, 177.31.

### Benzyl 3-(((S)-1-hydroxy-3-((S)-2-oxopyrrolidin-3-yl)propan-2-yl)carbamoyl)-2-azaspiro[4.5]decane-2-carboxylate (MPI61f)

**MPI61f** was prepared as a white solid following a general procedure **D**. Yield (52%). ^1^H NMR (400 MHz, CDCl3) δ 7.33-7.19 (m, 5H), 6.40-6.06 (m, 1H), 5.16 – 4.87 (m, 2H), 4.29-4.13 (m, 1H), 3.98-3.75 (m, 1H), 3.65 – 3.05 (m, 6H), 2.48 – 2.21 (m, 1H), 2.21 – 1.60 (m, 6H), 1.48 – 1.16 (m, 10H).

### Benzyl 3-(((S)-1-oxo-3-((S)-2-oxopyrrolidin-3-yl)propan-2-yl)carbamoyl)-2-azaspiro[4.5]decane-2-carboxylate (MPI61)

**MPI61** was prepared as a white solid following a general procedure **E**. Yield (58%). ^1^H NMR (400 MHz, DMSO) δ 9.55 – 8.92 (m, 1H), 8.49 (dd, J = 12.5, 7.2 Hz, 1H), 7.56 (d, J = 10.1 Hz, 1H), 7.42 – 6.98 (m, 5H), 5.08 – 4.81 (m, 2H), 4.30 – 3.95 (m, 2H), 3.38 (t, J = 12.6 Hz, 1H), 3.14 – 2.76 (m, 3H), 2.32 – 0.96 (m, 17H).

### Synthesis of methyl (1R,2S,5S)-3-((4-chlorophenyl)glycyl)-6,6-dimethyl-3-azabicyclo[3.1.0]hexane-2-carboxylate (MPI62c)

**MPI62c** was prepared as a white solid following a general procedure **A**. Yield (69%). ^1^H NMR (400 MHz, Chloroform-d) δ 7.12 – 6.94 (m, 2H), 6.50 – 6.35 (m, 2H), 3.83 – 3.75 (m, 1H), 3.75 – 3.67 (m, 5H), 1.56 – 1.46 (m, 1H), 1.46 – 1.34 (m, 1H), 1.01 (d, J = 3.6 Hz, 3H), 0.87 (d, J = 14.6 Hz, 3H).

### Synthesis of (1R,2S,5S)-3-((4-chlorophenyl)glycyl)-6,6-dimethyl-3-azabicyclo[3.1.0]hexane-2-carboxylic acid (MPI62d)

To a stirred solution of 2 (300 mg, 0.1 mmol) in 1,4-dioxane (8 mL) was added a 4 M HCl solution in dioxane (8 mL). The reaction mixture was stirred at room temperature for 1 h and then concentrated in vacuo to get product **MPI62d**. ^1^H NMR (400 MHz, DMSO-d6) δ 7.07 (dd, J = 8.9, 2.4 Hz, 2H), 6.67 – 6.53 (m, 2H), 5.95 – 5.80 (m, 1H), 3.87 (d, J = 5.4 Hz, 1H), 3.63 (s, 2H), 1.63 – 1.53 (m, 1H), 1.41 (dd, J = 7.5, 3.4 Hz, 1H), 1.35 (d, J = 3.9 Hz, 1H), 1.27 – 1.15 (m, 1H), 1.03 (d, J = 2.0 Hz, 3H), 0.89 (d, J = 9.8 Hz, 3H).

### Synthesis of (1R,2S,5S)-3-((4-chlorophenyl)glycyl)-6,6-dimethyl-N-((S)-1-oxo-3-((S)-2-oxopyrrolidin-3-yl)propan-2-yl)-3-azabicyclo[3.1.0]hexane-2-carboxamide (MPI62)

**MPI62** was prepared with **MPI62d** and (S)-2-amino-3-((S)-2-oxopyrrolidin-3-yl)propanal a white solid following general procedure **C** (yield 35%).

### Synthesis of tert-butyl (S)-6-(((S)-1-methoxy-1-oxo-3-((S)-2-oxopyrrolidin-3-yl)propan-2-yl)carbamoyl)-5-azaspiro[2.4]heptane-5-carboxylate (MPI63c)

**MPI63c** was prepared with (S)-5-(tert-butoxycarbonyl)-5-azaspiro[2.4]heptane-6-carboxylic acid (MPI63a) and Int.i as a white solid following a general procedure **C** (yield 82%).

### Synthesis of methyl (S)-3-((S)-2-oxopyrrolidin-3-yl)-2-((S)-5-azaspiro[2.4]heptane-6-carboxamido)propanoate (MPI63i)

**MPI63i** was prepared as a white solid following a general procedure **F**.

### Synthesis of methyl (S)-2-((S)-5-(2-(2,4-dichlorophenoxy)acetyl)-5-azaspiro[2.4]heptane-6-carboxamido)-3-((S)-2-oxopyrrolidin-3-yl)propanoate (MPI63e)

**MPI63e** was prepared with 2-(2,4-dichlorophenoxy)acetic acid and Int.i as a white solid following a general procedure **C** (yield 44%). ^1^H NMR (400 MHz, Chloroform-d) δ 7.28 (dd, J = 5.4, 2.5 Hz, 1H), 6.86 (d, J = 8.9 Hz, 1H), 4.92 – 4.25 (m, 4H), 3.64 (d, J = 8.9 Hz, 3H), 3.32 – 3.13 (m, 2H), 2.52 – 2.35 (m, 1H), 2.35 – 2.19 (m, 1H), 2.19 – 1.92 (m, 2H), 1.92 – 1.63 (m, 4H), 1.48 – 1.24 (m, 2H), 0.70 – 0.52 (m, 3H), 0.52 – 0.36 (m, 1H).

### Synthesis of (S)-5-(2-(2,4-dichlorophenoxy)acetyl)-N-((S)-1-hydroxy-3-((S)-2-oxopyrrolidin-3-yl)propan-2-yl)-5-azaspiro[2.4]heptane-6-carboxamide (MPI63f)

**MPI63f** was prepared as a white solid following a general procedure **D**. (Yield 68%). ^1^H NMR (400 MHz, Chloroform-d) δ 7.91 (dd, J = 7.3, 3.5 Hz, 1H), 7.28 (t, J = 2.3 Hz, 1H), 7.14 – 7.00 (m, 1H), 6.87 (d, J = 8.8 Hz, 1H), 4.79 – 4.65 (m, 1H), 4.62 – 4.46 (m, 1H), 3.93 – 3.78 (m, 1H), 3.71 – 3.58 (m, 1H), 3.58 – 3.33 (m, 3H), 3.32 – 3.14 (m, 2H), 2.51 – 2.22 (m, 3H), 2.16 (dd, J = 12.8, 8.6 Hz, 1H), 2.01 – 1.91 (m, 1H), 1.91 – 1.81 (m, 1H), 1.81 – 1.71 (m, 1H), 1.55 (ddt, J = 19.3, 14.5, 4.6 Hz, 2H), 1.45 – 1.32 (m, 2H), 1.19 (d, J = 1.7 Hz, 1H), 0.64 – 0.43 (m, 4H).

### Synthesis of (S)-5-(2-(2,4-dichlorophenoxy)acetyl)-N-((S)-1-oxo-3-((S)-2-oxopyrrolidin-3-yl)propan-2-yl)-5-azaspiro[2.4]heptane-6-carboxamide (MPI63).)

**MPI63** was prepared as a white solid following a general procedure **E**. Yield 54%). ^1^H NMR (400 MHz, Chloroform-d) δ 9.44 – 9.32 (m, 1H), 8.29 – 8.14 (m, 1H), 7.99 – 7.76 (m, 2H), 7.66 (dt, J = 10.4, 7.4 Hz, 1H), 7.28 (dd, J = 6.6, 2.7 Hz, 1H), 7.13 – 7.01 (m, 1H), 7.01 – 6.77 (m, 2H), 4.80 – 4.70 (m, 2H), 4.70 – 4.63 (m, 1H), 4.25 – 4.04 (m, 1H), 3.70 – 3.49 (m, 2H), 3.43 (d, J = 9.7 Hz, 1H), 3.36 – 3.21 (m, 3H), 3.09 – 2.99 (m, 1H), 2.48 – 2.39 (m, 1H), 2.39 – 2.25 (m, 2H), 1.93 – 1.73 (m, 4H), 1.39 (dd, J = 19.1, 7.0 Hz, 2H), 0.80 (q, J = 7.2, 6.8 Hz, 2H), 0.68 – 0.37 (m, 5H).

### Methyl (1*R*,2*S*,5*S*)-3-(2-(cyclohexyloxy)acetyl)-6,6-dimethyl-3-azabicyclo[3.1.0]hexane-2-carboxylate (MPI64c)

**MPI64c** was prepared as a white solid following a general procedure **A**. Yield (67%). ^1^H NMR (400 MHz, DMSO) δ 4.21 – 3.81 (m, 3H), 3.77 – 3.63 (m, 4H), 3.58 – 3.44 (m, 1H), 3.31 – 3.19 (m, 1H), 1.90 – 1.74 (m, 2H), 1.72 – 1.59 (m, 2H), 1.58 – 1.51 (m, 1H), 1.50 – 1.37 (m, 2H), 1.20 (tq, *J* = 9.8, 3.1 Hz, 5H), 1.02 (s, 3H), 0.88 (d, *J* = 4.6 Hz, 3H).

### (1*R*,2*S*,5*S*)-3-(2-(cyclohexyloxy)acetyl)-6,6-dimethyl-3-azabicyclo[3.1.0]hexane-2-carboxylic acid (MPI64d)

**MPI64d** was prepared with as a white gummy solid following general procedure **B**. ^1^H NMR (400 MHz, CDCl_3_) δ 4.64-4,32(m, 1H), 4.09 – 3.88 (m, 2H), 3.80 – 3.52 (m, 2H), 3.34-3,15 (m, 1H), 1.90 – 1.76 (m, 2H), 1.64 (dd, *J* = 9.0, 6.2 Hz, 2H), 1.49 – 1.39 (m, 2H), 1.32 – 1.07 (m, 6H), 1.00 (s, 3H), 0.88 (s, 3H).

### Methyl (*S*)-2-((1*R*,2*S*,5*S*)-3-(2-(cyclohexyloxy)acetyl)-6,6-dimethyl-3-azabicyclo[3.1.0]hexane-2-carboxamido)-3-((*S*)-2-oxopyrrolidin-3-yl)propanoate (MPI64e)

MPI64e was prepared with MPI64d and Int.i as a white solid following a general procedure **C** (yield 75%). ^1^H NMR (400 MHz, CDCl_3_) δ 8.81 – 8.19 (m, 1H), 7.67-7.24 (m, 1H), 4.53 – 4.25 (m, 2H), 4.05 – 3.95 (m, 2H), 3.77 (dd, *J* = 10.6, 5.1 Hz, 1H), 3.72 – 3.68 (m, 1H), 3.66 (s, 3H), 3.33 – 3.20 (m, 3H), 2.39-2.30 (m, 1H), 2.17 – 2.07 (m, 1H), 1.88-1.80 (m, 3H), 1.49 – 1.43 (m, 2H), 1.31 – 1.07 (m, 9H), 0.98 (s, 3H), 0.85 (s, 3H).

### (1*R*,2*S*,5*S*)-3-(2-(cyclohexyloxy)acetyl)-6,6-dimethyl-*N*-((*S*)-1-oxo-3-((*S*)-2-oxopyrrolidin-3-yl)propan-2-yl)-3-azabicyclo[3.1.0]hexane-2-carboxamide (MPI64)

MPI64 was prepared as a white solid following a general procedures **D** & **E**. Yield (60%). ^1^H NMR (400 MHz, CDCl_3_) δ 9.44 (s, 1H), 8.12 (d, *J* = 5.9 Hz, 1H), 5.84 (s, 1H), 4.32 (s, 1H), 4.26 – 4.12 (m, 1H), 4.09 – 3.95 (m, 2H), 3.82 – 3.69 (m, 1H), 3.50 (d, *J* = 10.5 Hz, 1H), 3.34 – 3.25 (m, 3H), 2.52 – 2.31 (m, 2H), 1.95 – 1.82 (m, 4H), 1.50 – 1.40 (m, 3H), 1.31 – 1.11 (m, 6H), 0.98 (s, 3H), 0.86 (s, 3H).

### methyl (1R,2S,5S)-3-(3-cyclohexylpropanoyl)-6,6-dimethyl-3-azabicyclo[3.1.0]hexane-2-carboxylate (MPI65c)

**MPI65c** was prepared as a white solid following a general procedure **A**. Yield (88%). ^1^H NMR (400 MHz, CDCl_3_) δ 4.38 (s, 1H), 3.82 (dd, *J* = 10.1, 5.3 Hz, 1H), 3.75 (s, 3H), 3.48 (d, *J* = 10.1 Hz, 1H), 2.29 – 2.19 (m, 2H), 1.74 – 1.58 (m, 5H), 1.57 – 1.44 (m, 3H), 1.41 (d, *J* = 7.4 Hz, 1H), 1.31 – 1.09 (m, 5H), 1.05 (s, 3H), 0.95 (s, 3H), 0.92 – 0.81 (m, 2H).

### (1R,2S,5S)-3-(3-cyclohexylpropanoyl)-6,6-dimethyl-3-azabicyclo[3.1.0]hexane-2-carboxylic acid (MPI65d)

**MPI65d** was prepared as a white solid following a general procedure **B**.

### methyl (S)-2-((1R,2S,5S)-3-(3-cyclohexylpropanoyl)-6,6-dimethyl-3-azabicyclo[3.1.0]hexane-2-carboxamido)-3-((S)-2-oxopyrrolidin-3-yl)propanoate (MPI65e)

**MPI65e** was prepared with MPI65d and Int.i as a white solid following a general procedure **C** (yield 73%).^1^H NMR (400 MHz, CDCl_3_) δ 7.59 (d, *J* = 7.2 Hz, 1H), 5.92 (s, 1H), 4.58 (ddd, *J* = 11.0, 7.2, 4.2 Hz, 1H), 4.33 (s, 1H), 3.85 (dd, *J* = 10.3, 5.3 Hz, 1H), 3.75 (s, 3H), 3.54 – 3.47 (m, 1H), 3.44 – 3.32 (m, 2H), 2.44 (td, *J* = 7.9, 3.9 Hz, 1H), 2.31 – 2.11 (m, 3H), 2.02 – 1.81 (m, 2H), 1.69 (t, *J* = 9.8 Hz, 4H), 1.58 (d, *J* = 7.6 Hz, 1H), 1.32 – 1.11 (m, 4H), 1.07 (s, 3H), 0.95 (s, 3H), 0.88 (d, *J* = 11.7 Hz, 2H).

### (1R,2S,5S)-3-(3-cyclohexylpropanoyl)-N-((S)-1-hydroxy-3-((S)-2-oxopyrrolidin-3-yl)propan-2-yl)-6,6-dimethyl-3-azabicyclo[3.1.0]hexane-2-carboxamide (MPI65f)

**MPI65f** was prepared as a white solid following a general procedure **D**. (Yield 66%). ^1^H NMR (400 MHz, CDCl_3_) δ 7.56 (d, *J* = 7.2 Hz, 1H), 5.93 (d, *J* = 32.5 Hz, 1H), 4.25 (s, 1H), 3.98 (tt, *J* = 7.2, 4.0 Hz, 1H), 3.90 (dd, *J* = 10.3, 5.3 Hz, 1H), 3.76 (ddd, *J* = 11.6, 3.9, 2.2 Hz, 1H), 3.54 – 3.43 (m, 2H), 3.33 (dd, *J* = 9.2, 4.4 Hz, 2H), 2.55 – 2.47 (m, 1H), 2.44 – 2.35 (m, 1H), 2.24 (dq, *J* = 18.2, 7.5 Hz, 2H), 2.03 (ddd, *J* = 14.5, 10.7, 6.6 Hz, 1H), 1.89 – 1.74 (m, 1H), 1.70 – 1.59 (m, 6H), 1.53 – 1.44 (m, 4H), 1.18 (tdd, *J* = 20.8, 12.4, 9.4 Hz, 5H), 1.04 (d, *J* = 2.4 Hz, 3H), 0.91 (s, 3H), 0.88 – 0.78 (m, 2H).

### (1R,2S,5S)-3-(3-cyclohexylpropanoyl)-6,6-dimethyl-N-((S)-1-oxo-3-((S)-2-oxopyrrolidin-3-yl)propan-2-yl)-3-azabicyclo[3.1.0]hexane-2-carboxamide (MPI65)

**MPI65** was prepared as a white solid following a general procedure **E** (Yield 80%). ^1^H NMR (400 MHz, CDCl_3_) δ 9.45 (d, *J* = 0.8 Hz, 1H), 8.13 (d, *J* = 6.2 Hz, 1H), 6.09 (s, 1H), 4.29 (d, *J* = 9.5 Hz, 2H), 3.80 (dd, *J* = 10.3, 5.2 Hz, 1H), 3.42 (d, *J* = 10.3 Hz, 1H), 3.38 – 3.22 (m, 3H), 2.48 (ddd, *J* = 13.5, 6.5, 4.4 Hz, 1H), 2.33 (dddd, *J* = 12.1, 8.8, 6.0, 3.1 Hz, 1H), 2.27 – 2.04 (m, 3H), 2.01 – 1.69 (m, 3H), 1.67 – 1.51 (m, 4H), 1.49 – 1.38 (m, 4H), 1.21 – 1.05 (m, 4H), 0.99 (s, 3H), 0.87 (s, 3H), 0.79 (d, *J* = 10.8 Hz, 2H).

### tert-butyl 3-(((S)-1-amino-1-oxo-3-((S)-2-oxopyrrolidin-3-yl)propan-2-yl)carbamoyl)-2-azaspiro[4.4]nonane-2-carboxylate (MPI66-1h)

**MPI66-1h** was prepared with MPI66-1g and Int.ii as a white solid following a general procedure **C** (yield 73%).. ^1^H NMR (400 MHz, DMSO) δ 8.22 – 7.88 (m, 1H), 7.62 (d, J = 12.6 Hz, 1H), 7.24 (d, J = 44.3 Hz, 1H), 7.05 (s, 1H), 4.29 – 4.18 (m, 1H), 4.18 – 4.05 (m, 1H), 3.30 – 3.04 (m, 4H), 2.40 – 1.90 (m, 4H), 1.78 – 1.41 (m, 10H), 1.33 (t, J = 20.8 Hz, 9H).

### N-((S)-1-amino-1-oxo-3-((S)-2-oxopyrrolidin-3-yl) propan-2-yl)-2-azaspiro[4.4]nonane-3-carboxamide (MPI66-1i)

**MPI66-1i** was prepared as a white solid following a general procedure **F** (Yield 80%).

### Benzyl3-(((S)-1-amino-1-oxo-3-((S)-2-oxopyrrolidin-3-yl)propan-2-yl)carbamoyl)-2-azaspiro[4.4]nonane-2-carboxylate (MPI66-1k)

**MPI66-1k** was prepared as a white solid following a general procedure **G** (Yield 75%). ^1^H NMR (400 MHz, DMSO) δ 8.28 – 8.09 (m, 1H), 7.56 (t, J = 14.3 Hz, 1H), 7.41 – 7.17 (m, 5H), 7.03 (d, J = 16.2 Hz, 1H), 5.09 – 4.91 (m, 2H), 4.39 – 4.17 (m, 2H), 3.32 – 2.80 (m, 4H), 2.39 – 2.28 (m, 1H), 2.21 – 1.88 (m, 3H), 1.73 (dt, J = 19.2, 6.1 Hz, 1H), 1.65 – 1.34 (m, 10H).

### benzyl 3-(((S)-1-cyano-2-((S)-2-oxopyrrolidin-3-yl)ethyl)carbamoyl)-2-azaspiro[4.4]nonane-2-carboxylate (MPI66-1)

**MPI66-1** was prepared as a white solid following a general procedure **H** (Yield 52%). ^1^H NMR (400 MHz, DMSO) δ 8.98 – 8.75 (m, 1H), 7.70 (d, J = 3.9 Hz, 1H), 7.45-7.20 (m, 5H), 5.09 – 5.00 (m, 2H), 4.99-4.92 (m, 1H), 4.27 – 4.10 (m, 1H), 3.39 (d, J = 10.4 Hz, 1H), 3.25 (t, J = 10.4 Hz, 1H), 3.19 – 2.97 (m, 2H), 2.24 – 2.02 (m, 3H), 1.82 – 1.69 (m, 2H), 1.67 – 1.40 (m, 9H).

### 3-chlorobenzyl 3-(((S)-1-amino-1-oxo-3-((S)-2-oxopyrrolidin-3-yl)propan-2-yl)carbamoyl)-2-azaspiro[4.4]nonane-2-carboxylate (MPI66-2k)

To 3-chlorobenzyl alcohol (0.3 g, 2.104 mmol) in CH_3_CN (10 mL) were added DIPEA (1.1 mL, 6.311 mmol) and N,N’-disuccinimidyl carbonate (753 mg, 2.94 mmol) at 0 °C. After 10 h at rt, **MPI66-2i** (750 mg, 2.104 mmol) was added one portion at 0 °C. After 10 h at rt, the reaction mixture was evaporated in vacuo. Purification by silica gel chromatography (Dichloromethane/MeOH = 9:1). 350 mg of compound isolated. Yield 50%. ^1^H NMR (400 MHz, DMSO) δ 8.37 – 8.10 (m, 1H), 7.72 – 7.52 (m, 1H), 7.43 – 6.97 (m, 5H), 5.13 – 4.90 (m, 2H), 4.46 – 4.17 (m, 2H), 3.38 (t, *J* = 11.3 Hz, 1H), 3.20 (d, *J* = 10.7 Hz, 1H), 3.16 – 2.89 (m, 2H), 2.13 (ddd, *J* = 25.8, 12.5, 7.8 Hz, 2H), 1.84 – 1.24 (m, 13H).

### 3-chlorobenzyl 3-(((S)-1-cyano-2-((S)-2-oxopyrrolidin-3-yl)ethyl)carbamoyl)-2-azaspiro[4.4]nonane-2-carboxylate (MPI66-2)

**MPI66-2** was prepared as a white solid following a general procedure **H** (Yield 48%). ^1^H NMR (400 MHz, CDCl_3_) δ 8.59 (dd, *J* = 18.8, 6.1 Hz, 0.5H), 8.27 (dd, *J* = 75.9, 7.0 Hz, 0.5H), 7.29 – 7.10 (m, 3H), 6.57 – 6.07 (m, 1H), 5.13 – 4.88 (m, 2H), 4.82 – 4.54 (m, 1H), 4.31 – 4.09 (m, 1H), 3.50 – 3.35 (m, 1H), 3.35 – 3.06 (m, 3H), 2.57 – 1.64 (m, 7H), 1.63 – 1.26 (m, 8H).

### N-((S)-1-hydroxy-3-((S)-2-oxopyrrolidin-3-yl)propan-2-yl)-2-(4-methoxy-1H-indole-2-carbonyl)-2-azaspiro[4.4]nonane-3-carboxamide (MPI66-3f)

**MPI66-3f** was prepared by 4-methoxy-1H-indole-2-carboxylic acid (0.345 mmol, 66 mg) and N-((S)-1-hydroxy-3-((S)-2-oxopyrrolidin-3-yl)propan-2-yl)-2-azaspiro[4.4]nonane-3-carboxamide (0.345 mmol, 120 mg) in general procedure **C** (Yield 36%). ^1^H NMR (400 MHz, CDCl_3_) δ 10.95 (s, 0.5H), 10.14 (s, 0.5H), 7.65 (s, 1H), 7.19 – 6.83 (m, 3H), 6.46 (d, *J* = 7.7 Hz, 1H), 4.64 (d, *J* = 67.9 Hz, 1H), 4.19 – 4.06 (m, 1H), 3.95 (d, *J* = 5.8 Hz, 3H), 3.91 – 3.76 (m, 2H), 3.68 (d, *J* = 16.7 Hz, 1H), 3.51 (s, 1H), 3.14 (s, 1H), 2.93 (s, 1H), 2.58 – 2.31 (m, 2H), 2.27 – 1.91 (m, 3H), 1.80 – 1.23 (m, 10H).

### 2-(4-methoxy-1H-indole-2-carbonyl)-N-((S)-1-oxo-3-((S)-2-oxopyrrolidin-3-yl)propan-2-yl)-2-azaspiro[4.4]nonane-3-carboxamide (MPI66-3)

**MPI66-3** was prepared as a white solid following a general procedure **E** (Yield 48%). ^1^H NMR (400 MHz, CDCl_3_) δ 9.46 (s, 1H), 8.38 – 8.20 (m, 1H), 7.17 – 6.86 (m, 3H), 6.44 (d, *J* = 7.7 Hz, 1H), 5.57 (s, 0.5H), 5.10 (s, 0.5H), 4.77 (t, *J* = 7.7 Hz, 0.5H), 4.66 (t, *J* = 8.3 Hz, 0.5H), 4.31 (dd, *J* = 16.3, 9.3 Hz, 1H), 3.94 – 3.72 (m, 5H), 3.31 – 2.90 (m, 2H), 2.55 – 2.46 (m, 1H), 2.39 – 2.08 (m, 2H), 2.02 – 1.86 (m, 1H), 1.82 – 1.59 (m, 6H), 1.49 (d, *J* = 30.5 Hz, 5H).

### tert-butyl 3-(((S)-1-amino-1-oxo-3-((S)-2-oxopiperidin-3-yl)propan-2-yl)carbamoyl)-2-azaspiro[4.4]nonane-2-carboxylate (MPI66-4h)

**MPI66-4h** was prepared with MPI66-4g and Int.ii as a white solid following a general procedure **C** (yield 82%).. ^1^H NMR (400 MHz, CDCl_3_) δ 4.33 (s, 1H), 4.13 (d, *J* = 9.0 Hz, 1H), 3.24 (h, *J* = 11.3 Hz, 4H), 2.37 – 1.99 (m, 3H), 1.96 – 1.71 (m, 4H), 1.68 – 1.29 (m, 19H).

### N-((S)-1-amino-1-oxo-3-((S)-2-oxopiperidin-3-yl)propan-2-yl)-2-azaspiro[4.4]nonane-3-carboxamide hydrogen chloride (MPI66-4i)

**MPI66-4i** was prepared as a white solid following a general procedure **F** (150 mg).

### N-((S)-1-amino-1-oxo-3-((S)-2-oxopiperidin-3-yl)propan-2-yl)-2-(4-methoxy-1H-indole-2-carbonyl)-2-azaspiro[4.4]nonane-3-carboxamide (MPI66-4k)

**MPI66-4k** was prepared with MPI66-4i and **4-methoxy-1H-indole-2-carboxylic acid** as a white solid following a general procedure **C** (yield 39%).. ^1^H NMR (400 MHz, DMSO) δ 11.57 (s, 1H), 8.43 (d, *J* = 8.5 Hz, 1H), 7.59 (s, 1H), 7.26 (s, 1H), 7.22 – 7.09 (m, 2H), 7.04 (d, *J* = 8.3 Hz, 1H), 6.93 (s, 1H), 6.53 (d, *J* = 7.7 Hz, 1H), 4.55 (dd, *J* = 9.7, 7.0 Hz, 1H), 4.28 – 4.13 (m, 1H), 3.89 (s, 3H), 3.87 – 3.79 (m, 2H), 3.20 – 3.04 (m, 2H), 2.24 – 2.04 (m, 3H), 2.01 – 1.75 (m, 3H), 1.63 (td, *J* = 7.1, 4.2 Hz, 7H), 1.51 – 1.23 (m, 4H).

### N-((S)-1-cyano-2-((S)-2-oxopiperidin-3-yl)ethyl)-2-(4-methoxy-1H-indole-2-carbonyl)-2-azaspiro[4.4]nonane-3-carboxamide (MPI66-4):)

**MPI66-4** was prepared as a white solid following a general procedure **H** (Yield 59%). ^1^H NMR (400 MHz, DMSO) δ 11.57 (d, *J* = 2.3 Hz, 1H), 8.87 (d, *J* = 8.1 Hz, 1H), 7.53 (s, 1H), 7.12 (q, *J* = 8.0 Hz, 1H), 7.04 (d, *J* = 8.2 Hz, 1H), 6.94 (d, *J* = 2.3 Hz, 1H), 6.53 (d, *J* = 7.6 Hz, 1H), 5.05 (q, *J* = 7.9 Hz, 1H), 4.48 (dd, *J* = 9.2, 7.5 Hz, 1H), 3.90 (s, 3H), 3.82 (d, *J* = 5.3 Hz, 2H), 3.11 (qd, *J* = 7.3, 4.8 Hz, 2H), 2.35 – 2.22 (m, 2H), 2.13 (dd, *J* = 12.2, 7.4 Hz, 1H), 1.93 – 1.85 (m, 1H), 1.77 (dd, *J* = 9.3, 2.7 Hz, 3H), 1.67 – 1.55 (m, 7H), 1.49 – 1.33 (m, 3H).

### benzyl 3-(((S)-1-amino-1-oxo-3-((S)-2-oxopyrrolidin-3-yl)propan-2-yl)carbamoyl)-2-azaspiro[4.5]decane-2-carboxylate (MPI67k)

**MPI67k** was prepared as a white solid following a general procedure **G** (Yield 79%). ^1^H NMR (400 MHz, CDCl3) δ 8.46 (d, J = 6.0 Hz, 1H), 7.31 – 7.22 (m, 5H), 7.15 (s, 1H), 6.15 (s, 1H), 5.42 (d, J = 11.8 Hz, 1H), 5.15-5.00 (m, 2H), 4.25 – 4.20 (m, 1H), 3.72 – 3.55 (m, 1H), 3.48 (dd, J = 18.8, 10.0 Hz, 1H), 3.28 (d, J = 8.6 Hz, 2H), 3.16-3.04 (m, 1H), 2.14 (dd, J = 12.7, 8.0 Hz, 2H), 2.10 – 1.96 (m, 2H), 1.95-1.75 (m, 2H), 1.71 – 1.54 (m, 2H), 1.46 – 1.27 (m, 12H).

### benzyl 3-(((S)-1-cyano-2-((S)-2-oxopyrrolidin-3-yl)ethyl)carbamoyl)-2-azaspiro[4.5]decane-2-carboxylate (MPI67)

**MPI67** was prepared as a white solid following a general procedure **H** (Yield 62%). ^1^H NMR (400 MHz, DMSO) δ 8.80 (t, J = δ 7.8 Hz, 1H), 7.63 (s, 1H), 7.32 – 7.18 (m, 5H), 5.02 – 4.91 (m, 2H), 4.86 (dd, J = 15.2, 7.5 Hz, 1H), 4.12 (dt, J = 29.4, 8.1 Hz, 1H), 3.46 – 3.34 (m, 1H), 3.10 – 2.99 (m, 3H), 2.12 – 2.00 (m, 2H), 1.70 – 1.61 (m, 1H), 1.56 – 1.44 (m, 1H), 1.42-1.21 (m, 10H).

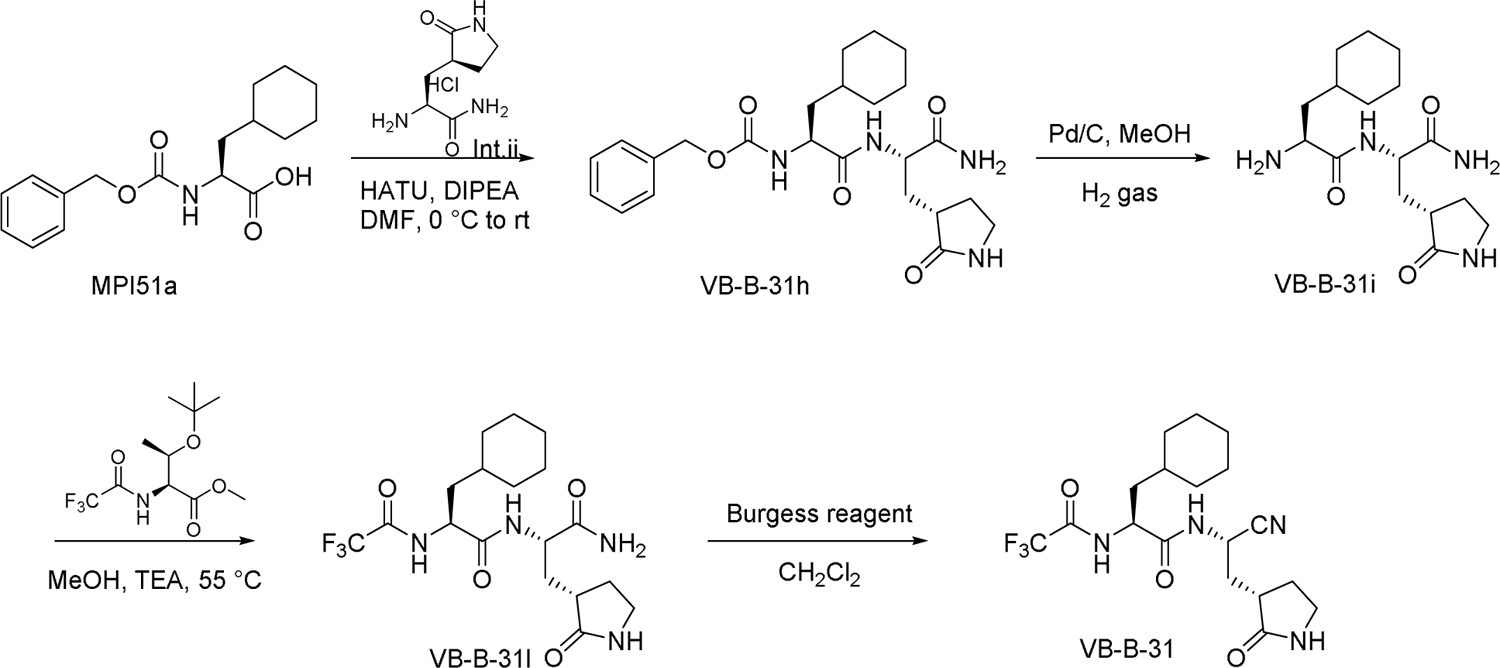

### Benzyl ((*S*)-1-(((*S*)-1-amino-1-oxo-3-((*S*)-2-oxopyrrolidin-3-yl)propan-2-yl)amino)-3-cyclohexyl-1-oxopropan-2-yl)carbamate (VB-B-31h):)

VB-B-31h was prepared with MPI51a and Int.ii as a white solid following a general procedure **C** (yield 82%).^1^H NMR (400 MHz, CDCl_3_) δ 8.24 (d, *J* = 6.8 Hz, 1H), 7.31 – 7.17 (m, 5H), 6.97 (s, 1H), 6.74 (s, 1H), 6.10 (s, 1H), 5.92 (d, *J* = 7.3 Hz, 1H), 5.08 – 4.93 (m, 2H), 4.38 (dt, *J* = 10.1, 6.2 Hz, 1H), 4.23 – 4.07 (m, 1H), 3.24-3.08 (m, 2H), 2.31-2.10 (m, 2H), 2.03 – 1.79 (m, 2H), 1.78 – 1.47 (m, 7H), 1.46-1.22 (m, 2H), 1.08 (p, *J* = 11.6 Hz, 3H), 0.93 – 0.68 (m, 2H). ^13^C NMR (101 MHz, CDCl_3_) δ 180.70, 174.41, 173.53, 156.64, 136.38, 128.53, 128.14, 127.96, 66.96, 53.58, 52.43, 40.73, 39.96, 38.49, 34.07, 33.75, 32.80, 32.29, 28.46, 26.22, 26.03.

### Methyl *O*-(*tert*-butyl)-*N*-(2,2,2-trifluoroethanethioyl)-*L*-threoninate (VB-B-31i)

To a solution of **VB-B-31h** (250 mg, 0.47 mmol) in methanol (10 mL) was added 10% Pd/C (50 mg). The reaction mixture was stirred under H_2_ balloon at room temperature for 3 h. The reaction mixture was filtered with celite, and the filtrate was concentrated *in vacuo* to yield **VB-B-31h** as colorless oil (168 mg, 90%), which was used without further purification. ^1^H NMR (400 MHz, CDCl_3_) δ 8.45 (s, 1H), 4.98 – 4.77 (m, 1H), 4.30 (qd, *J* = 6.3, 1.8 Hz, 1H), 3.69 (s, 3H), 1.15 (d, *J* = 6.4 Hz, 3H), 1.08 (s, 9H). ^13^C NMR (101 MHz, CDCl_3_) δ 184.85, 184.49, 168.63, 118.77, 115.99, 75.02, 67.14, 63.00, 52.71, 28.23, 21.43.

### (*S*)-*N*-((*S*)-1-amino-1-oxo-3-((*S*)-2-oxopyrrolidin-3-yl)propan-2-yl)-3-cyclohexyl-2-(2,2,2-trifluoroethanethioamido)propenamide (VB-B-31l)

methyl O-(tert-butyl)-N-(2,2,2-trifluoroacetyl)-L-threoninate (1.5 eq) and VB-B-31i (1 eq) in methanol was added TEA (2.5 eq) stirred at 55 °C for 48 h. Remove the solvent by rotavapor and work up with ethyl acetate. Purified by silics gel column chromatography. ^1^H NMR (400 MHz, DMSO) δ 8.52 – 8.28 (m, 1H), 7.67 (s, 1H), 7.36 (d, *J* = 5.5 Hz, 1H), 7.09 (s, 1H), 4.95 (dd, *J* = 11.1, 4.1 Hz, 1H), 4.30 (ddd, *J* = 11.8, 8.2, 4.1 Hz, 1H), 3.29 – 3.01 (m, 3H), 2.35 (t, *J* = 10.8 Hz, 1H), 2.14 – 1.94 (m, 2H), 1.74 – 1.66 (m, 6H), 1.59 – 1.51 (m, 1H), 1.30 (s, 1H), 1.25 – 1.11 (m, 5H), 1.00 – 0.88 (m, 2H).

### (*S*)-*N*-((*S*)-1-cyano-2-((*S*)-2-oxopyrrolidin-3-yl)ethyl)-3-cyclohexyl-2-(2,2,2-trifluoroethanethioamido)propenamide (VB-B-31)

VB-B-31 was prepared as a white solid following a general procedure **H** (Yield 62%). ^1^H NMR (400 MHz, CDCl_3_) δ 8.98 (d, *J* = 5.7 Hz, 1H), 6.23 (s, 1H), 4.98 – 4.79 (m, 1H), 4.66 (dt, *J* = 11.0, 5.3 Hz, 1H), 3.40 – 3.30 (m, 2H), 2.51 – 2.21 (m, 3H), 2.05 – 1.97 (m, 1H), 1.89 – 1.74 (m, 3H), 1.72 – 1.54 (m, 5H), 1.23 – 0.82 (m, 7H).

### tert-butyl (S)-7-(((S)-1-methoxy-1-oxo-3-((S)-2-oxopyrrolidin-3-yl)propan-2-yl)carbamoyl)-6-azaspiro[3.4]octane-6-carboxylate (YR-B-101e)

**YR-B-101e** was prepared by **(S)-6-(tert-butoxycarbonyl)-6-azaspiro[3.4]octane-7-carboxylic acid** and **Int.i** using general procedure **C**. (Yield 79%). ^1^H NMR (400 MHz, CDCl_3_) δ 4.51 (d, *J* = 54.1 Hz, 1H), 4.22 (t, *J* = 6.7 Hz, 1H), 3.72 (s, 3H), 3.40 (dd, *J* = 15.8, 8.3 Hz, 4H), 2.43 (s, 2H), 2.15 (dd, *J* = 17.5, 7.7 Hz, 3H), 1.99 – 1.77 (m, 8H), 1.44 (s, 9H).

### tert-butyl (S)-7-(((S)-1-hydroxy-3-((S)-2-oxopyrrolidin-3-yl)propan-2-yl)carbamoyl)-6-azaspiro[3.4]octane-6-carboxylate (Yr-B-101f)

**MPI101f** was prepared as a white solid following a general procedure **D** (Yield 62%). ^1^H NMR (400 MHz, CDCl_3_) δ 4.18 (ddd, *J* = 8.2, 5.6, 2.2 Hz, 1H), 4.02 (s, 1H), 3.80 – 3.23 (m, 6H), 2.37 (s, 2H), 2.24 – 1.73 (m, 11H), 1.44 (s, 9H).

### tert-butyl (S)-7-(((S)-1-oxo-3-((S)-2-oxopyrrolidin-3-yl)propan-2-yl)carbamoyl)-6-azaspiro[3.4]octane-6-carboxylate (YR-C-101)

**YR-B-101** was prepared as a white solid following a general procedure **H** (Yield 54%). (35 mg, 54%). ^1^H NMR (400 MHz, CDCl_3_) δ 9.51 (s, 1H), 4.28 (t, *J* = 6.8 Hz, 2H), 3.56 – 3.28 (m, 4H), 2.39 (s, 2H), 2.26 – 2.16 (m, 2H), 2.08 – 1.76 (m, 9H), 1.44 (s, 9H).

### (1R,2S,5S)-6,6-dimethyl-N-((S)-1-oxo-3-((S)-2-oxopyrrolidin-3-yl)propan-2-yl)-3-(2-(4-(trifluoromethoxy)phenoxy)acetyl)-3-azabicyclo[3.1.0]hexane-2-carboxamide (MI09)

**MI-09** was synthesized according to the literature. ^1^H NMR (400 MHz, Chloroform-*d*) δ 9.48 (s, 1H), 8.39 (d, *J* = 5.8 Hz, 1H), 7.17 – 7.04 (m, 2H), 6.90 (d, *J* = 9.2 Hz, 2H), 6.08 (s, 1H), 4.69 – 4.56 (m, 2H), 4.42 (s, 1H), 4.34 – 4.25 (m, 1H), 3.94 (dd, *J* = 10.3, 4.6 Hz, 1H), 3.56 (d, *J* = 10.3 Hz, 1H), 3.35 – 3.21 (m, 2H), 2.57 – 2.47 (m, 1H), 2.38 – 2.27 (m, 1H), 1.96 – 1.86 (m, 2H), 1.83 – 1.73 (m, 1H), 1.63 – 1.52 (m, 2H), 1.06 (s, 3H), 0.90 (s, 3H). ^13^C NMR (101 MHz, CDCl_3_): δ 199.91, 180.39, 172.15, 166.45, 156.47, 143.32, 143.30, 122.44, 115.62, 67.38, 61.35, 58.16, 46.42, 40.63, 38.26, 30.76, 29.61, 28.84, 27.60, 26.17, 19.42, 12.61.

### (1R,2S,5S)-3-(2-(2,4-dichlorophenoxy)acetyl)-6,6-dimethyl-N-((S)-1-oxo-3-((S)-2-oxopyrrolidin-3-yl)propan-2-yl)-3-azabicyclo[3.1.0]hexane-2-carboxamide (MI-14)

**MI-14** was synthesized according to the literature. ^1^H NMR (400 MHz, Chloroform-*d*) δ 9.40 (dd, *J* = 5.3, 1.0 Hz, 1H), 7.27 (t, *J* = 2.7 Hz, 1H), 7.10 – 7.00 (m, 1H), 6.80 (dd, *J* = 8.9, 6.3 Hz, 1H), 4.64 (s, 2H), 4.22 – 4.09 (m, 1H), 3.95 – 3.77 (m, 1H), 3.59 (d, *J* = 10.4 Hz, 1H), 3.35 – 3.18 (m, 2H), 2.49 – 2.36 (m, 1H), 2.36 – 2.19 (m, 1H), 2.00 – 1.63 (m, 3H), 1.55 – 1.37 (m, 6H), 0.98 (d, *J* = 2.0 Hz, 3H), 0.82 (d, *J* = 14.1 Hz, 3H).

### (1S,3aR,6aS)-2-(2-(2,4-dichlorophenoxy)acetyl)-N-((S)-1-oxo-3-((S)-2-oxopyrrolidin-3-yl)propan-2-yl)octahydrocyclopenta[c]pyrrole-1-carboxamide (MI30)

**MI-30** was synthesized according to the literature. ^1^H NMR (400 MHz, Chloroform-*d*) δ 9.41 (d, *J* = 0.9 Hz, 1H), 8.24 (d, *J* = 6.0 Hz, 1H), 7.29 (d, *J* = 2.5 Hz, 1H), 7.07 (dd, *J* = 8.8, 2.6 Hz, 1H), 6.81 (d, *J* = 8.9 Hz, 1H), 6.14 (s, 1H), 4.76 – 4.63 (m, 2H), 4.33 (d, *J* = 3.0 Hz, 1H), 4.21 (ddd, *J* = 8.9, 7.3, 5.9 Hz, 1H), 3.81 (dd, *J* = 10.6, 7.9 Hz, 1H), 3.42 (dd, *J* = 10.4, 4.0 Hz, 1H), 3.22 (ddd, *J* = 15.9, 9.6, 7.2 Hz, 2H), 2.85 – 2.67 (m, 2H), 2.48 – 2.37 (m, 1H), 2.30 – 2.22 (m, 1H), 1.99 – 1.90 (m, 1H), 1.87 – 1.78 (m, 3H), 1.74 – 1.62 (m, 2H), 1.60 – 1.46 (m, 2H), 1.43 – 1.34 (m, 1H). ^13^C NMR (101 MHz, CDCl): 199.85, 180.28, δ 172.70, 166.44, 152.46, 68.25, 67.08, 57.95, 52.66, 47.16, 43.25, 40.62, 38.19, 32.57, 31.99, 29.65, 28.79, 25.43.

### (1S,3aR,6aS)-2-(2-(3,4-dichlorophenoxy)acetyl)-N-((S)-1-oxo-3-((S)-2-oxopyrrolidin-3-yl)propan-2-yl)octahydrocyclopenta[c]pyrrole-1-carboxamide (MI-31)

**MI-31** was synthesized according to the literature. ^1^H NMR (400 MHz, Chloroform-d) δ 9.4 (s, 1H), δ8.4 (d, J = 5.7 Hz, 1H), 7.3 – 7.2 (m, 1H), 7.0 (d, J = 3.0 Hz, 1H), 6.8 (td, J = 8.8, 2.9 Hz, 1H), 6.2 (s, 1H), 4.6 (d, J = 4.0 Hz, 2H), 4.4 (d, J = 2.9 Hz, 1H), 4.3 – 4.1 (m, 1H), 3.8 (dd, J = 10.4, 8.0 Hz, 1H), 3.3 – 3.1 (m, 3H), 2.8 (dp, J = 12.2, 4.3, 3.9 Hz, 1H), 2.7 (tdd, J = 8.0, 5.6, 2.9 Hz, 1H), 2.4 (dt, J = 16.1, 8.0 Hz, 1H), 2.3 (ddd, J = 12.3, 6.5, 2.3 Hz, 1H), 2.0 – 1.9 (m, 1H), 1.8 (tt, J = 13.6, 6.4 Hz, 3H), 1.8 – 1.6 (m, 2H), 1.5 (ddd, J = 22.4, 12.8, 6.0 Hz, 2H), 1.4 (tt, J = 12.6, 5.6 Hz, 1H). ^13^C NMR (100 MHz, Chloroform-d) δ200.0, 180.3, 172.8, 166.6, 157.1, 132.8, 130.7, 124.8, 116.7, 114.9, 67.1, 67.0, 58.1, 55.0, 52.6, 47.3, 43.2, 40.6, 32.5, 32.0, 29.7, 28.8, 25.4.

## Associated Content

## Author Information

## Acknowledgement

This work was supported by Welch Foundation (grant A-1715), National Institutes of Health (grants R35GM145351 to W.R.L., R21AI164088 to S.X., and R21EB032983 to W.R.L.), Texas A&M X Grants, and the Texas A&M EDGE Fellowship Program.

