## supporting information for "A Systematic Survey of Reversibly Covalent Dipeptidyl Inhibitors of the SARS-CoV-2 Main Protease"

| Table S1. Data Collection and refinement statistics. | | |  |
| --- | --- | --- | --- |
|  | **MPI48 (7SD9)** | **MPI49 (7SDA)** | **MI-09 (7SDC)** |
| Resolution Range | 24.28  - 1.85 (1.916  - 1.85 | 24.32  - 1.85 (1.916  - 1.85) | 24.35 - 1.85 (1.916 - 1.85) |
| Space Group | I 1 2 1 | I 1 2 1 | I 1 2 1 |
| Unit Cell | 51.661 80.8574 89.6971 90 96.6634 90 | 51.6838 81.3006 89.2993 90 97.0934 90 | 54.3646 80.9874 87.8118 90 97.2942 90 |
| Unique Reflections | 29850 (2976) | 31237 (3071) | 31726 (3108) |
| Completeness (%) | 95.31 (95.05) | 99.37 (97.93) | 98.23 (96.40) |
| Wilson B-factor | 10.94 | 17.17 | 20.72 |
| Reflections used in refinement | 29849 (2975) | 31128 (3069) | 31696 (3104) |
| Reflections used for R-free | 1508 (161) | 1561 (165) | 1542 (151) |
| R-work | 0.2192 (0.3465) | 0.2124 (0.3151) | 0.2600 (0.4761) |
| R-free | 0.2483 (0.3864) | 0.2378 (0.3362) | 0.3006 (0.5238) |
| Number of non-hydrogen atoms | 2610 | 2629 | 2566 |
| macromolecules | 2360 | 2360 | 2360 |
| ligands | 31 | 33 | 36 |
| solvent | 219 | 236 | 170 |
| Protein Residues | 301 | 301 | 301 |
| RMS(bonds) | 0.010 | 0.010 | 0.016 |
| RMS(angles) | 1.21 | 1.22 | 1.59 |

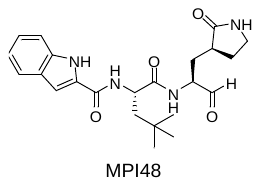

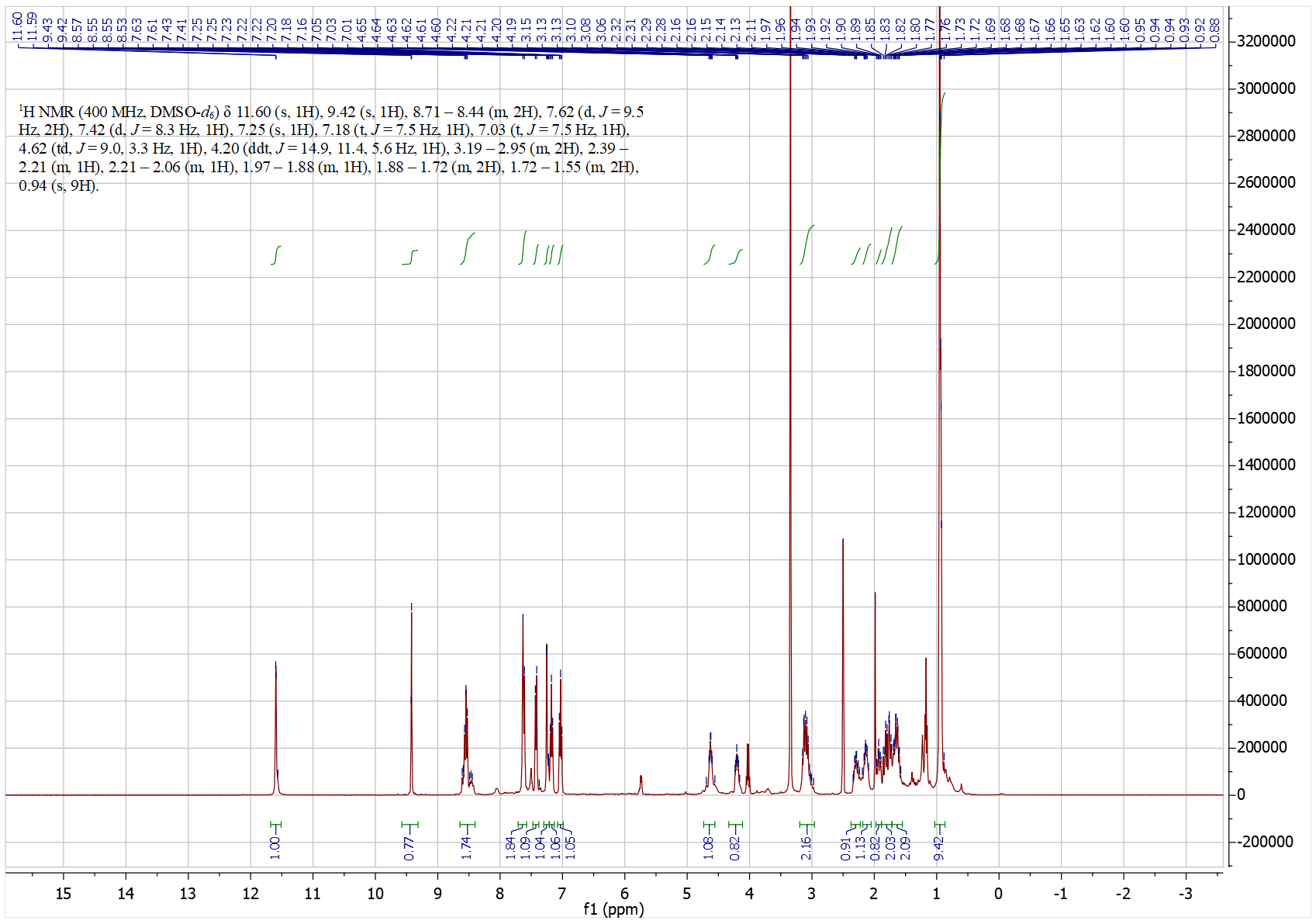

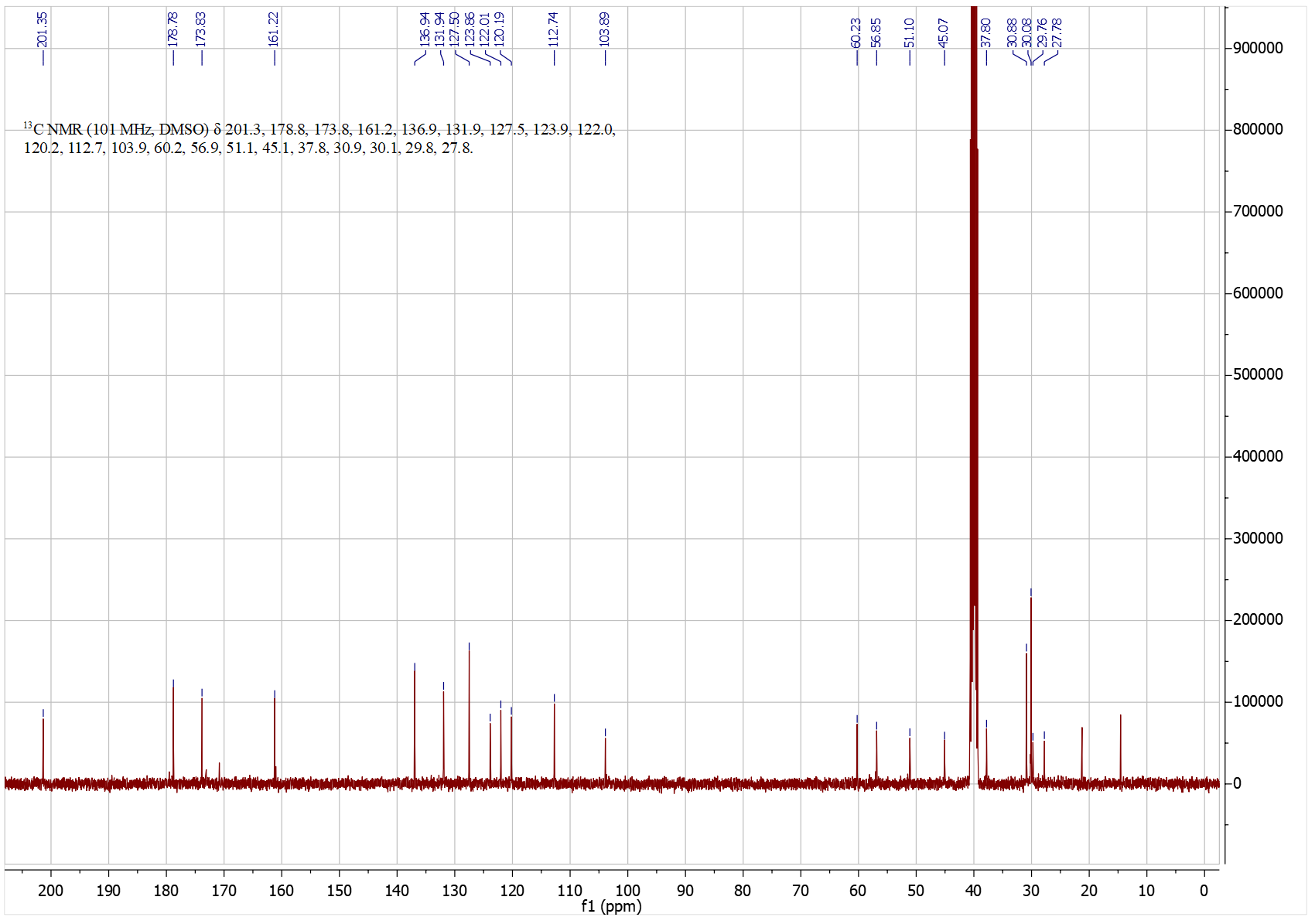

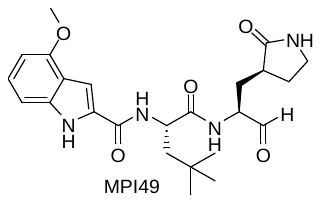

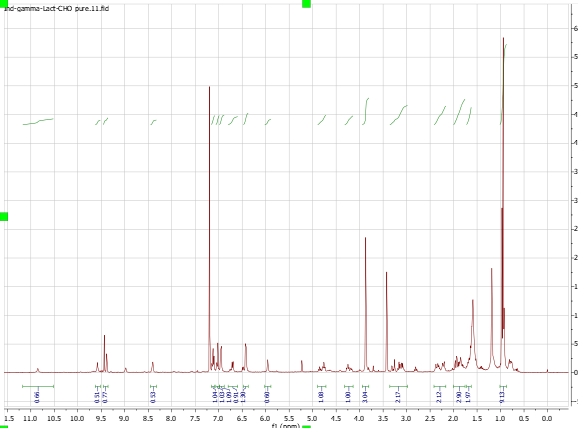

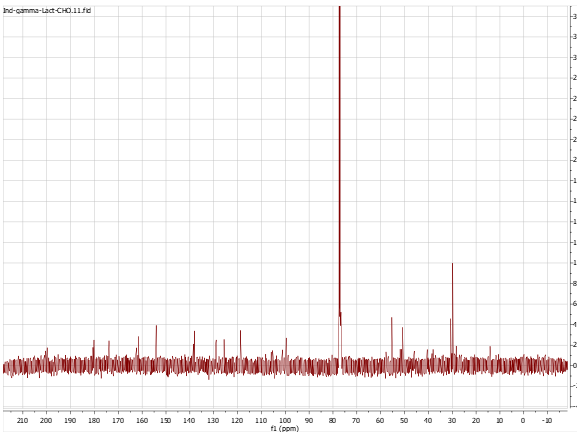

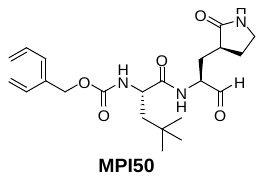

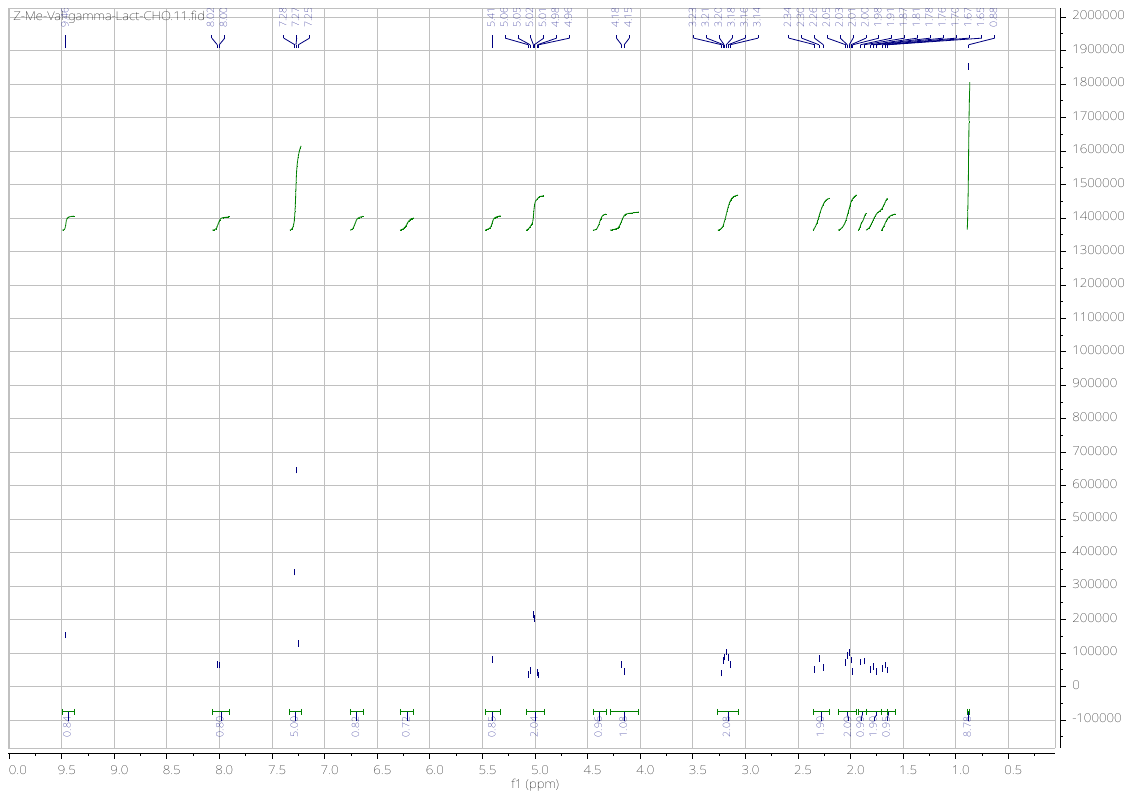

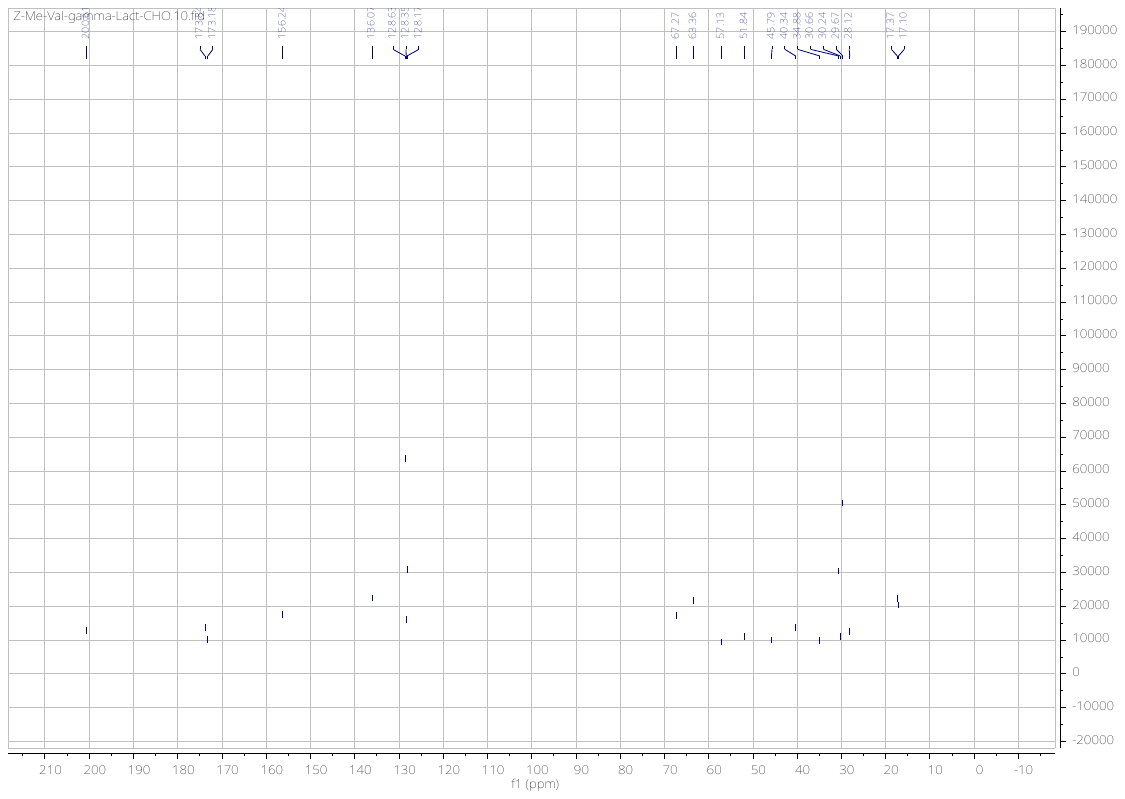

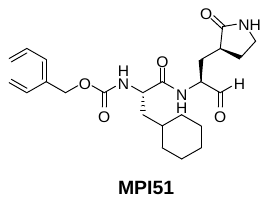

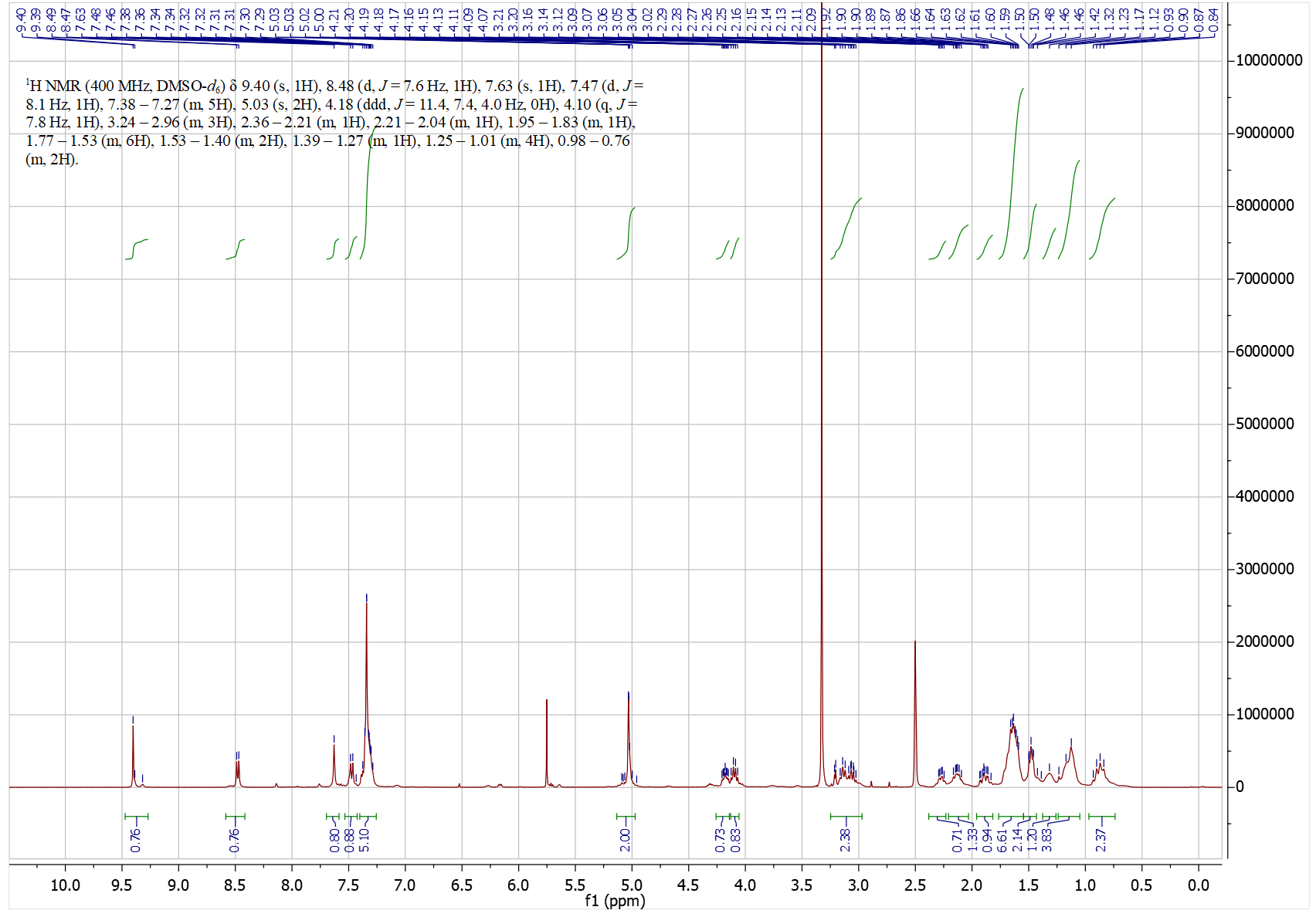

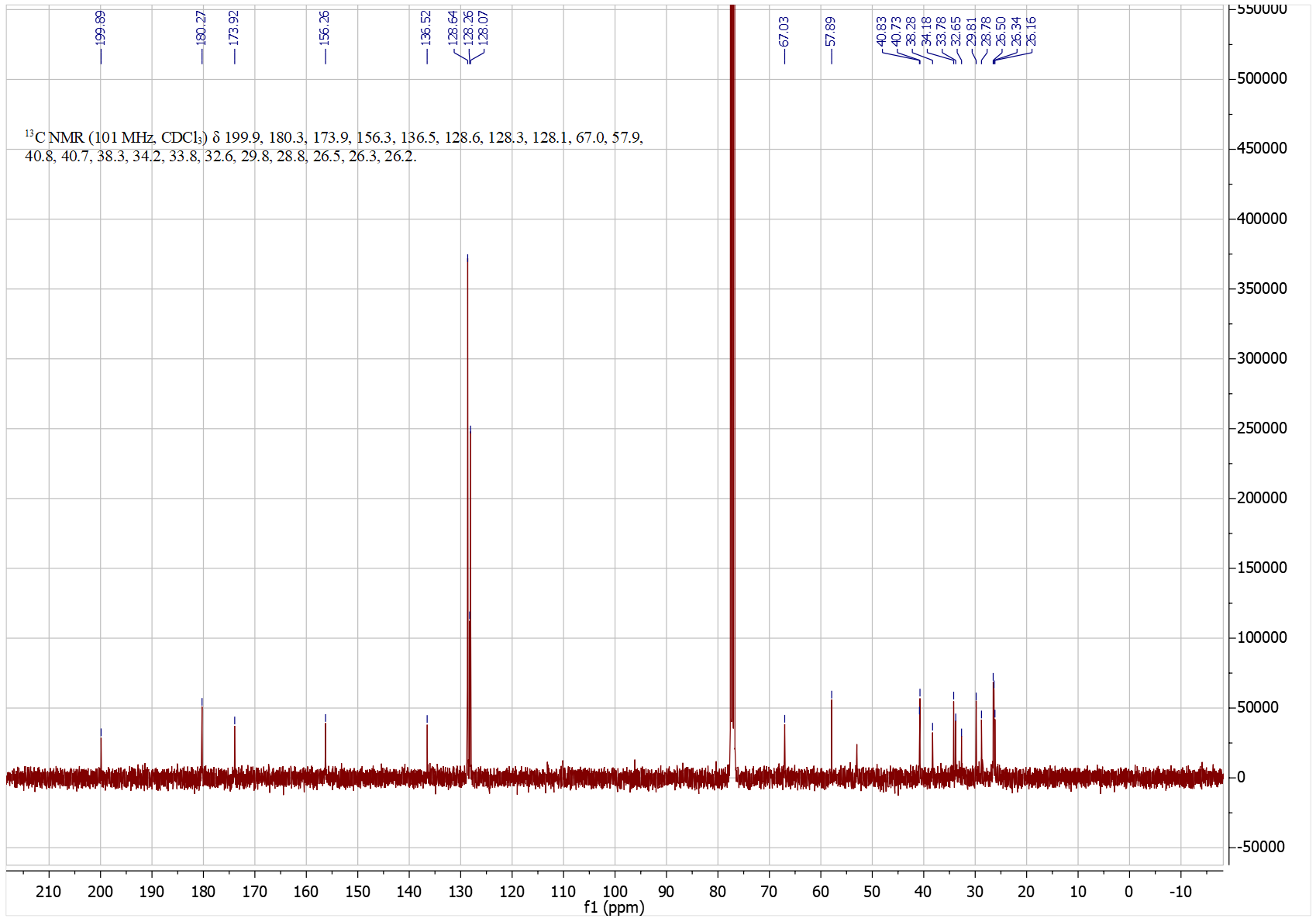

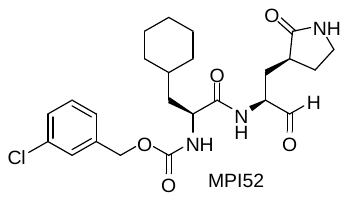

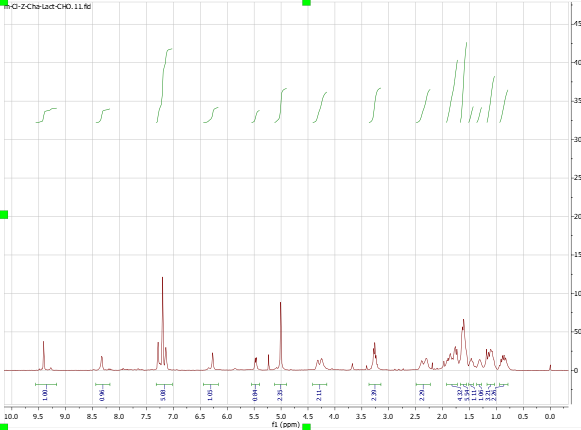

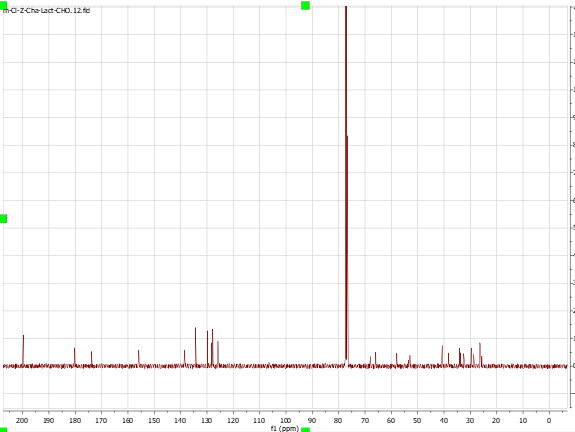

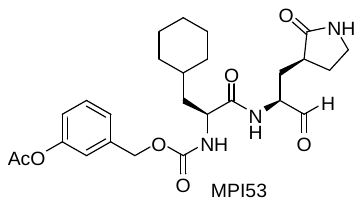

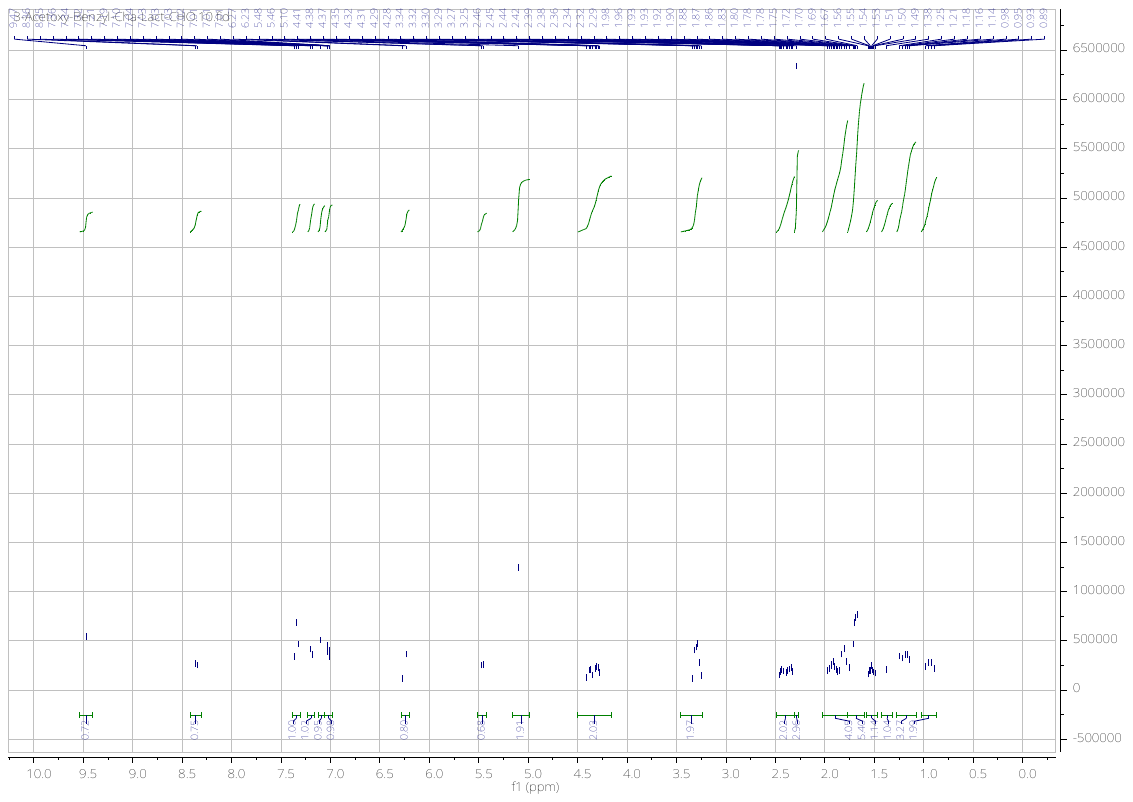

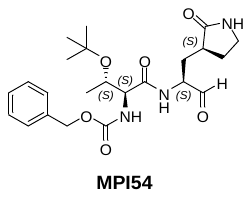

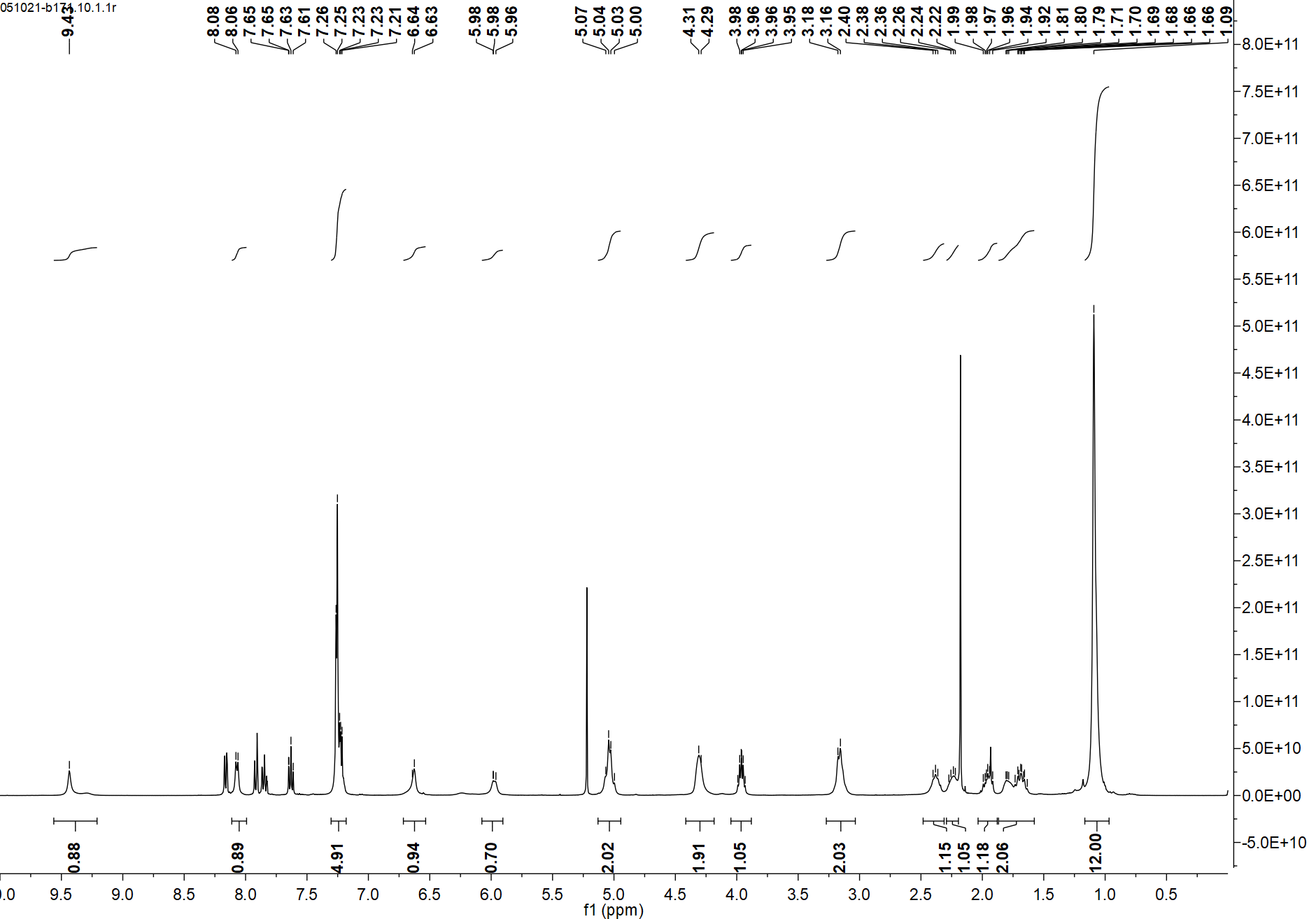

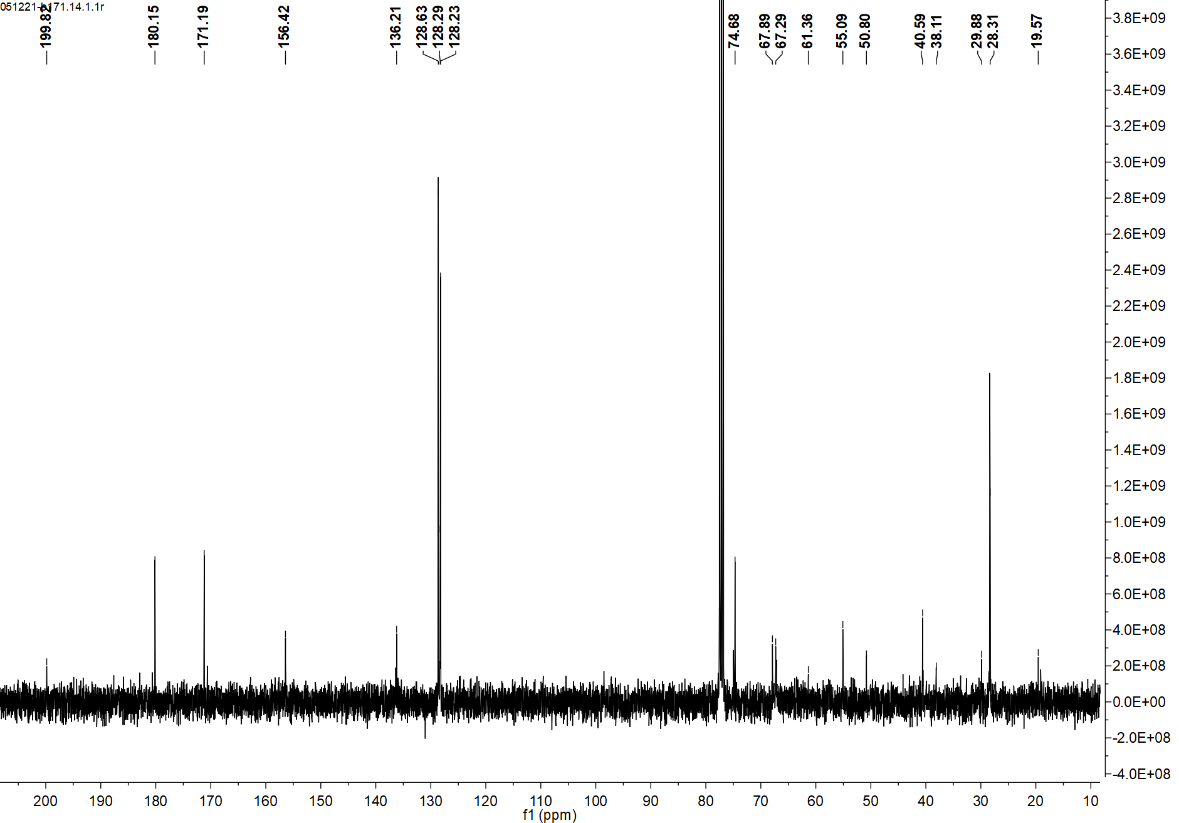

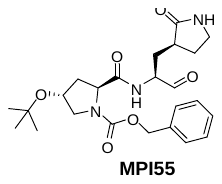

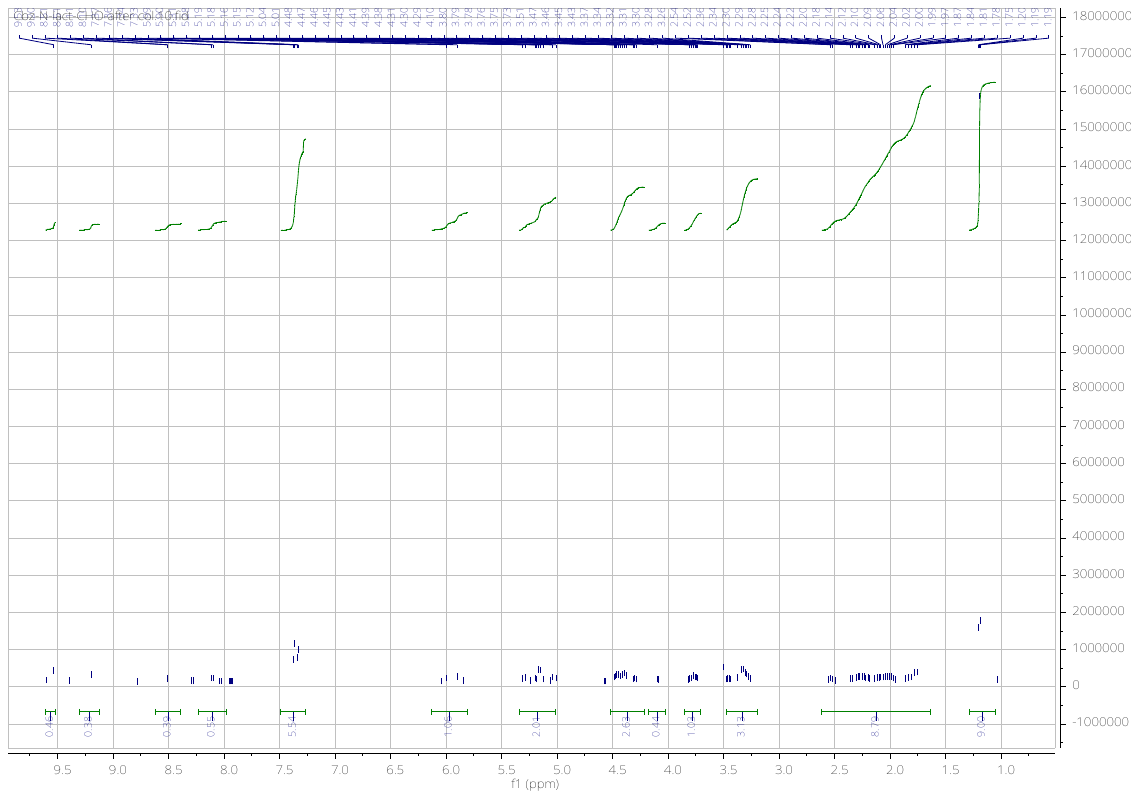

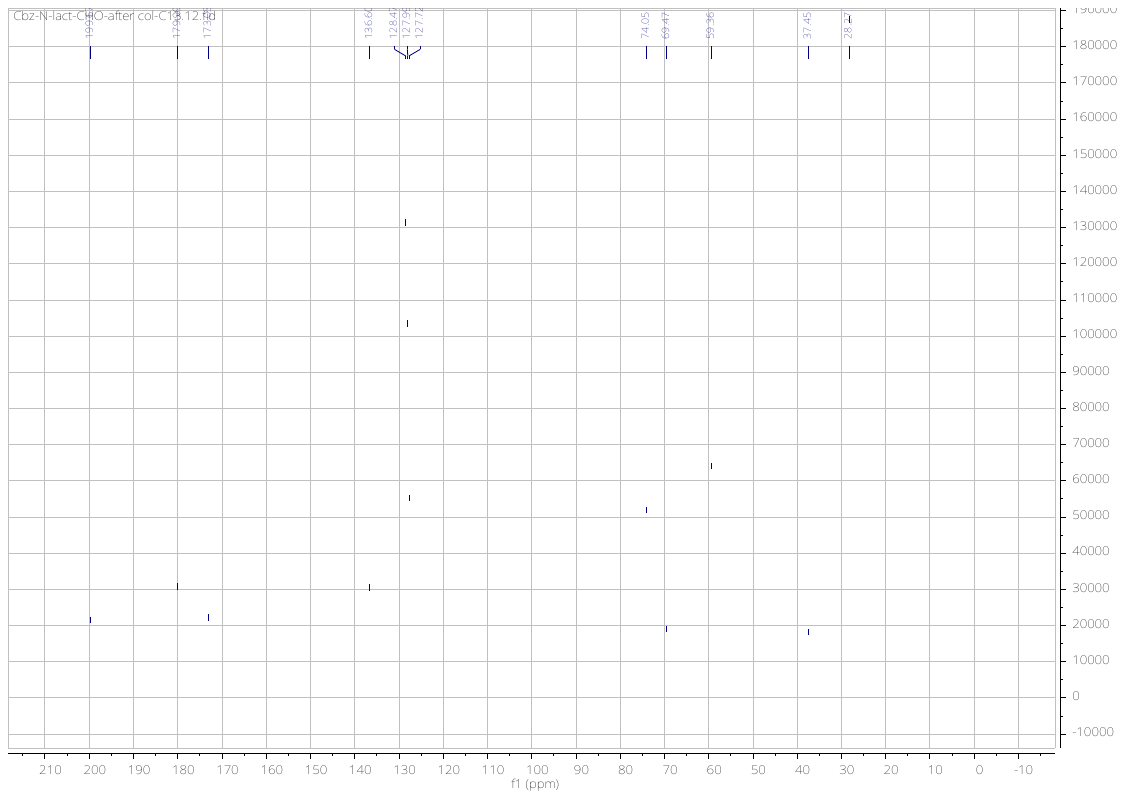

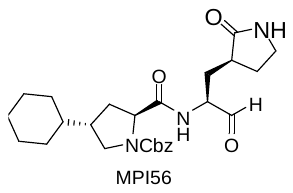

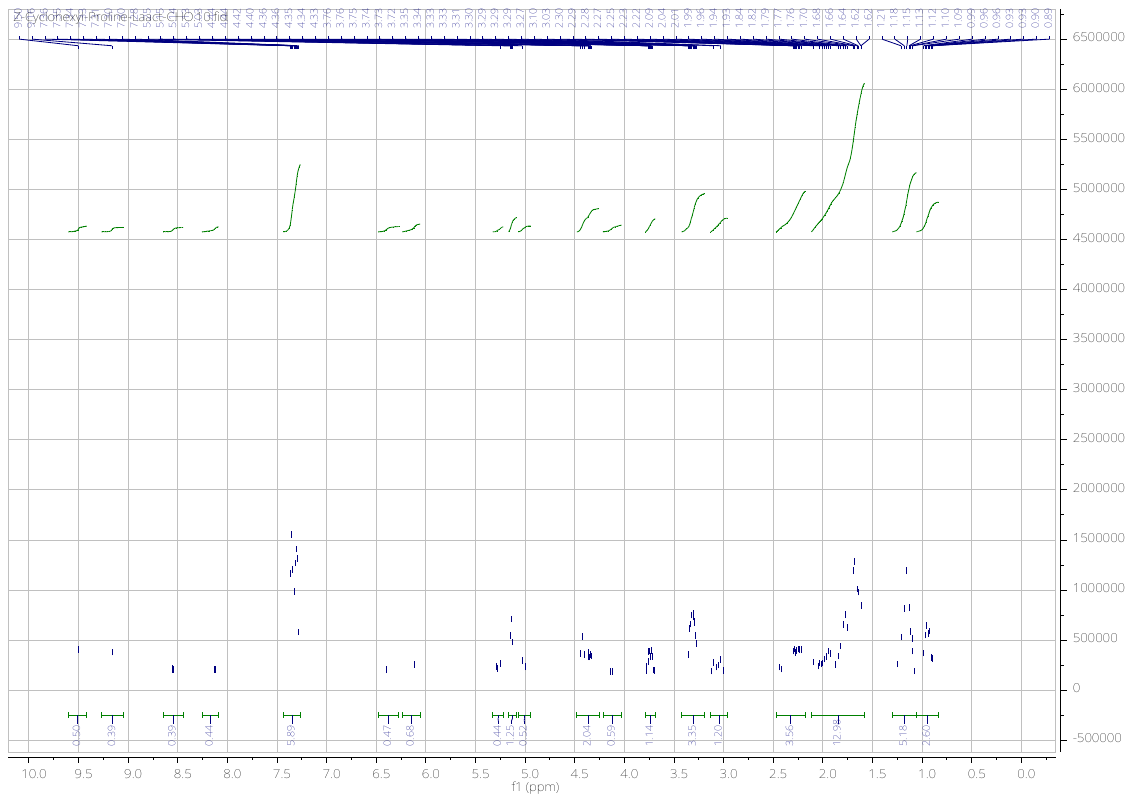

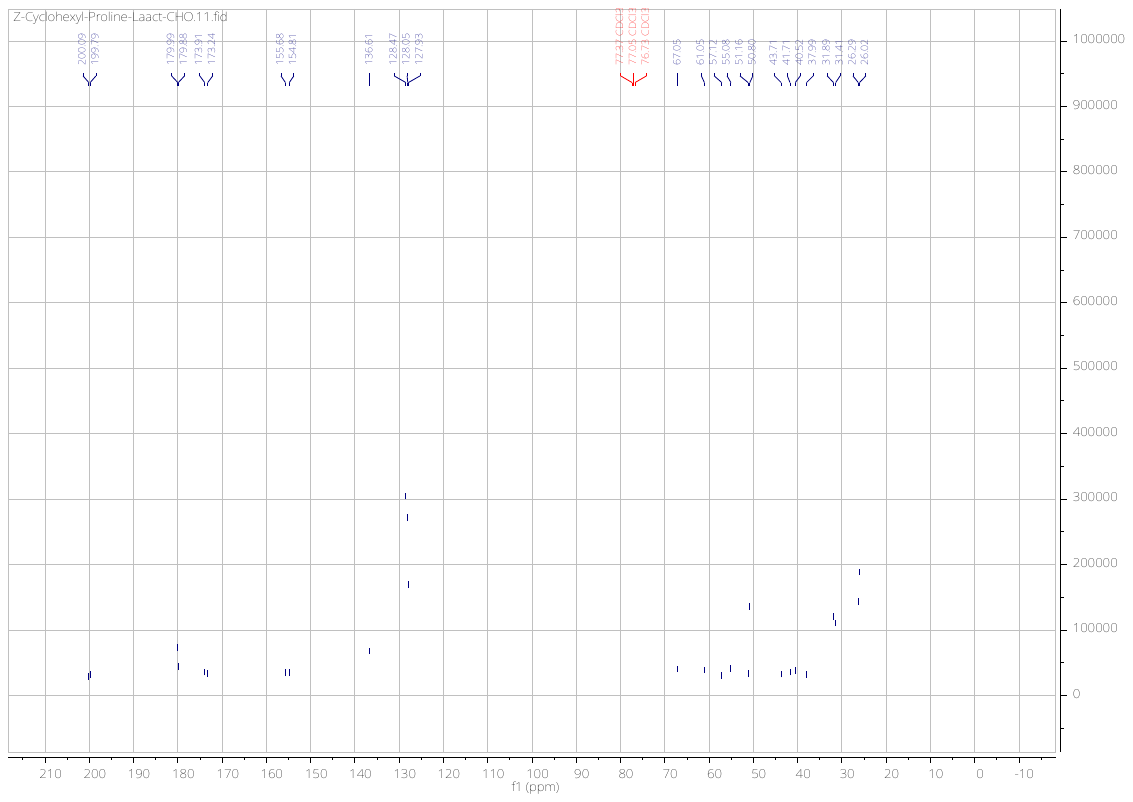

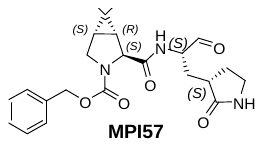

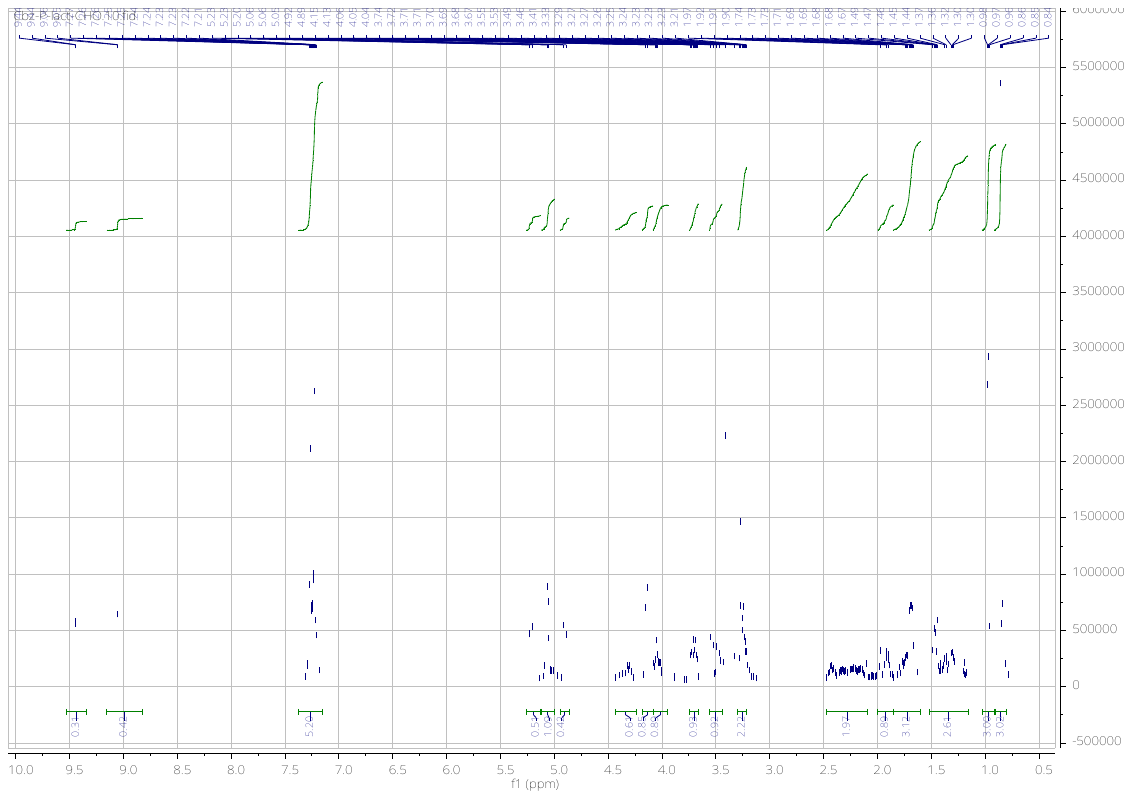

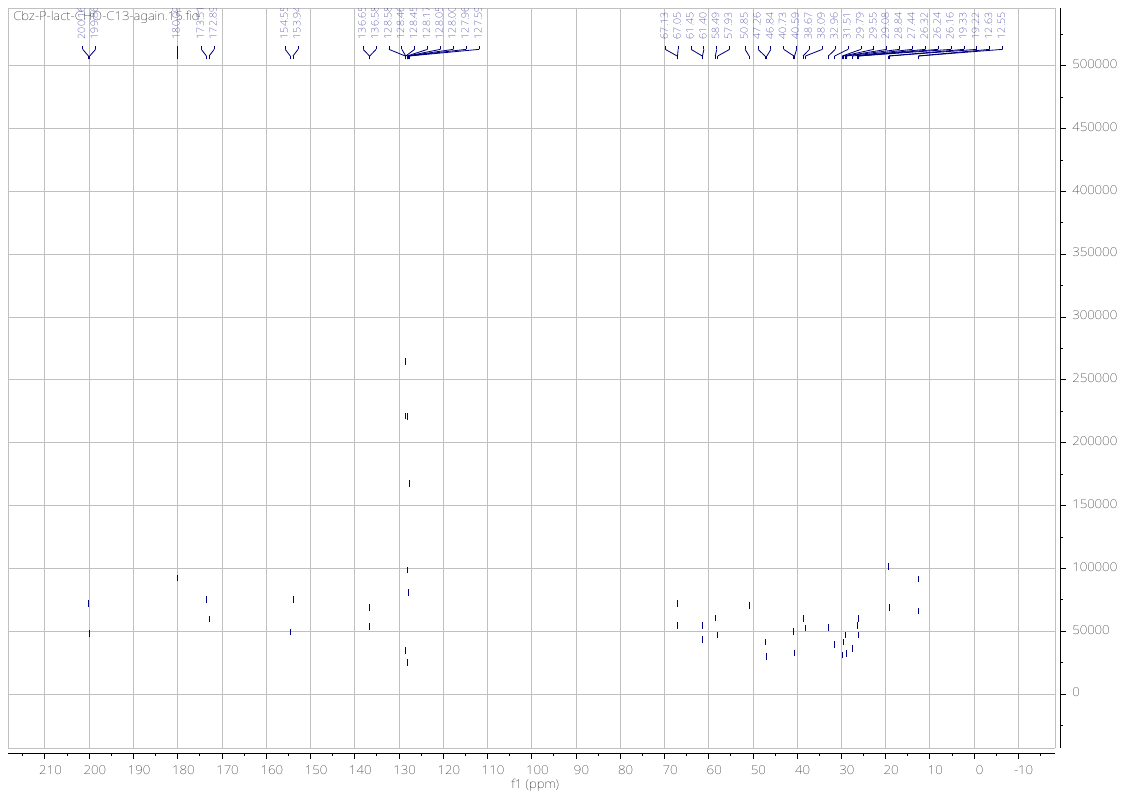

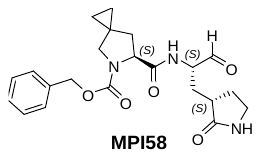

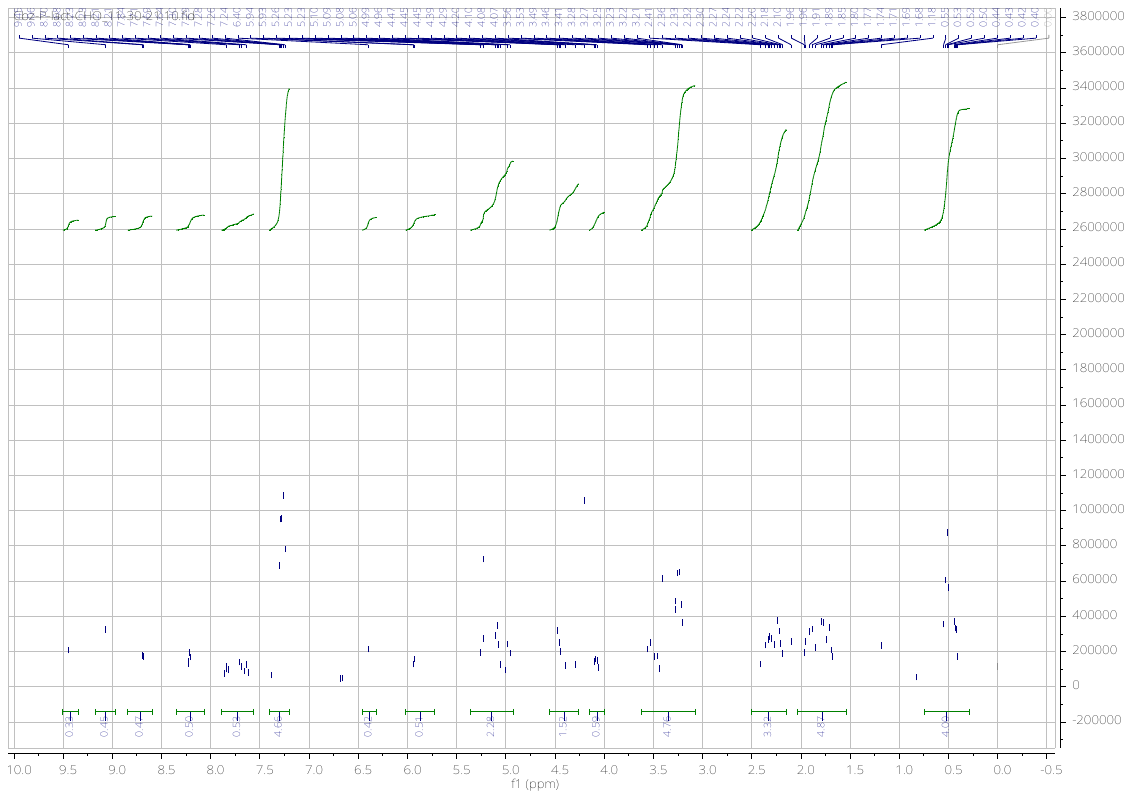

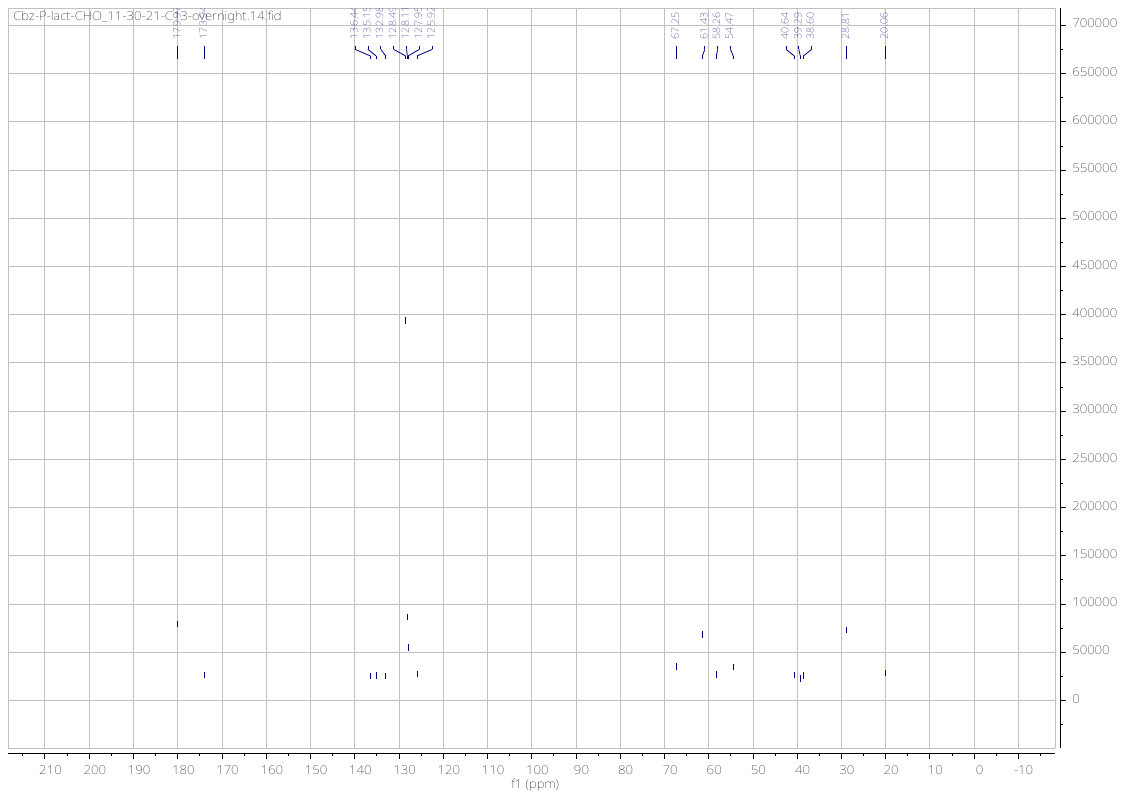

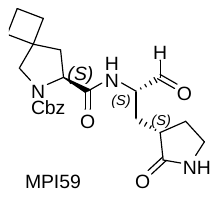

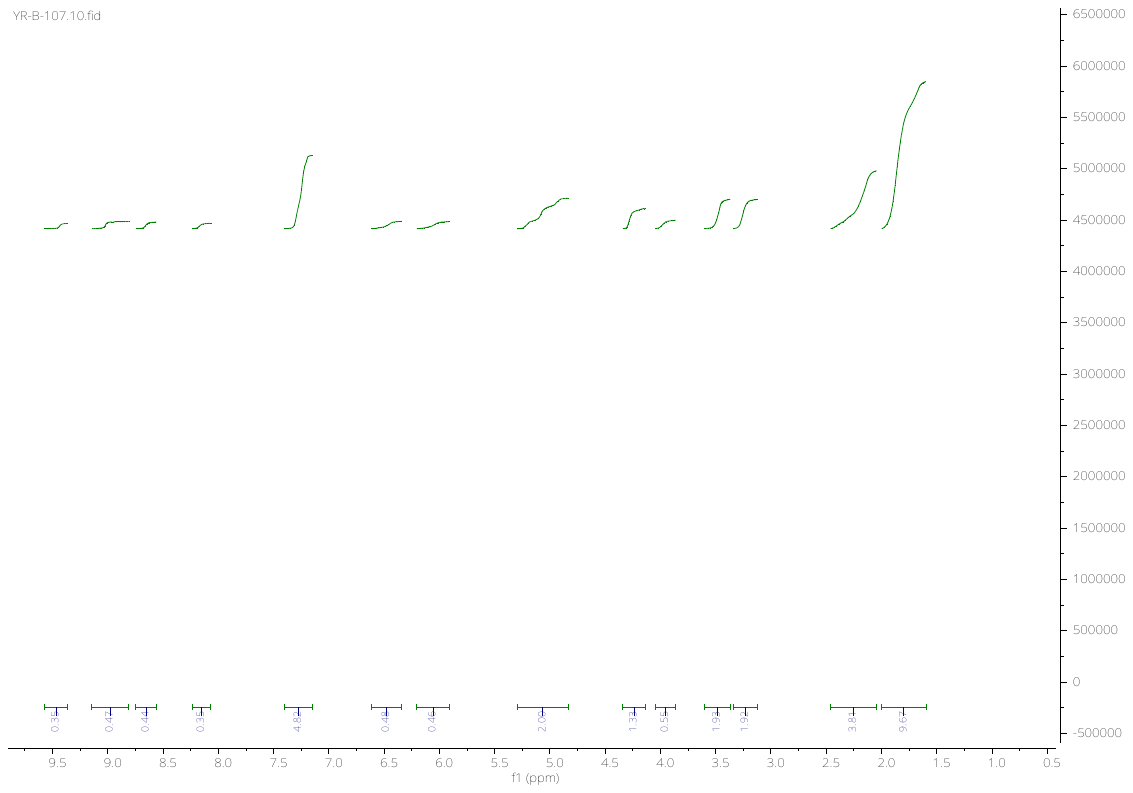

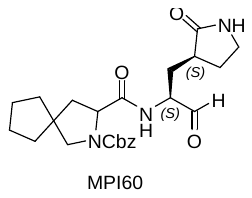

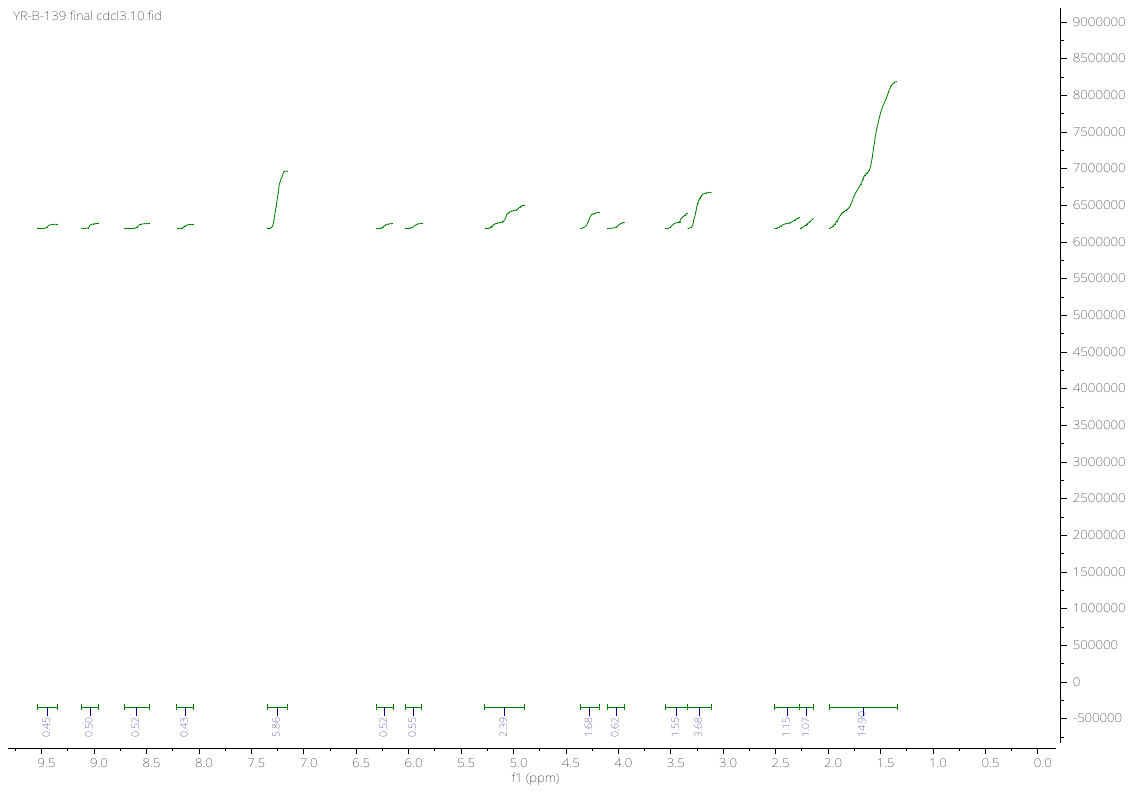

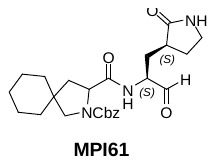

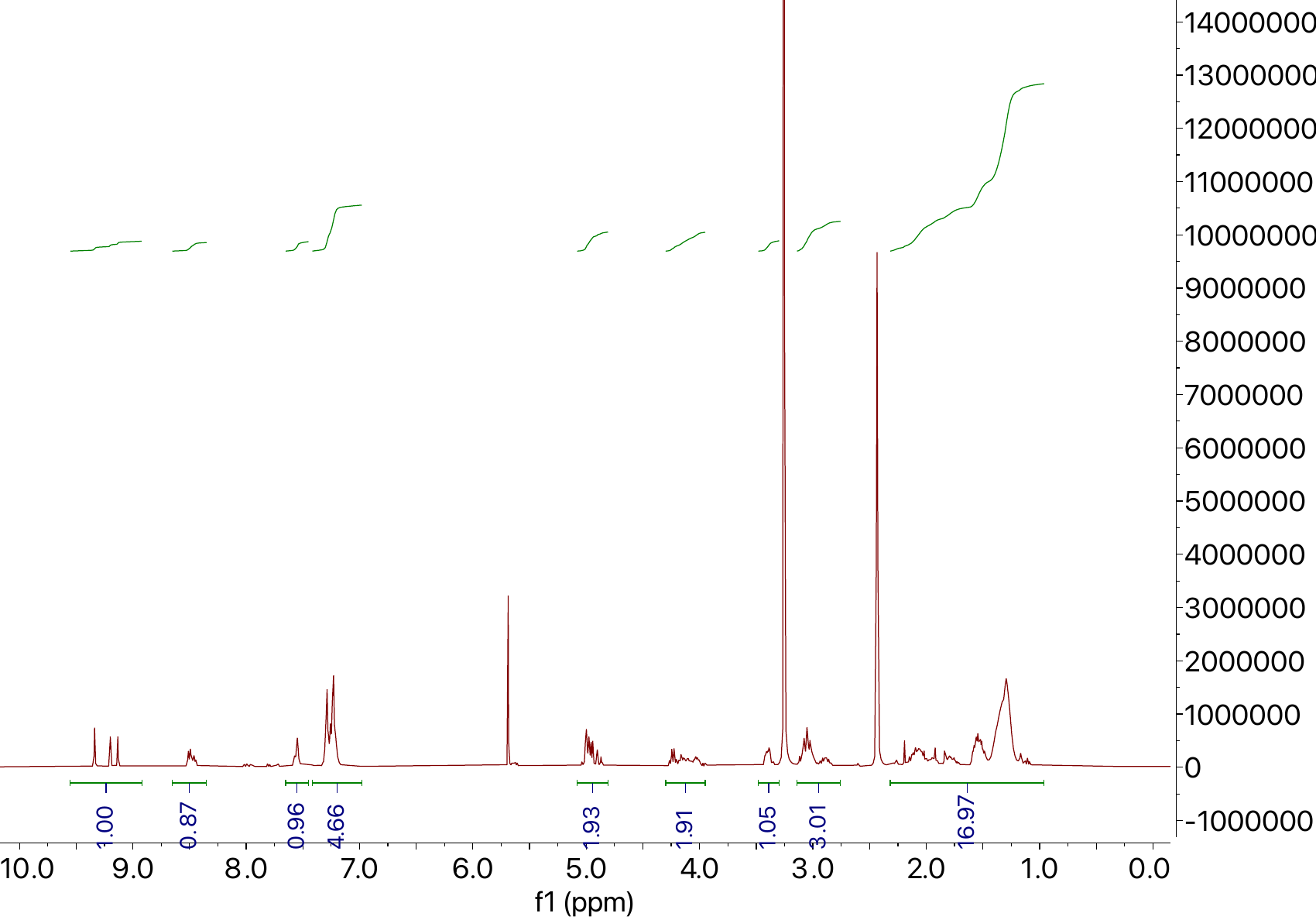

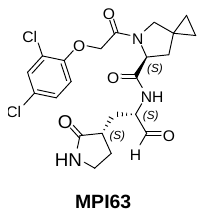

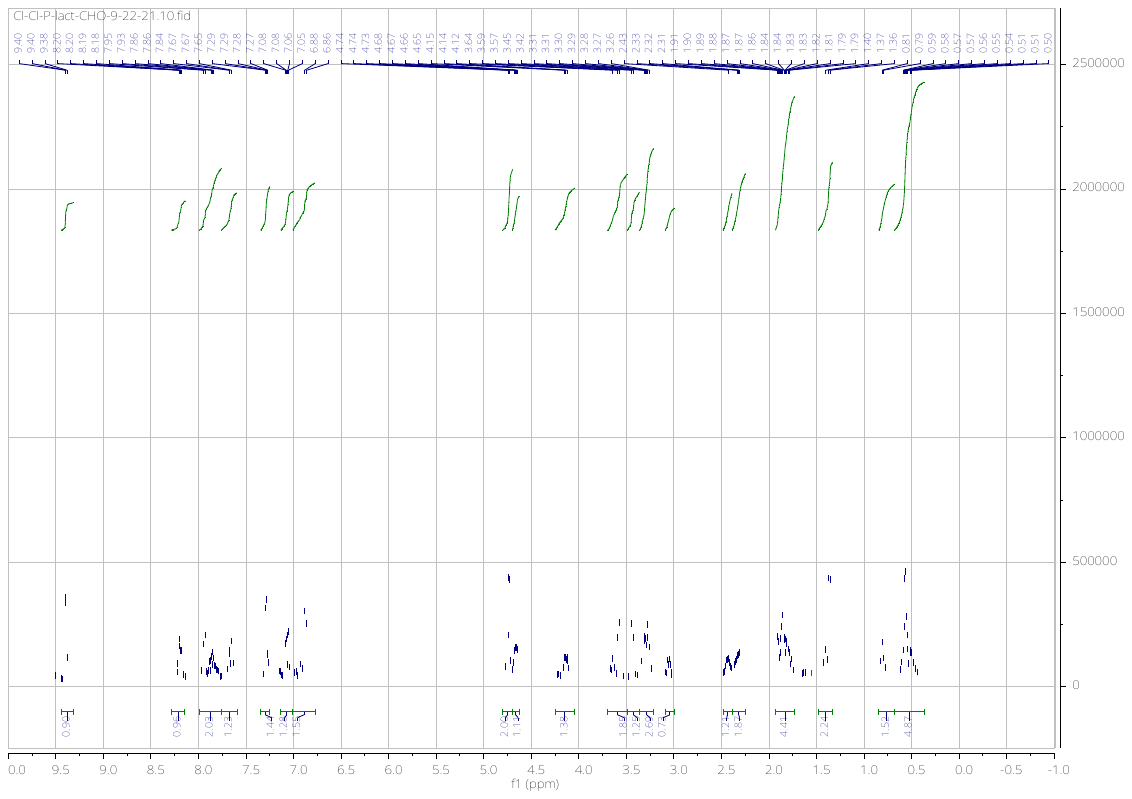

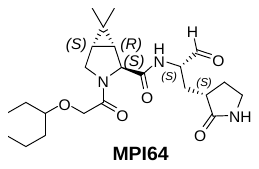

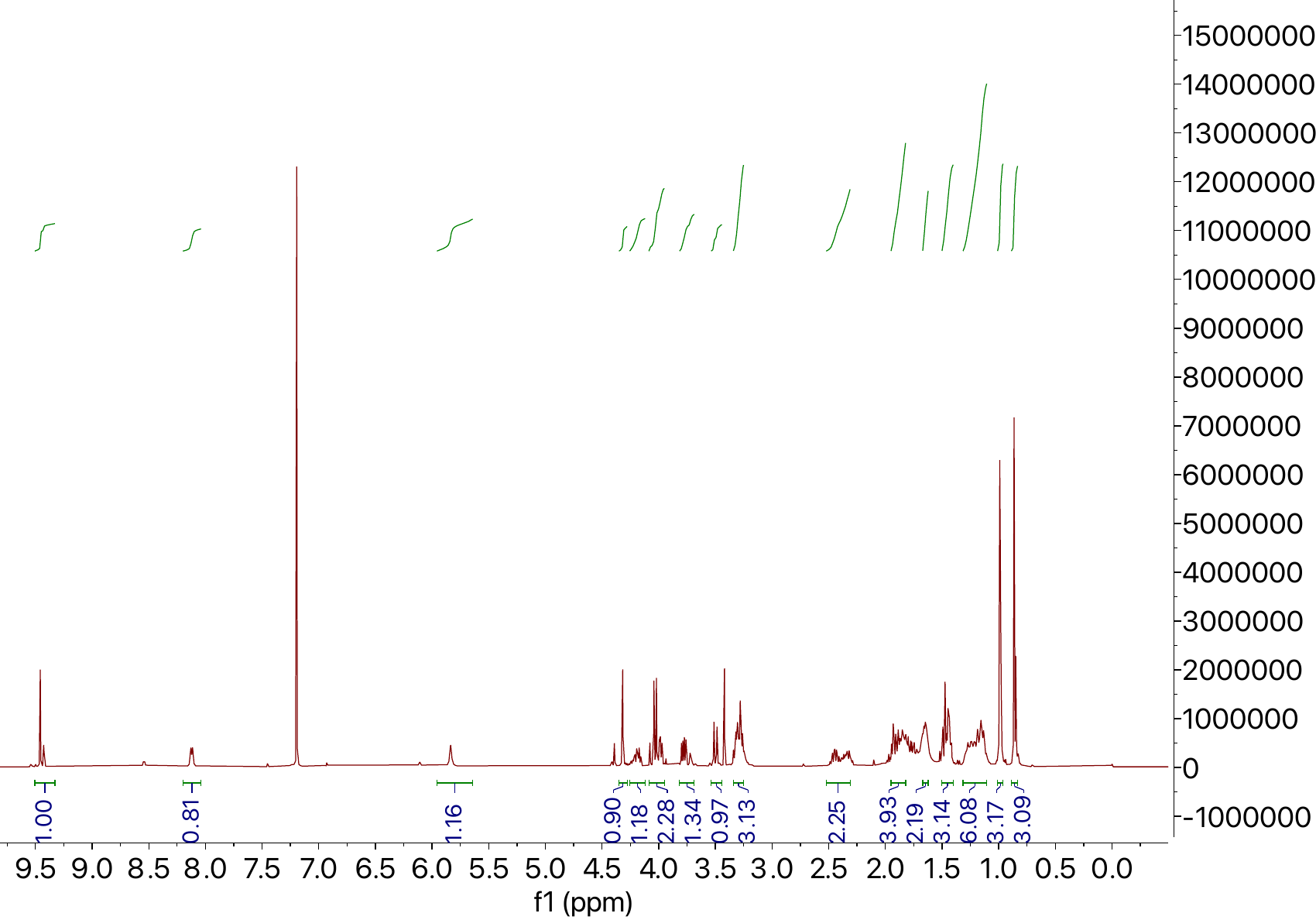

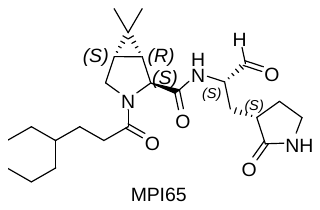

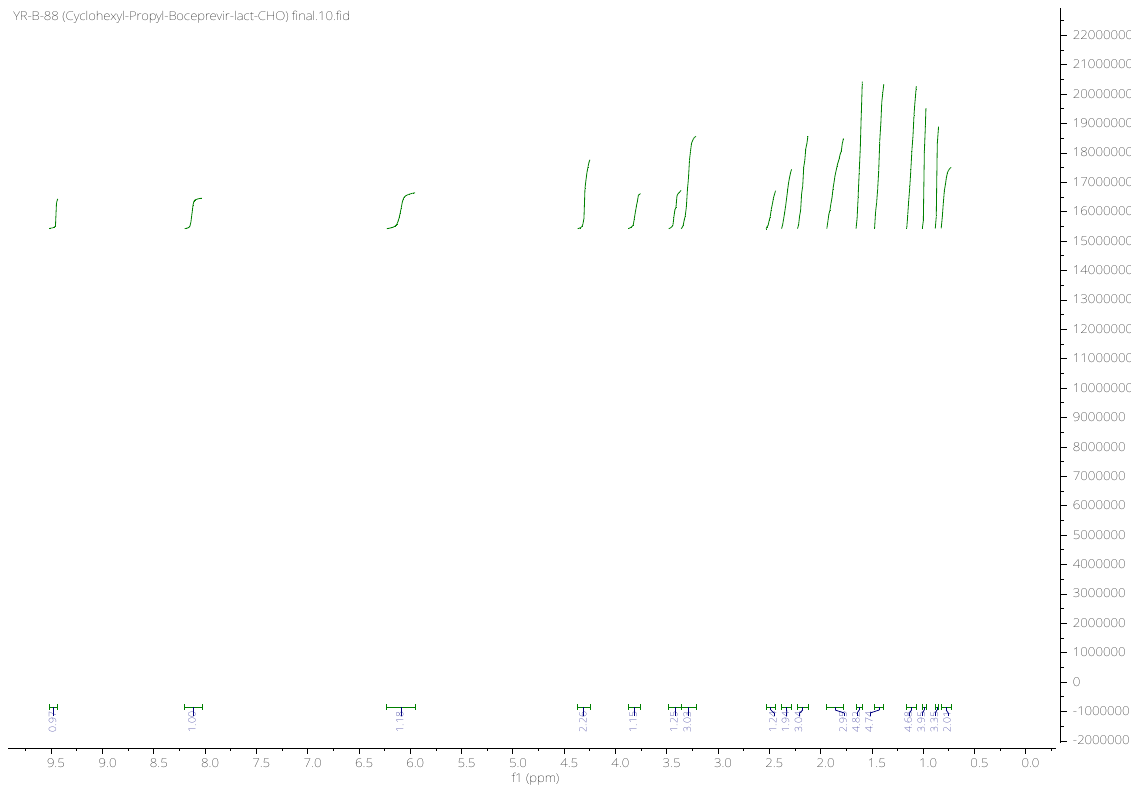

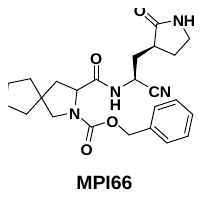

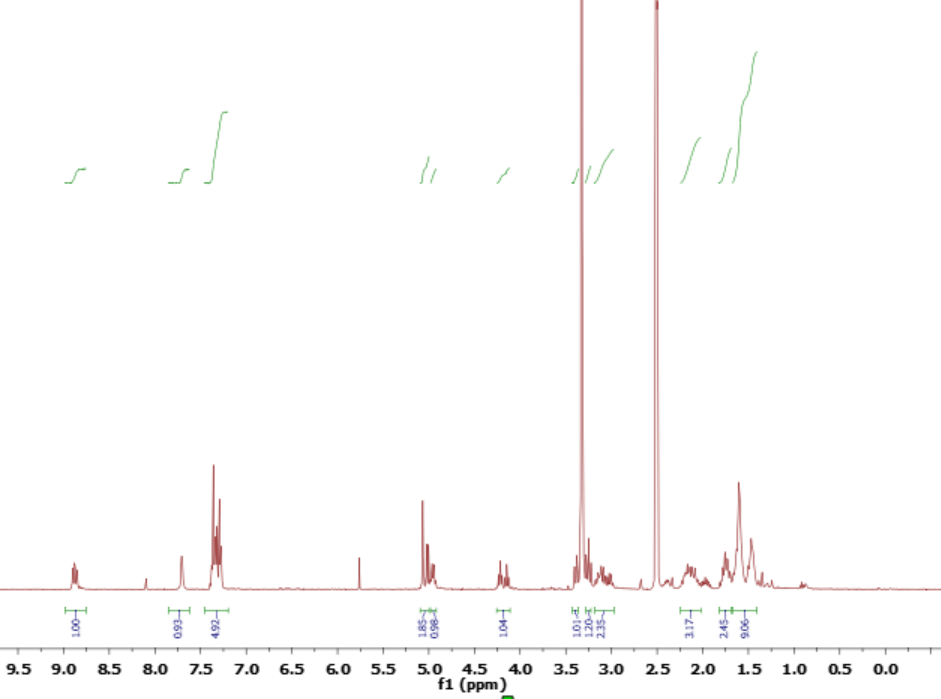

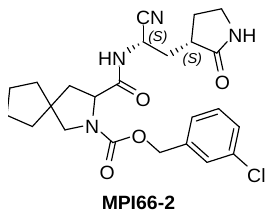

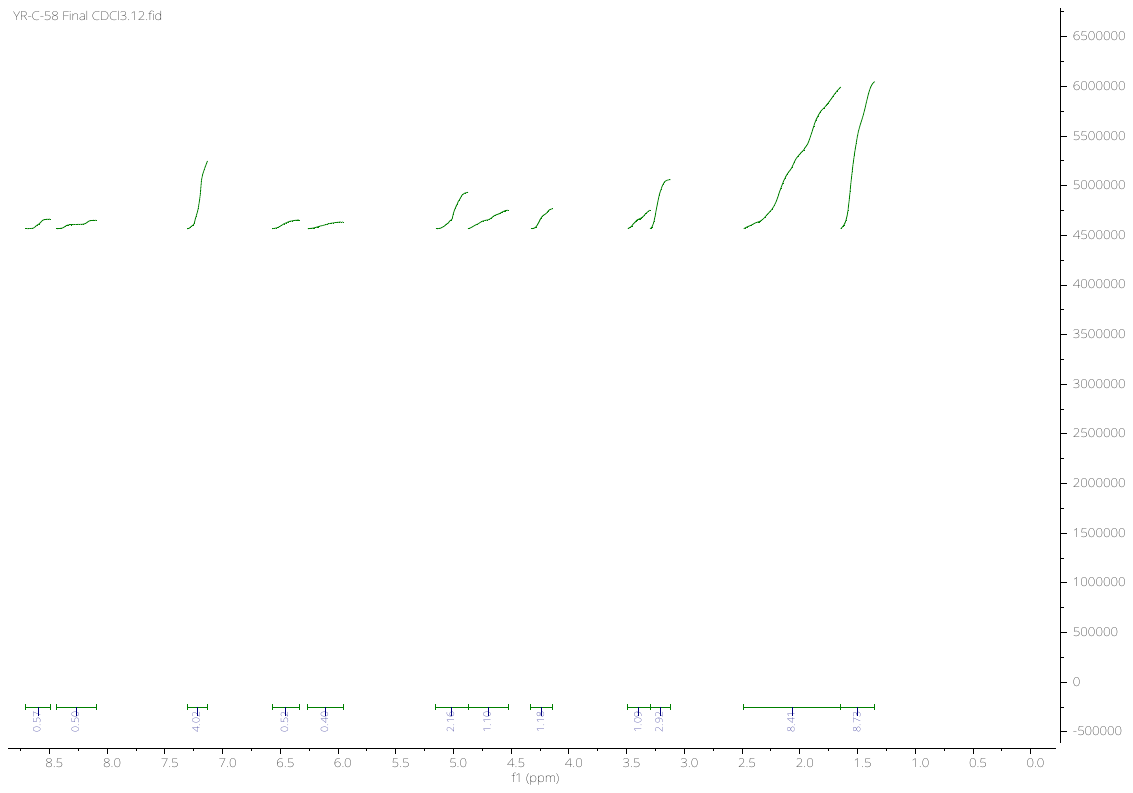

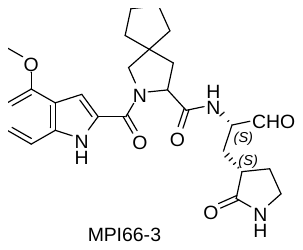

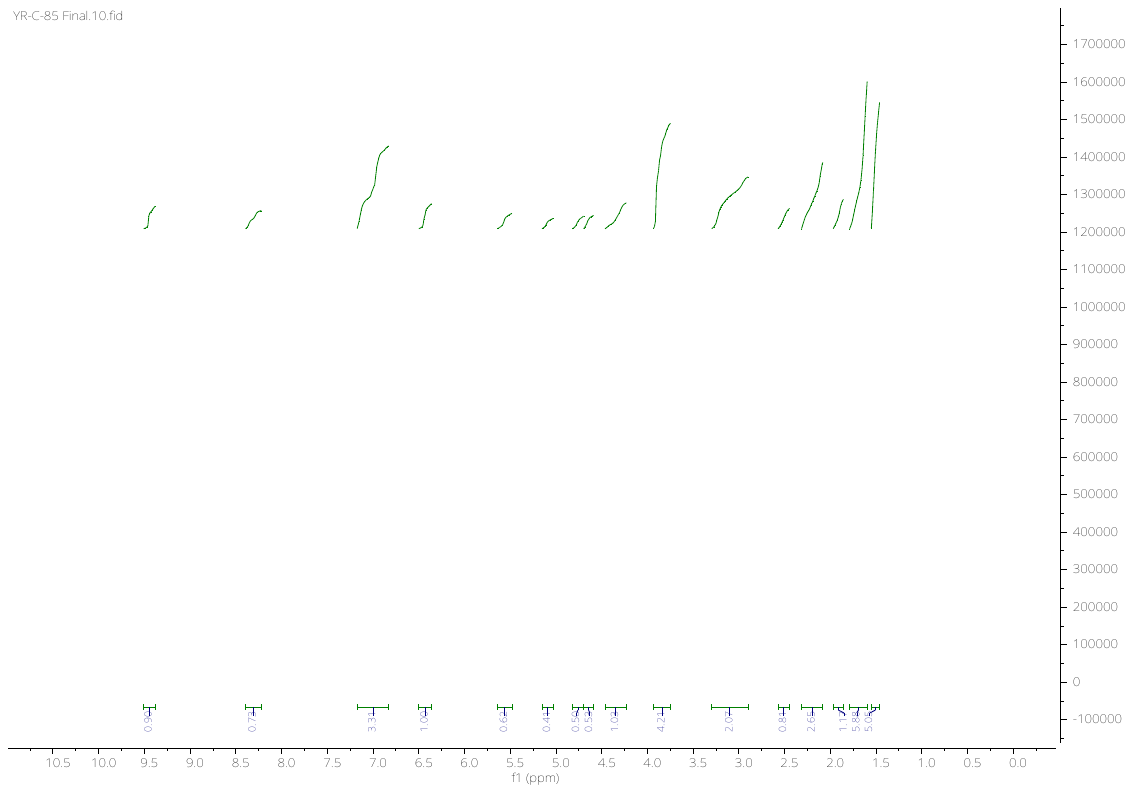

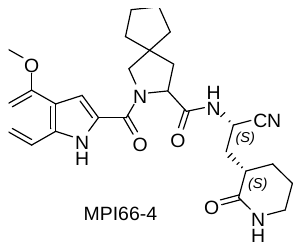

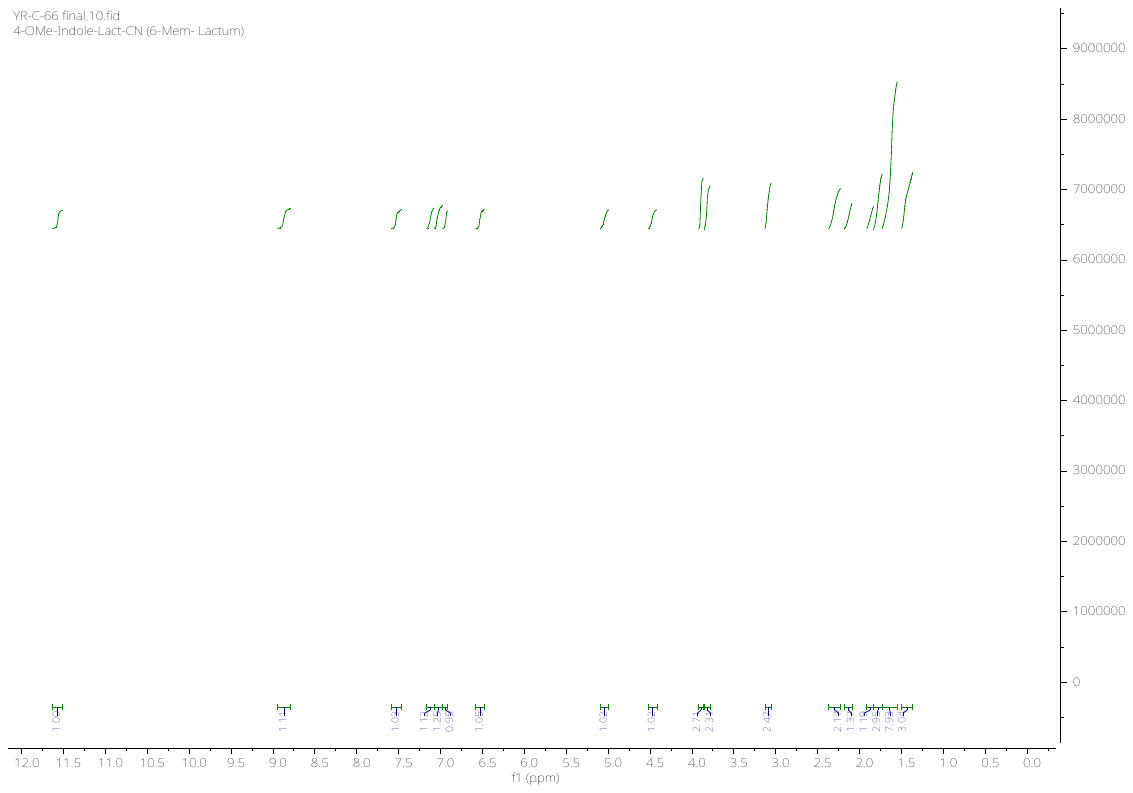

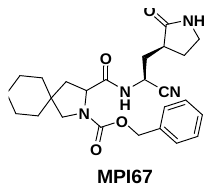

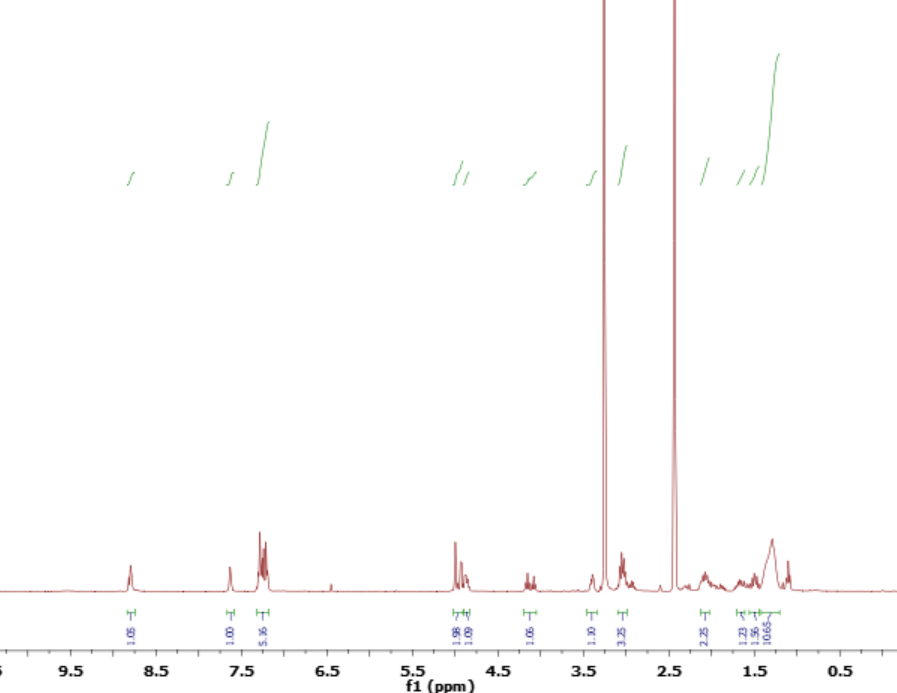

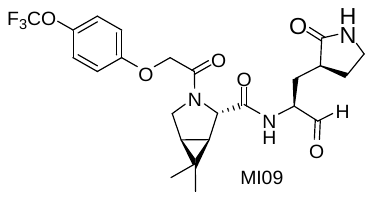

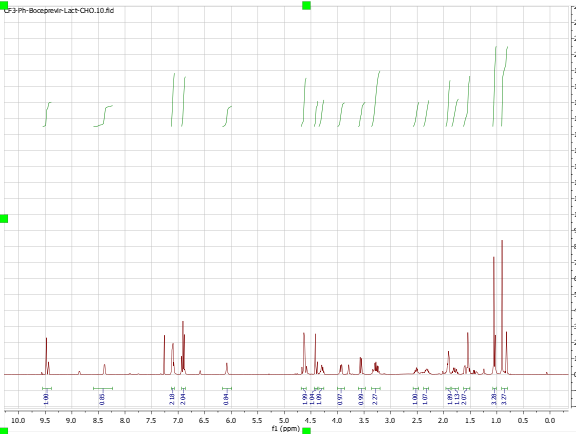

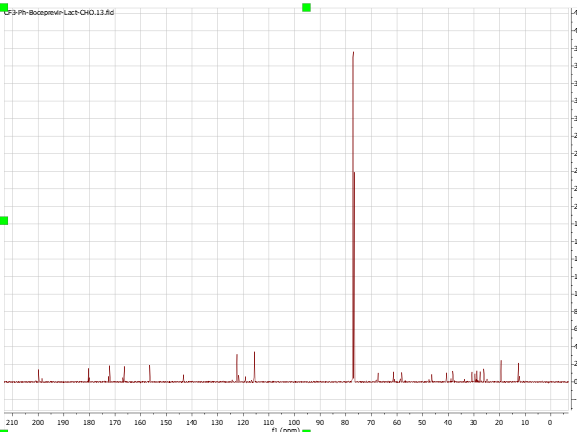

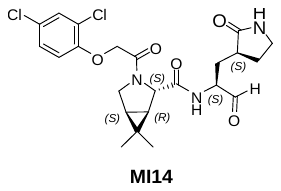

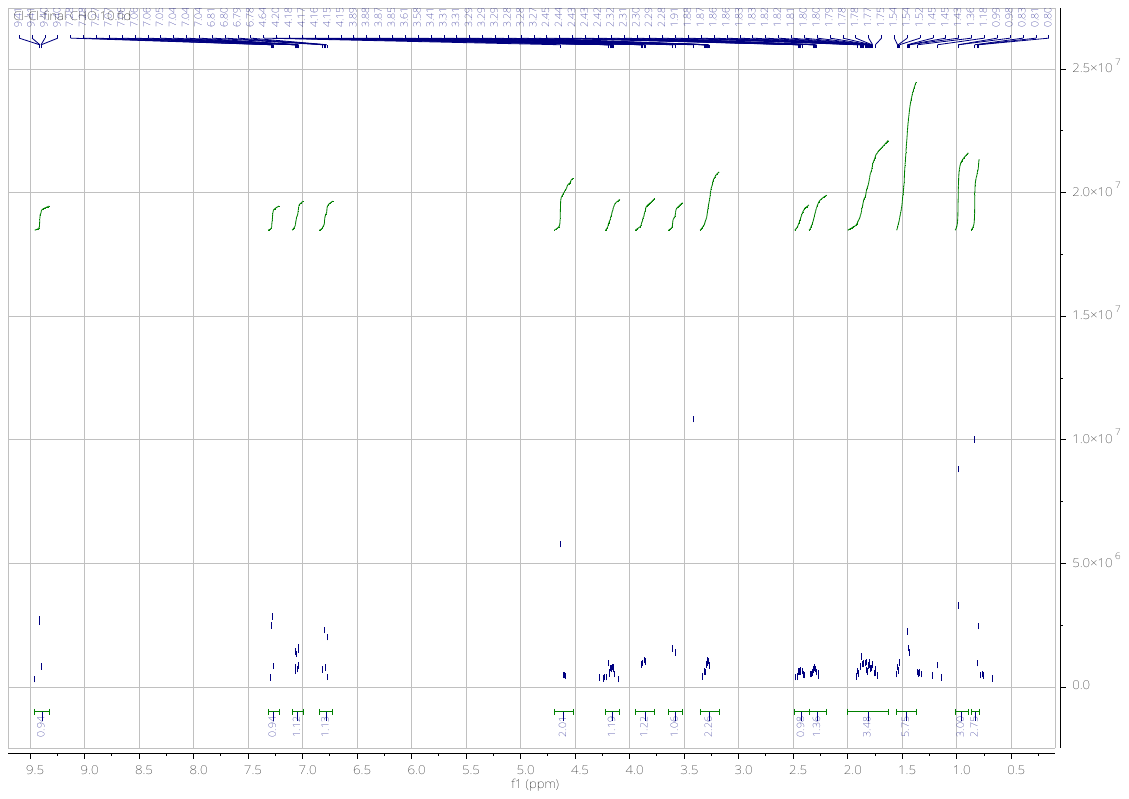

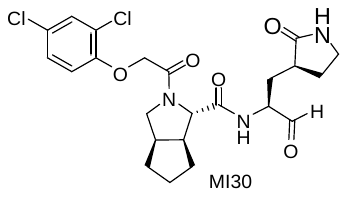

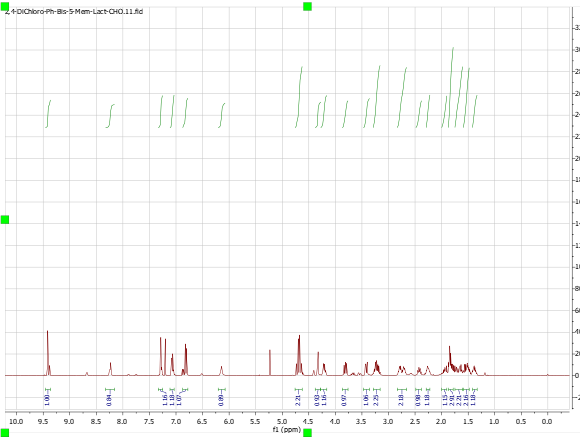

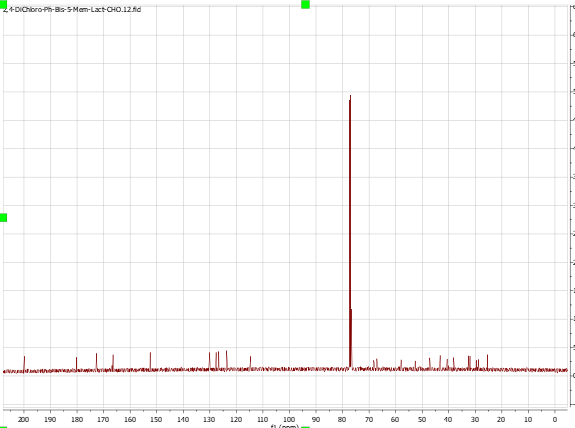

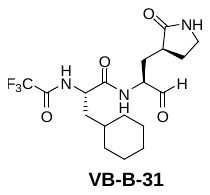

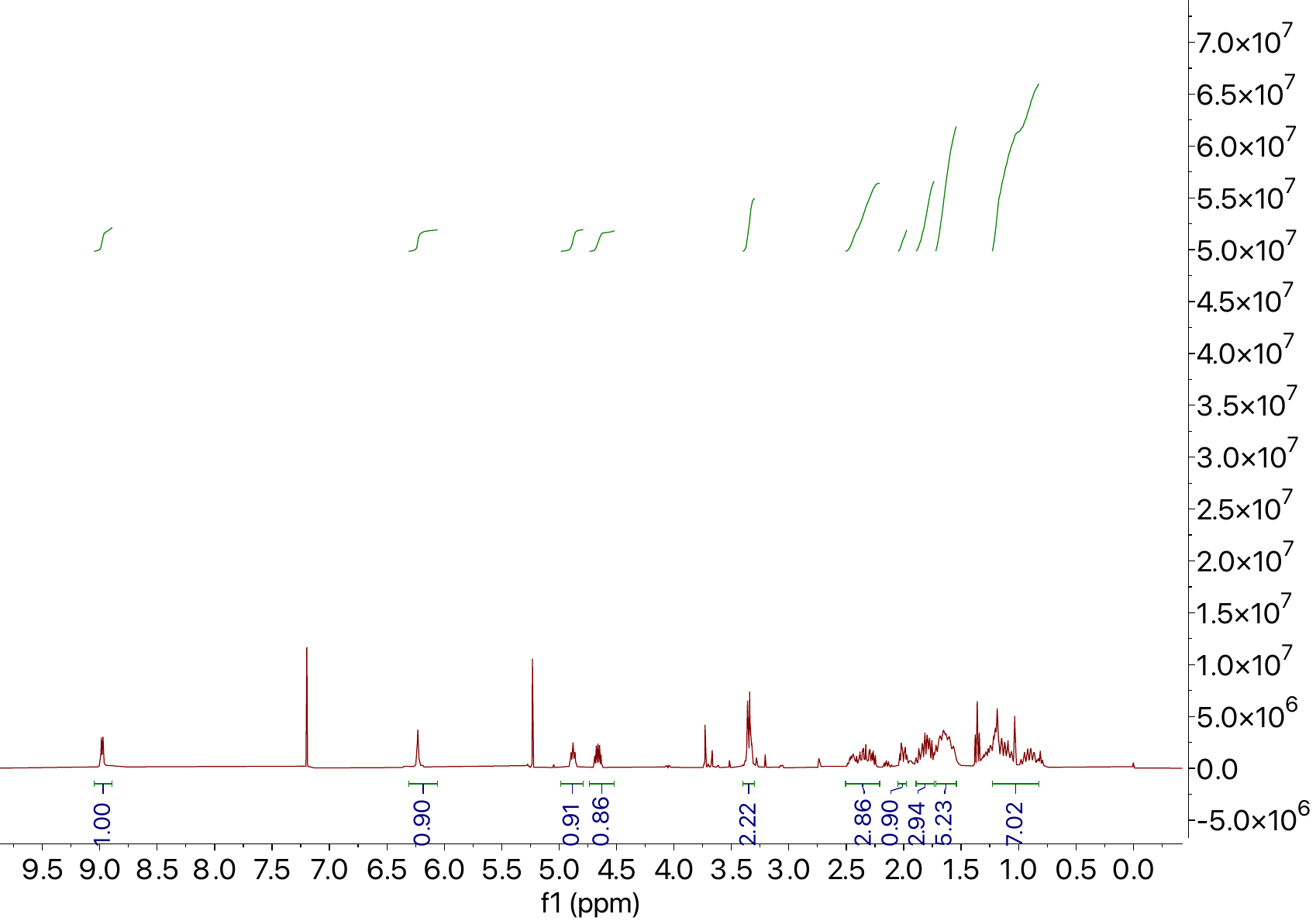

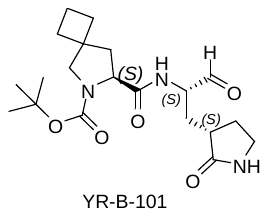

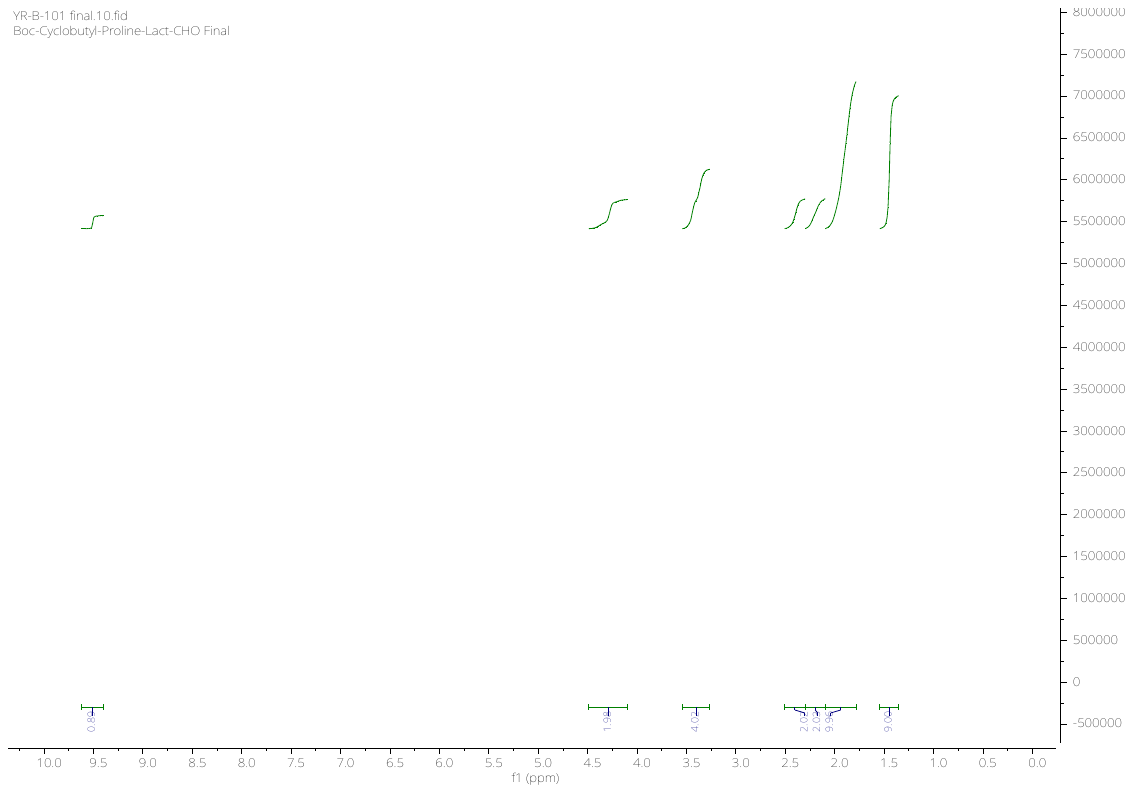
